# Indoline CD4-mimetic Compounds Mediate Potent and Broad HIV-1 Inhibition and Sensitization to Antibody-dependent Cellular Cytotoxicity

**DOI:** 10.1101/2023.01.02.522483

**Authors:** Christopher Fritschi, Saumya Anang, Zhen Gong, Mohammadjavad Mohammadi, Jonathan Richard, Catherine Bourassa, Kenny T. Severino, Hannah Richter, Derek Yang, Hung-Ching Chen, Ta-Jung Chiu, Michael Seaman, Navid Madani, Cameron Abrams, Andrés Finzi, Wayne A. Hendrickson, Joseph Sodroski, Amos B. Smith

**Author notes:** These authors contributed equally to this work (Christopher Fritschi, Saumya Anang, Zhen Gong). **Author Contributions**: C.F., S.A., Z.G., W.A.H., J.S., and A.B.S. designed research; C.F., S.A., Z.G., M.M., J.R., C.B., K.T.S., H.R., D.Y., H.C.C. and T.J.C. performed research; C.F., S.A., Z.G., M.M., J.R., C.B., K.T.S., H.R., D.Y., H.C.C., T.J.C., M.S., N.M., C.A., A.F., W.A.H., J.S. and A.B.S. analyzed data; and C.F., S.A., Z.G., W.A.H., J.S., and A.B.S. wrote the paper.

## Abstract

Binding to the host cell receptors, CD4 and CCR5/CXCR4, triggers large-scale conformational changes in the human immunodeficiency virus (HIV-1) envelope glycoprotein (Env) trimer [(gp120/gp41)_3_] that promote virus entry into the cell. CD4-mimetic compounds (CD4mcs) comprise small organic molecules that bind in the highly conserved CD4-binding site of gp120 and prematurely induce inactivating Env conformational changes, including shedding of gp120 from the Env trimer. By inducing more “open,” antibody-susceptible Env conformations, CD4mcs also sensitize HIV-1 virions to neutralization by antibodies and infected cells to antibody-dependent cellular cytotoxicity (ADCC). Here, we report the design, synthesis and evaluation of novel CD4mcs based on an indoline scaffold. Compared with our current lead indane scaffold CD4mc, BNM-III-170, several indoline CD4mcs exhibit increased potency and breadth against HIV-1 variants from different geographic clades. Viruses that were selected for resistance to the lead indane CD4mc, BNM-III-170, are susceptible to inhibition by the indoline CD4mcs. The indoline CD4mcs also potently sensitize HIV-1-infected cells to ADCC mediated by plasma from HIV-1-infected individuals. Crystal structures indicate that the indoline CD4mcs gain potency compared to the indane CD4mcs through more favorable π-π overlap from the indoline pose and by making favorable contacts with the vestibule of the CD4-binding pocket on gp120. The rational design of indoline CD4mcs thus holds promise for further improvements in antiviral activity, potentially contributing to efforts to treat and prevent HIV-1 infection.

## Introduction

The human immunodeficiency virus (HIV-1) establishes persistent infections that, if untreated, lead to the life-threatening acquired immunodeficiency syndrome (AIDS). The HIV-1 pandemic represents a significant challenge to global health, with 38 million people currently infected and 1.5 million new infections occurring annually (1). Antiretroviral treatments have extended the lives of infected individuals, but treatment needs to be continued indefinitely to prevent viral rebound in most cases (2). Furthermore, development of drug-resistant HIV-1 strains and drug side effects can limit effective treatment options (3). To date, practical measures to prevent HIV-1 transmission by antiretroviral drugs or vaccines remain elusive (2, 3). Additional approaches to limit HIV-1 replication would complement ongoing efforts to address these challenges.

Entry of HIV-1 into host cells is mediated by the envelope glycoprotein (Env) trimer, which consists of three gp120 exterior subunits noncovalently associated with three gp41 transmembrane subunits (4–6). As the only virus-specific protein on the surface of virions and infected cells, Env serves as a target for host antibodies that neutralize viruses and kill infected cells through antibody-dependent cellular cytotoxicity (ADCC) (7–11). The unliganded Env largely resides in a “closed” pretriggered conformation that resists the binding of potentially neutralizing or ADCC-mediating antibodies elicited during natural infection (12–16). CD4 binding drives Env into more “open” conformations that engage the CCR5/CXCR4 co-receptor, promoting additional Env transitions into a gp41 six-helix bundle that fuses the viral and cell membranes (17–22).

The binding site for CD4 is a conserved Env gp120 structure that is conformationally altered by CD4 binding (23, 24). In turn, CD4 binding creates an internal cavity (the Phe 43 cavity) in Env bounded by well conserved residues from gp120 and a single phenylalanine residue (Phe 43) from CD4 (*Figure 1A and B*). CD4-mimetic compounds (CD4mcs) comprise organic small-molecule HIV-1 entry inhibitors that bind in the Phe 43 cavity, as well as occupy the gp120 “vestibule” leading into the cavity (25). Highly conserved gp120 residues in the vestibule make critical contacts with CD4; therefore, by occupying the gp120 vestibule, the CD4mcs directly compete with CD4 (26, 27). In addition, CD4mcs prematurely trigger conformational changes in Env similar to those induced by CD4 (26, 28, 29). In the absence of coreceptor-expressing target cells, these prematurely activated Envs rapidly and irreversibly become non-functional, in some cases shedding gp120 (28, 29). At concentrations that do not completely inhibit HIV-1 infection, CD4mcs induce “open” conformations of Env, thereby sensitizing HIV-1 virions to neutralization or HIV-1-infected cells to ADCC by otherwise ineffectual antibodies (30–34). CD4mcs have been shown to synergize with ADCC-mediating antibodies to decrease the HIV-1-infected cell reservoir in humanized mice (35). CD4mcs directly protect humanized mice from vaginal acquisition of HIV-1 and, in combination with vaccine-elicited antibodies, CD4mcs protect monkeys from a stringent heterologous simian-human immunodeficiency virus (SHIV) mucosal challenge (36, 37).

**Figure 1.**
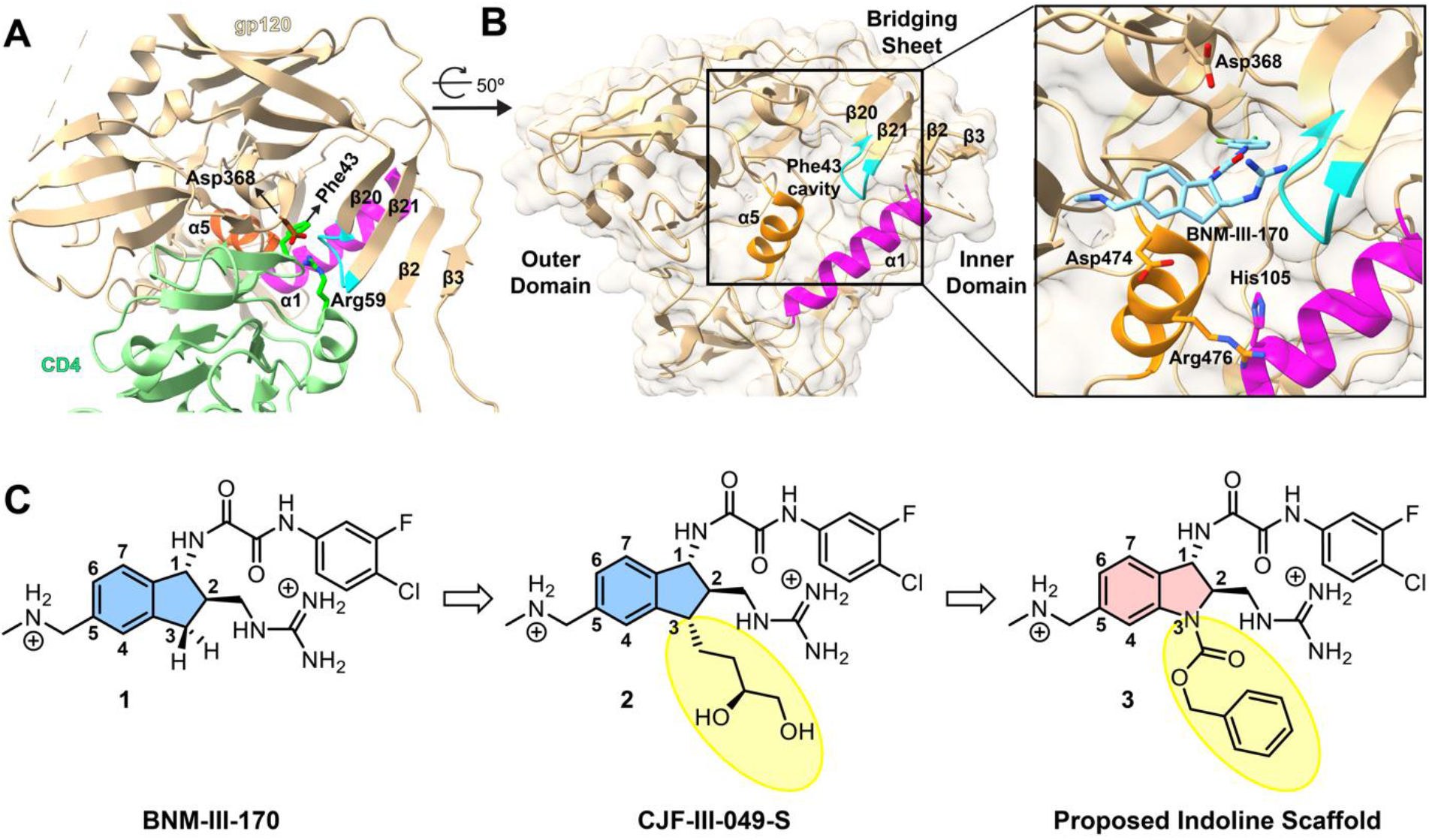
Targeting the Phe 43 cavity and surrounding vestibule of HIV-1 gp120. (**A**) Interactions between HIV-1_HXBc2_ gp120 and CD4 (PDB: 1GC1) are displayed with structural features crucial for CD4 binding highlighted. HIV-1_HXBc2_ gp120 and CD4 are shown in beige and light green ribbon diagrams respectively. The α5 (orange) and α1 (magenta) helices, as well as the β20/21 loop (cyan), harbor residues that contribute to native CD4 binding. Side chains of Asp368 of gp120 and Arg59 of CD4 are in stick representation to show interactions between gp120 and CD4. Residue Phe43 of CD4 is also depicted reaching into the heart of gp120. The gp120 carbon atoms, CD4 carbon atoms, and oxygen and nitrogen atoms are shown in brown, dark green, red and blue, respectively. **(B)** A protomer of HIV-1_BG505_ gp120 trimer (PDB: 5THR) is drawn as in **(A)** and oriented as indicated. The marked inset shows BNM-III-170, from a complex with HIV-1_C1086_ (PDB: 5F4P), in the vestibule of the Phe 43 cavity. Side chains are shown for gp120 residues targeted for CD4mc interaction as in structures with CJF-III-049-S (40) and with CD4 (23). (**C**) Evolution of more potent CD4mcs from BNM-III-170 (**1**) (38).

Early CD4mcs discovered by Debnath and co-workers (27) using a gp120-CD4 screen exhibited weak antiviral potency against a limited range of HIV-1 isolates. Replacement of the tetramethyl-piperidine scaffold of these prototypic CD4mcs with an indane scaffold represented a key advance (24, 37). Rationally designed indane CD4mcs (*Figure 1C*) were able to achieve more contacts with the gp120 vestibule, resulting in increased antiviral potency and breadth (38). Nonetheless, some primary HIV-1 strains remain resistant even to the most potent indane CD4mc, BNM-III-170 (**1**). HIV-1 passaged in the presence of BNM-III-170 develops resistance to this CD4mc as a result of Env changes in the Phe 43 cavity and in a gp120 element that allosterically modulates CD4 binding (39).

The practical utility of CD4mcs as antiviral agents would be enhanced by the availability of more potent analogs with better coverage of HIV-1 strains. Guided by crystal structures of BNM-III-170 and congeners thereof, we found that 3-substituents on the indane ring of BNM-III-170 can contact a vestibule pocket situated at the interface of the gp120 inner and outer domains, between the α1 and α5 helices (*Figure 1B*). As a result, some 3-substituted indane analogs, such as CJF-III-049-S (**2**, *Figure 1C*), exhibited modest (3-fold) improvements in antiviral activity compared with BNM-III-170; however, only the viruses with His 105 in the gp120 α1 helix showed such an increase (40).

Based on the above observations, we hypothesized that replacing the C3 carbon with a nitrogen (*Figure 1C*) to create an indoline scaffold might have several benefits. First, the N3 nitrogen of the indoline scaffold would permit facile incorporation of diverse functional groups (*vide infra*). Second, the nitrogen at N3 would be expected to limit conformational freedom of the 5-membered ring, thereby decreasing the number of conformational states that the oxalamide (at C1) and methyl guanidinium (at C2) can adopt. Only one state has been observed by X-ray for the indane CD4mc complexes with gp120 (24, 37, 39), but an entropic penalty would be expected in selecting this pose upon complexation. Lastly, the rigidified 5-membered ring of the indolines may lead to different geometries of the substituents at the N3 position, compared to the reported indane congeners (40). This would permit different branching angles of the substituents, thereby allowing exploration of broader chemical space, given the highly modular nature of the indoline scaffold.

## Results

### A. Synthetic Chemistry

Based on the rationale above, we set out to design a synthetic route to obtain the *indoline* CD4mc framework. As illustrated in *Figure 2A*, anthranilic ester **4** was first protected as a benzyl carbamate. This was followed by one-pot alkylation with methyl bromoacetate (**5**), and then an intramolecular Claisen condensation employing cesium carbonate to furnish the oxindole **6**.

**Figure 2.**
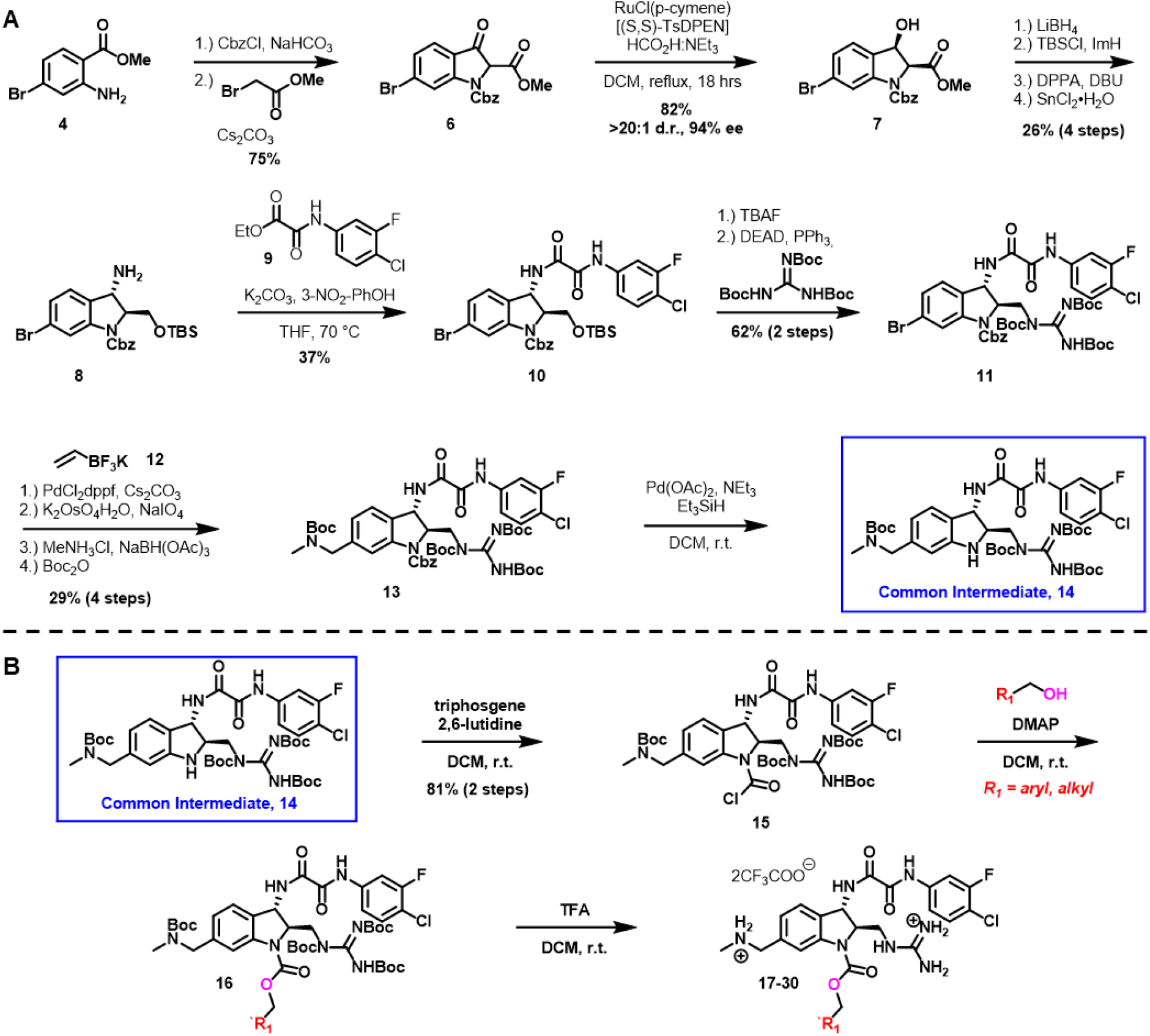
Synthesis of indoline core CD4mcs. (**A**) Synthetic route towards a common intermediate for analog synthesis starting from commercially available methyl 2-amino-4-bromobenzoate **4**. This route affords the indoline scaffold-containing intermediate **14** in 15-steps and 1.1% overall yield. (**B**) Functionalization strategy of the common intermediate **14** for analog synthesis.

Judicious choice of the aniline nitrogen protecting group proved important for late-stage functionalization, as we desired to react selectively at this N3 nitrogen in the presence of both the scaffold’s guanidinium (at C2) and methylamino methyl (at C5) substituents. Taking inspiration from our process synthesis developed for BNM-III-170 (41), a dynamic kinetic resolution was performed on the oxindole **6** to give the β-hydroxy ester **7** (42). From **7**, it was necessary to install an amine at the benzylic position of the indoline core. To accomplish this, we first reduced the ester with LiBH_4_, followed by protection of the primary alcohol.

Stereoinversion was then accomplished via displacement of the secondary alcohol using diphenylphosphorylazide (DPPA) to furnish an azide, that was next reduced with tin (II) chloride to give the desired primary amine **8**. Installation of our previously optimized oxalamide (43) utitlizing 9 on the derived primary amine gave intermediate **10**, which was subjected to tetrabutylammonium fluoride (TBAF) to effect removal of the silyl protecting group, followed by Mitsunobu union of the resulting primary alcohol with tri-N-bocguanidine to yield intermediate **11**. With these two crucial structural features installed, we turned towards installation of the methylamino methyl side-chain. Suzuki coupling (44) of aryl bromide **11** with potassium vinyltrifluoroborate (**12**), oxidative cleavage, followed by reductive amination of the resulting aldehyde with methylamine, and protection of the free amine as the *tert*-butyl carbamate provided intermediate **13**.

Initially, removal of the benzyl carbamate proved difficult, as employing standard hydrogenation conditions resulted in dechlorination of the *m*-F-*p*-Cl aromatic ring. Instead, transfer hydrogenation utilizing Pd(OAc)_2_ and Et_3_SiH as the hydride source permitted selective removal of the benzyl carbamate protecting group to give **14**. It should be noted that the indoline scaffold is very sensitive to acidic conditions. Significant transfer of the *tert*-butyloxycarbonyl from the adjacent protected guanidine to the indoline nitrogen was observed even when subjected to weak acids, such as pyridinium hydrochloride. With release of the indoline nitrogen having been achieved, acylation with triphosgene and 2,6-lutidine as base yielded carbamoyl chloride **15**, which proved to be stable, and as such could be used directly without purification in conjunction with various alcohols to produce carbamates such as **16** in excellent yield. Finally, removal of the protecting groups resulted in the carbamates as TFA salts (**17**–**30**, *Figure 2B* and *Figure 3*).

**Figure 3.**
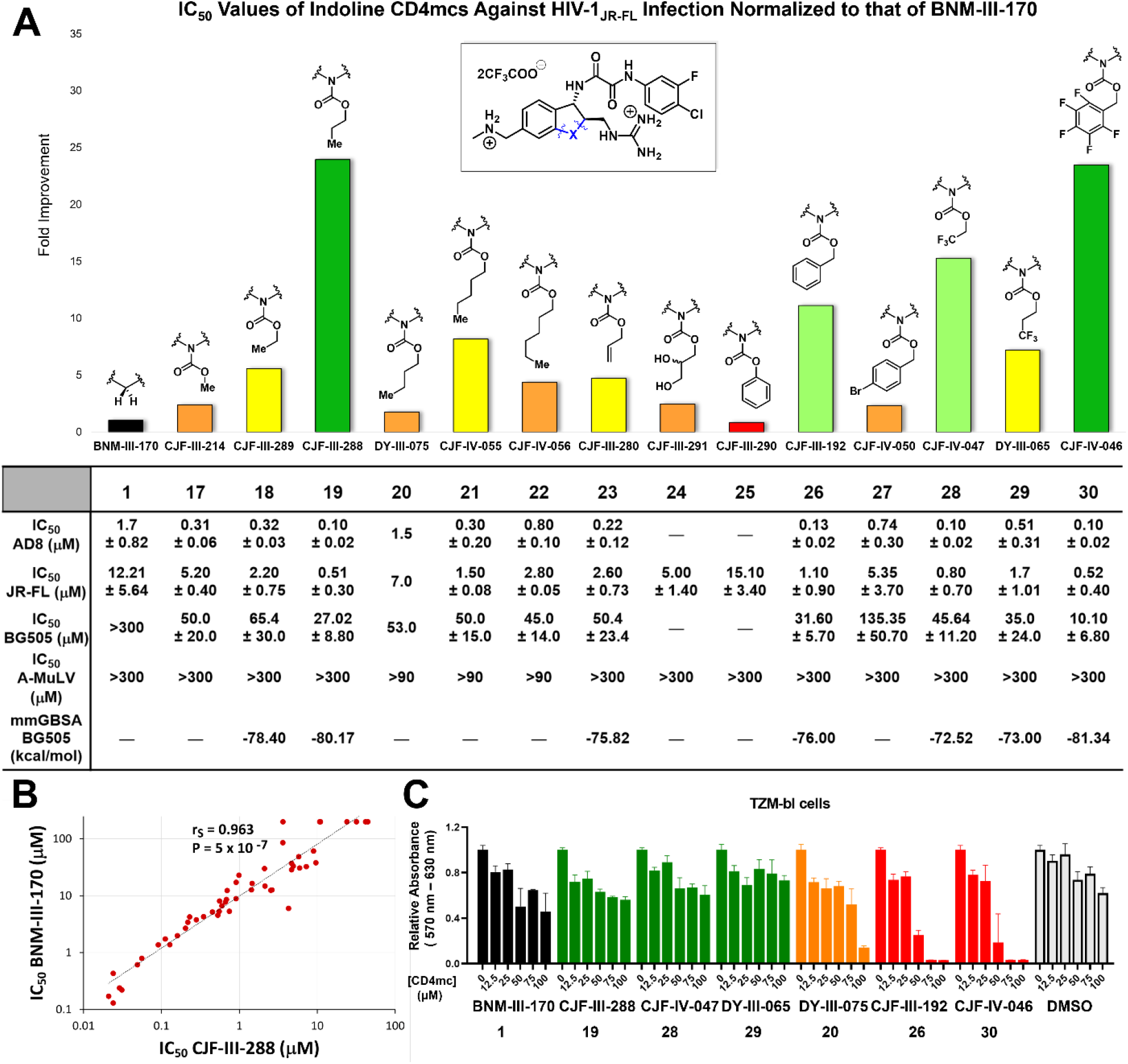
Indoline CD4mcs inhibit HIV-1 entry more effectively than the previous lead indane CD4mc BNM-III-170. (**A**) Inhibition of infectivity of recombinant HIV-1 pseudotypes by CD4mc analogs. The bar graph shows the inhibitory activity of indoline CD4mcs against infection of recombinant virus pseudotyped with the Env of HIV-1_JR-FL_, normalized to that of the IC_50_ of BNM-III-170. The table beneath reports the IC_50_ values of indoline CD4mcs against viruses pseudotyped with the Envs of the AD8, JR-FL and BG505 HIV-1 strains. Only three of the compounds (DY-III-075, CJF-IV-055 and CJF-IV-056) inhibited the control virus pseudotyped by the amphotropic murine leukemia virus (A-MLV) Env at concentrations of 300 μM or less. The table also reports the free energy of binding for the indoline CD4mc analogs predicted by an HIV-1_BG505_ gp120 docking model. (**B**) Comparison of the inhibition of infectivity by BNM-III-170 (38) and CJF-III-288 against a panel of recombinant viruses pseudotyped with the Envs of HIV-1 strains from multiple phylogenetic clades. Compared with BNM-III-170, CJF-III-288 displays an approximately 10-fold increase in potency for all HIV-1 pseudotypes examined, with the exception of clade A/E recombinants. Neither BNM-III-170 nor CJF-III-288 inhibited the control virus pseudotyped by the amphotropic murine leukemia virus (A-MLV) Env at concentrations of 100 μM or less. (**C**) The cell proliferation assay was conducted at varying concentrations of BNM-III-170 and indoline CD4mcs prepared from stock solutions in 100% DMSO. Compounds were incubated with TZM-bl cells for 72 h, after which the relative viability of the cells was evaluated with respect to the cells without treatment. The volume of DMSO used in the DMSO control corresponds to the volume of the CD4mc used at each indicated concentration. Measurements were performed in triplicate. Error bars indicate mean ± SEM. The levels of TZM-bl cell toxicity for the indoline CD4mcs are indicated as minimal (green), moderate (orange) or severe (red).

Synthetic modifications at the indoline N3 position proceeded with prioritization based on a combination of chemical feasibility, structural considerations, and feedback from viral inhibition assays. Molecular modeling of the CD4mc-gp120 complexes was informed by the disposition of the indane C3 substituent in CJF-III-049-S (**2**), similarity between the indoline and indane scaffolds, and the topography of the gp120 vestibule surface (*Figure S5 and S6)*. Crystal structures of gp120 complexes with selected compounds tested and refined hypotheses in this structure-based development course.

### B. Inhibition of HIV-1 Infection by Indoline CD4mcs

Initially, several carbamate analogs were synthesized and tested in an assay measuring the inhibition of single-round HIV-1_JR-FL_ infection. We were curious to know if the simplest carbamate (**17**, X = NCO_2_Me) would retain the ability to inhibit Env function. Gratifyingly, this analog displayed a modest 2.5-fold increase in potency (*Figure 3A*) compared to the previous lead CD4mc, BNM-III-170 (**1**). We also found that indoline **17** bound to gp120 in a similar manner as for indanes **1** and **2**. These results suggested that a wide series of indoline CD4mc warranted further investigation. Carbamates in the alkyl series of ethyl (**18**), propyl (**19**), butyl (**20**), pentyl (**21**), and hexyl (**22**) carbon chain replacements for the methyl group were readily synthesized and tested for inhibition of JR-FL virus pseudotypes (*Figure 3A*). As the carbon chain length grew, virus inhibition initially increased and then decreased, with peak antiviral activity 24-fold higher than for BNM-III-170 for the three-carbon chain [CJF-III-288 (**19**), *Figure 3A*].

We also explored the consequences of unsaturated bonds in the carbamate system to see if any effect of addition of an electron pi-cloud could be observed. Allyl carbamate **23** and benzyl carbamate **26** were therefore constructed and tested. The allyl analog **23** proved to be less potent than its propyl counterpart **19**, thus providing no indication of significant pi interactions in the targeted area (*Figure 3A*). However, the benzyl carbamate **26** demonstrated an 11-fold increase in potency relative to BNM-III-170 (*Figure 3A*), indicating that the proposed targeted area on Env is sufficient to accommodate an aromatic ring without issue. In general, hydrophobic substituents on the N3-carbamate appear to interact efficiently with the targeted Env region, leading to more potent viral inhibition.

To probe further the structural features of the CD4mc that could increase antiviral potency, we examined the effect of deletion of the one-carbon spacer between the phenyl ring and the carbamate oxygen in CJF-III-192 (**26**). To this end, the phenyl carbamate derivative **25** was constructed and tested. Interestingly, the antiviral potency of this CD4mc was decreased compared with that of the benzyl carbamate, CJF-III-192 (**26**) (*Figure 3A*). The crystal structure of CJF-III-192 (**26**) complexed to gp120 (*Figures S1F and S3C*) shows that the one-carbon spacer is essential for providing the rotatable bond needed for the phenyl ring to adopt a favorable binding conformation. Introduction of a *para*-bromine on the phenyl ring of **26** to yield **27** decreased the antiviral activity (*Figure 3A*); this decrease is not readily explained by the crystal structure of **26**, since the para position projects away from gp120 into the solvent.

Allyl carbamate **23** proved useful for analog synthesis as well. A 1,2-diol functionality was readily introduced from the alkene via dihydroxylation. The resulting diol **24** is structurally similar to the indane CD4mc CJF-III-049-S (*Figure 1C*), but the butanediol substituent is placed further out from the scaffold. A 2.5-fold improvement in viral inhibition by **24** (*Figure 3A*) compared to BNM-III-170 was observed. These results indicate that **24** is comparable to CJF-III-049-S (**2**) (*Figure 1C*) in antiviral activity but less active than **23**; therefore, the addition of a diol at this location did not apparently enhance positive interactions with Env.

Selective fluorine incorporation has proved useful in pharmaceutical development (45); we therefore explored such derivatization here for CD4mc indoline compounds. In addition, given the positive effect of 3-substituent hydrophobicity observed in the alkyl and aryl series, we were intrigued by the potential contribution of fluorine incorporation to this effect. Specifically, pentafluorobenzyl carbamate **30**, 1,1,1-trifluoroethyl carbamate **28** and 1,1,1-trifluoropropyl carbamate **29**, analogs of indolines **26**, **18** and **19**, respectively, were synthesized and tested for the ability to inhibit HIV-1_JR-FL_ infection. The fluorinated analogs **30** and **28** showed 2.1- to 2.7-fold enhanced antiviral activity compared to their respective fully hydrogenated counterparts **26** and **18** (Figure 3). Pleasingly, the pentafluorobenzyl carbamate in **30** (CJF-IV-046) resulted in a 23-fold improvement in viral inhibition compared to BNM-III-170. The antiviral activity of **29** was similar to that of **19**, indicating that the addition of fluorine did not enhance potency against HIV-1 in this instance. Thus, for some indoline CD4mcs, fluorination can be beneficial to antiviral potency (*Figure 3A*).

To examine the breadth of the observed antiviral activity, we tested selected indoline CD4mcs against HIV-1_AD8_, a different clade B virus, and against HIV-1_BG505_, a clade A virus that has proven to be more difficult to inhibit with current CD4mcs. The tested indoline CD4mcs (**17-23**, **26**-**30**) all inhibited HIV-1_AD8_ with IC_50_ values 3-10-fold better than those for HIV-1_JR-FL_ (*Figure 3A*). The clade A HIV-1_BG505_ was not susceptible to inhibition by our previous lead CD4mc, BNM-III-170 (**1**, IC_50_>300 μM) or other tested indane CD4mcs. By contrast, most of the tested *indoline* CD4mcs inhibited HIV-1_BG505_ infection with IC_50_ values in the micromolar range (*Figure 3A*). Of these CD4mcs, CJF-IV-046 (**30**), CJF-III-288 (**19**), CJF-III-192 (**26**) and DY-III-065 (**29**) stood out as the most potent CD4mcs, with BG505 IC_50_ values of 10.1 ± 6.8 μM, 27.0 ± 8.8 μM, 31.6 ± 5.7 μM and 35.0 ± 24.0 μM respectively. The antiviral IC_50_ values of the five most potent indoline CD4mcs and BNM-III-170, as determined in side-by-side assays, are shown in Table S3. To further investigate the underlying binding mechanism of these CD4mcs to HIV-1_BG505_, we then used molecular docking and the free energy of binding calculation using molecular mechanics/generalized born surface area (MM-GBSA) to evaluate binding mode of the CD4mcs and their correlation with experimental IC_50_ values (*Figure S5 and S6*). These results indicate that the carbamate substituents at the indoline N3 nitrogen can confer not only improved potency but also greater breadth against HIV-1 strains.

We next examined the ability of two potent indoline CD4mc analogs, CJF-III-288 (**19**) and CJF-IV-046 (**30**), to inhibit infection of TZM-bl target cells by a panel of global HIV-1 strains from phylogenetic clades A, B, C, D and G, as well as circulating recombinant forms AE, AG, BC and CD. With the exception of the clade AE HIV-1, which have a His 375 residue that fills the Phe 43 cavity, all of the viruses were inhibited by CJF-III-288 (**19**) and CJF-IV-046 (**30**); 92% and 98% of these viruses were inhibited by CJF-III-288 and CJF-IV-046, respectively, with IC_50_ values less than 15 μM (*Figure 3B, Table S1*). By contrast, 25% of the viruses were either not inhibited or required more than 15 μM to be inhibited by BNM-III-170 (**1**). The IC_50_ values of BNM-III-170 (**1**), CJF-III-288 (**19**) and CJF-IV-046 (**30**) for this panel of viruses were strongly correlated, with the slope of the regression line indicating an ~10-fold increase in potency for the two indoline CD4mcs over BNM-III-170 (*Figure 3B, Table S3, Figure S4*). These results indicate that the properties of the HIV-1 Env that determine sensitivity/resistance apply to both indane and indoline CD4mcs. A much weaker correlation was observed between the IC_50_ values of CJF-III-288 (**19**) and sCD4-Ig *(Figure S4)*.

### C. Cytotoxicity of the indoline CD4mcs

With the exception of DY-III-075 (**20**), CJF-IV-055 (**21**) and CJF-IV-056 (**22**), none of the indoline CD4mc analogs tested exhibited non-specific or toxic activity at concentrations up to 300 μM in the antiviral assays using Cf2Th-CD4/CCR5 target cells. However, at concentrations higher than 20 μM, CJF-IV-046 (**30**) exerted non-specific or toxic effects on the TZM-bl target cells used to evaluate the inhibition of infection by the panel of global HIV-1 strains (*Table S4*). To investigate potential effects on cell proliferation and viability, the potent indoline CD4mcs **19**, **20**, **26, 28**, **29** and **30** and BNM-III-170 were incubated with TZM-bl cells at varying concentrations (*Figure 3C*). We observed that CD4mcs bearing aromatic carbamate functionalities (**26**, **30**) inhibited TZM-bl cell proliferation beginning at a concentration of 50 μM. Most of the CD4mcs with alkyl carbamate substituents (**19**, **28**, **29**) did not affect cell viability; however, at high (100 μM) concentrations of **20**, toxicity for TZM-bl cells was observed. These results indicate that non-specific toxicity for TZM-bl cells is associated with indoline analogs with aromatic carbamates, a consideration for future indoline CD4mc design. Fortunately, most alkyl carbamate substituents appear to be non-toxic replacements for the aromatic carbamates, giving rise to CD4mcs that are equally effective in terms of antiviral potency. Moreover, the inclusion of fluorination in the ethyl or propyl carbamates (**28**, **29**) did not adversely affect cell viability. It is worth noting that at higher concentrations, DMSO also decreased the viability of TZM-bl cells.

### D. Crystal Structures of Indoline CD4mcs

In parallel with indoline CD4mc synthesis and evaluation of antiviral activity, we also analyzed crystal structures of selected compounds in complexes with the HIV-1_C1086_ gp120 core_e_ protein, which preferentially assumes a CD4-bound conformation (46). Pre-formed crystals were soaked with individual indoline CD4mcs [CJF-III-214 (**17**), CJF-III-289 (**18**), CJF-III-288 (**19**), CJF-III-192 (**26**), CJF-IV-047 (**28**), DY-III-065 (**29**), or CJF-IV-046 (**30**)], and the atomic structures were refined against X-ray diffraction measurements for the compounds at resolutions from 2.5 to 1.9 Å (*Table S1*). For comparisons, structures were also re-determined in this P2_1_2_1_2_1_ lattice at improved resolution for the previously reported complexes with the indane CD4mcs BNM-III-170 (38) and CJF-III-049-S (40). Electron density distributions were well defined, as shown in Figure 4B for **29** and for each ligand in Figures S1 A-I. The densities for each of the four copies in the asymmetric unit are highly similar (e.g., *Figure S2 A-D* for **29**); accordingly, we constrained the fitted gp120-CD4mc atomic structures to be identical.

**Figure 4.**
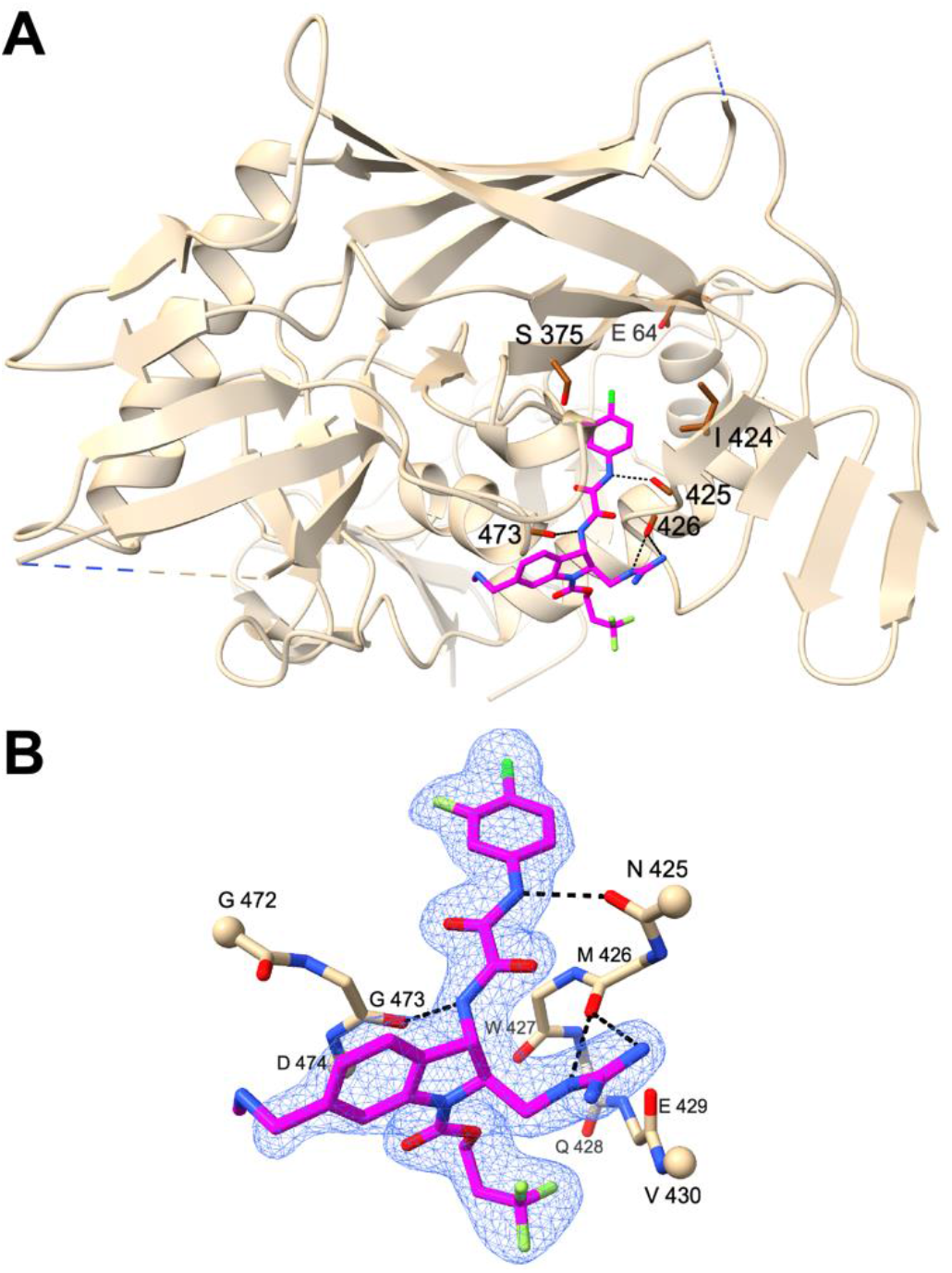
Crystal structure at 1.88 Å resolution of a representative indoline CD4mc, DY-III-065 (**29**), in complex with HIV-1_C1086_ gp120 core_e_. (**A**) DY-III-065 **(29)** in stick representation as complexed with gp120 displayed as a ribbon diagram (beige) and oriented as in Figure 1A. The carbonyl backbones of residues Asn425, Met426 and Gly473 of gp120 are in stick representation to demonstrate hydrogen bonds with DY-III-065. Hydrogen bonds are shown as black dashed lines. The sites of mutational resistance to indane CD4mc BNM-III-170 (Glu64, Ser375 and Ile424) are also shown in stick representation. The carbon atoms of DY-III-065, carbon atoms of gp120, and the oxygen, nitrogen, chlorine and fluorine atoms are shown in magenta, brown, red, blue, dark green and light green, respectively. (**B**) Crystal structure of DY-III-065 **(29)** as complexed with gp120 displayed in stick representation. The carbon atoms of gp120 are shown in beige. The C_α_ atoms of terminal residues in gp120 segments are represented in balls. The electron density is from a 2Fo-Fc synthesis contoured at about 1.5σ and selected for coverage within 2 Å of ligand atoms.

Indoline CD4mcs bind to gp120 in a manner typified by DY-III-065 in our best-defined structure at 1.88 Å resolution (Figure 4A), with interactions like those made by indane CD4mc BNM-III-170 (Figure 1B). The gp120 binding modes of all seven indoline CD4mcs are the same (Figures 5A and S3A), featuring hydrogen-bonding to main-chain carbonyls (Figure 4B). The substituents at indane positions 1, 2 and 5 of BNM-III-170 (**1**) are preserved identically in CJF-III-049-S (**2**) and in all of the indoline CD4mcs in this study. The methylamino methyl group at C5 makes no gp120 contacts, but the other two are stereo-specifically fixed by hydrogen bonding to the gp120 backbone. Thus, despite varied substitutions at N3 or C3, the binding poses of the indoline CD4mcs are similar to those of the indane CD4mcs (Figures 5B and S3B); however, because of geometric differences between the indoline and indane CD4mcs in the dispositions of the fixed C1 and C2 substituents on the five-membered rings, their very similar ring skeletons are oriented differently when bound to gp120 (Figure 5B). As a consequence, the aromatic six-membered ring is stacked against the G473-D474 peptide unit (Figure 5C) more closely for the indoline CD4mcs, which should provide greater π-π overlap and strengthened binding. Moreover, this redisposition in the indoline CD4mcs (Figure 5D) relative to the indane CD4mcs (Figure 5E) also moves several non-bonded contacts into highly favorable van der Waals interactions.

**Figure 5.**
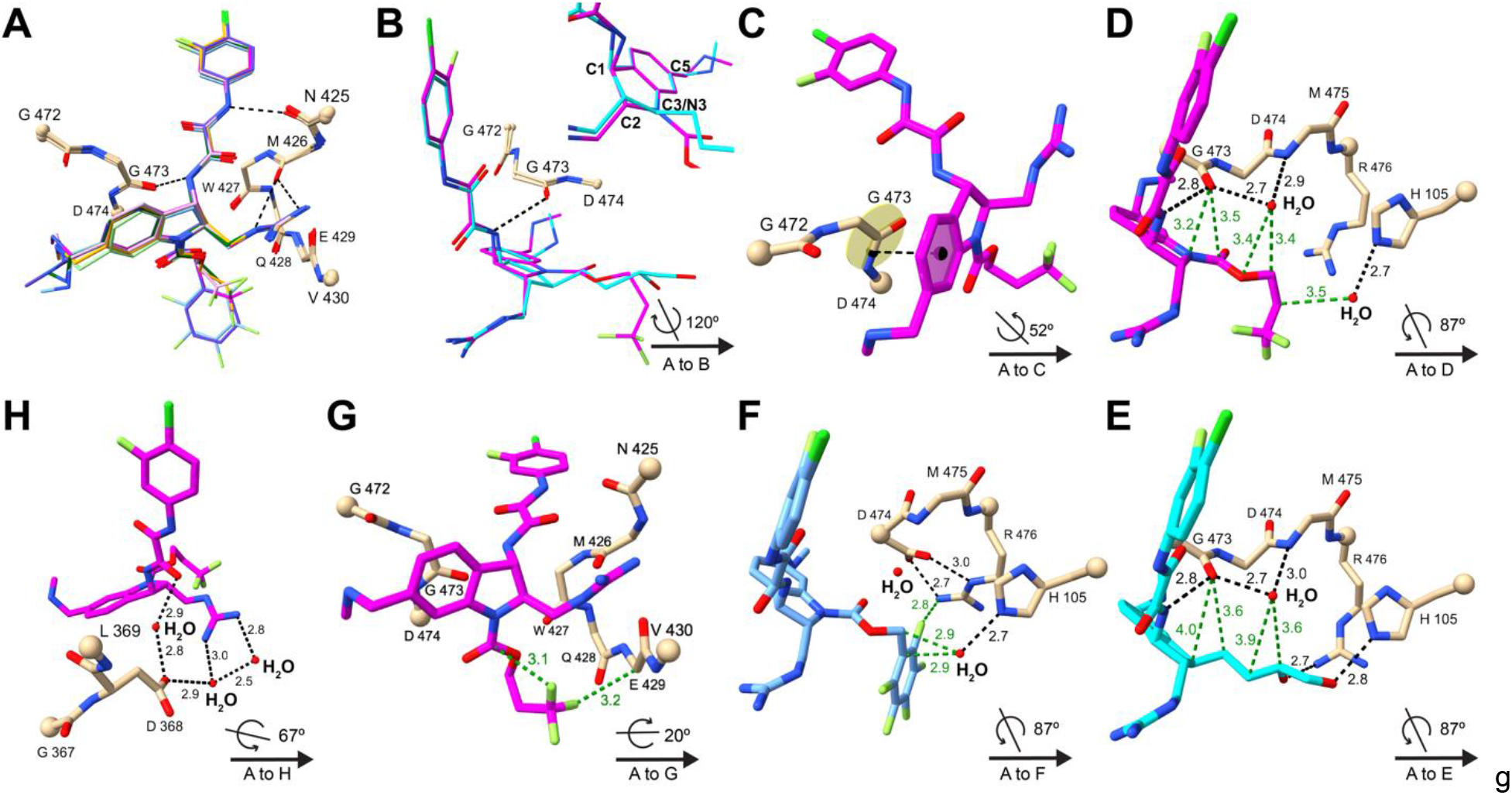
Crystal structures of CD4mcs in complex with HIV-1_C1086_ gp120 core_e_. **(A)** Comparison among crystal structures of indoline CD4mcs after superimposition of the gp120 cores: CJF-III-214 **(17)**, light green; CJF-III-289 **(18)**, dark green; CJF-III-288 **(19)**, orange; CJF-III-192 **(26)**, purple; CJF-IV-047 **(28)**, pink; DY-III-065 **(29)**, magenta; and CJF-IV-046 **(30)**, light blue. Carbon atoms of gp120 are in beige; other atoms are colored as in Figure 4. The C_α_ atoms of terminal residues in gp120 segments are represented in balls. A key is given at lower right in each of panels **(B)**-**(H)** for the orientation of its structure relative to that of **(A)**. (**B**) Comparison between indoline CD4mc DY-III-065 **(29)** and indane CD4mc CJF-III-049-S **(1)** after superimposition of the gp120 cores. The carbon atoms of DY-III-065 and CJF-III-049-S are in magenta and cyan, respectively. The insert image compares DY-III-065 and CJF-III-049-S after superimposition of their aromatic six-membered rings. (**C**) π-π interactions between the aromatic six-membered ring of DY-III-065 **(29)** and the G473-D474 peptide unit of gp120. The black dashed line connects the centroid of the aromatic six-membered ring of DY-III-065 and center of the G473-D474 peptide bond. (**D-F**) Hydrogen bonds and non-bonded contacts that DY-III-065 **(D)**, CJF-III-049-S **(E)** or CJF-IV-046 **(F)** form with gp120 and surrounding water molecules. Hydrogen bonds and non-bonded contacts are shown as black and green dashed lines, respectively. (**G**) Fluorine atoms of DY-III-065 **(29)** form favored non-bonded contacts with the C_α_ of E429 in gp120 and with its own carbamate carbon. (**H**) A water network at the protein-ligand interface in the structure of the DY-III-065 **(29)** complex associates the strictly conserved gp120 D368 carboxylate with the guanidinium group from C2 of the CD4mc.

The N3 carbamate ester substituents of the most efficacious indoline CD4mcs studied here (**19**, **29** and **30**) all occupy spaces in the Phe 43 vestibule of gp120 that are similar to, but distinctive from that for the C3 butanediol substituent of indane CD4mc **2** (Figures 5E and S3B). The carbamate ester cores of these indolines are chemically and structurally the same through the first carbon of the carbamate substituent, yet the antiviral activities of these compounds increase from the methyl to propyl, benzyl, trifluoropropyl and pentafluorobenzyl by factors of 4.3, 2.0, 1.4 and 4.2, respectively. The paucity of gp120 contacts from the carbamate substituents confounds the analysis, especially as compared with those from the C3 substituent in indane **2** (Table S2), which shows many more contacts despite having activities >17-fold lower than the best indoline CD4mcs (**19**, **29** and **30**). The benzyl carbamate (**26**) and its pentafluorobenzyl counterpart (**30**) adopt identical conformations as bound to gp120 (Figure S3C); and the propyl (**19**) and trifluoropropyl (**29**) carbamate are also very similar (Figure S3D). Thus, the fully hydrogenated and partially fluorinated pairs make very similar gp120 contacts, altogether hydrophobic for the propyl carbamate; nevertheless, fluorine atoms in **30** (Figures 5F) and **29** (Figure 5G) participate in contacts with the gp120 core_e_ protein.

The structure of the gp120 core_e_ complex with DY-III-065 (**29**) reveals water structure at protein-ligand interfaces. One water site, shown in Figure 5D, is hydrogen-bonded to the carbonyl oxygen of G473, the amino group of M475, and the carbonyl oxygen of W427 (not shown) and just outside hydrogen-bonding distance from the carbamate-linking oxygen atom. A second set of waters associates the C2 guanidinium group of **29** with the carboxyl group of the stringently conserved D368 of gp120, which is critical for CD4 binding (23). The C2 guanidinium nitrogens of **29** H-bond to two water molecules, one of which H-bonds to D368; water-mediated H-bonding also bridges D368 to a C1 oxalamide oxygen of **29** (Figure 5H). These water sites are also present in the indane CD4mc complexes, but they only become clearly evident here with the highest resolution yet achieved for this system. Synthetic efforts to create new indoline CD4mc analogs that mimic these water-based interactions with D368 are in progress.

It is noteworthy that interactions between DY-III-065 (**29**) and gp120 primarily involve main chain atoms or the side chain of the conserved CD4-binding residue D368 (23, 24). Water molecules that mediate the contacts between **29** and the side chains of polymorphic HIV-1 gp120 residues (e.g., His/Gln 105) can potentially form hydrogen bonds despite this variation. These characteristics should militate against the evolution of HIV-1 escape mutants.

### E. Antiviral Activity of Indoline CD4mcs Against Resistant HIV-1 Variants

Previously, CD4mc-resistant variants of HIV-1_AD8_ were selected by incubating infected cultures with increasing concentrations of BNM-III-170 (39). After selection, the derived mutant HIV-1_AD8_ 130-C was completely resistant to entry inhibition by concentrations of BNM-III-170 up to 300 μM. In this case, two changes in the Phe 43 cavity (I424T and S375N) and a change (E64G) in Layer 1 of the gp120 inner domain individually contributed to BNM-III-170 resistance, and acted additively to increase resistance to BNM-III-170 (39).

Given the improved breadth of infection inhibition exhibited by the indoline CD4mcs, we were curious if this class of CD4mcs could overcome these resistance-associated adaptations and inhibit the infection of these HIV-1 mutants. The evolved BNM-III-170-resistant HIV-1_AD8_ mutant 130-C and the HIV-1_AD8_ mutant containing the three resistance-associated gp120 changes (E64G+S375N+I424T) were inhibited by high concentrations of CJF-III-192 (**26**) (*Table 1*); the CD4mcs CJF-III-288 (**19**) and CJF-IV-046 (**30**) were less effective against these mutants (*Table 1*). The indoline CD4mcs were also evaluated for their ability to inhibit the HIV-1_AD8_ mutants containing single or pairwise combinations of the resistance-associated gp120 changes. All of the tested indoline CD4mcs (**19, 26** and **30**) inhibited the single-residue HIV-1_AD8_ mutants with IC_50_ values less than 10 μM. The inhibition of infection was diminished for the pairwise combinations of the gp120 changes; however, variants containing all three permutations of the pairwise mutations were still inhibited by the indoline CD4mcs **19**, **26** and **30** with IC_50_ values less than 75 μM. In all cases, the indoline CD4mcs were able to inhibit these resistance-selected HIV-1_AD8_ mutants more effectively than BNM-III-170 (39). Thus, although the gp120 changes that determine resistance to our current lead indane CD4mc, BNM-III-170, also result in relative resistance to the indoline CD4mcs, the increased potency of the indoline CD4mcs allows measurable inhibition of HIV-1 variants that are resistant to BNM-III-170.

**Table 1.**
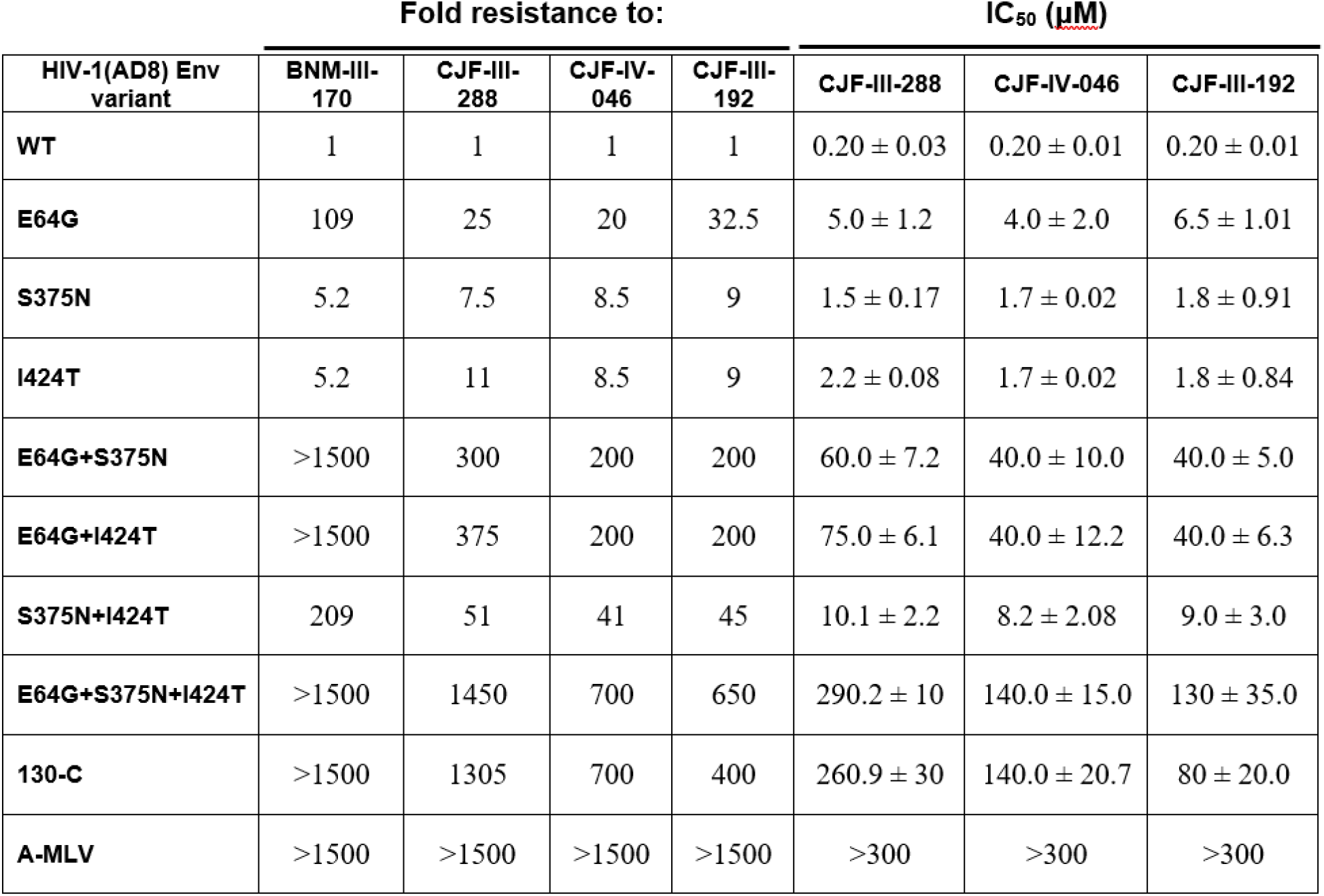
Sensitivity of BNM-III-170-resistant HIV-1 variants to inhibition by indoline CD4mcs The sensitivity of recombinant luciferase-expressing viruses pseudotyped with the indicated HIV-1_AD8_ Env variants to the indoline CD4mcs was determined using Cf2Th-CD4/CCR5 target cells, as described in Materials and Methods. The IC_50_ values (in μM) are shown in the right three columns. The level of resistance relative to that of the wild-type (WT) AD8 Env is shown in the left columns; the values for the indane CD4mc, BNM-III-170, were derived from reference 39.

### F. ADCC and Antibody Recognition

CD4mcs have the potential to sensitize HIV-1-infected cells to killing by ADCC (30). We therefore examined the ability of the indoline CD4mc inhibitors: (1) to enhance recognition of Env on the surface of infected cells by antibodies (Ab) present in plasma from HIV-1-infected individuals (HIV+ plasma); (2) to mediate recognition of antibody-bound infected cells by FcγRIIIa receptors in the presence of HIV+ plasma; and (3) to sensitize infected cells to ADCC mediated by HIV+ plasma. Potent indoline CD4mcs were evaluated first for their ability to enhance HIV+ plasma binding to primary CD4+ T cells infected by HIV-1_JR-FL_. As demonstrated in *Figure 6A and D*, the indoline CD4mcs CJF-III-288 (**19**), CJF-III-192 (**26**), and CJF-IV-046 (**30**) enhanced the binding of plasma from HIV-1-infected individuals to HIV-1_JR-FL_-infected cells more efficiently than BNM-III-170. We also evaluated whether the CD4mcs were able to enhance HIV+ plasma binding to HIV-1_JR-FL_-infected cells and promote subsequent FcyRIIIa receptor engagement (*Figure 6B and E*). All of the indoline CD4mcs examined [CJF-III-288 (**19**), CJF-III-192 (**26**), and CJF-IV-046 (**30**)] improved recombinant soluble dimeric FcyRIIIa engagement by HIV+ plasma to a greater degree than BNM-III-170. In addition, the indoline CD4mcs CJF-III-288 (**19**), CJF-III-192 (**26**) and CJF-IV-046 (**30**) all enhanced ADCC killing of HIV-1_JR-FL_-infected CD4+ T cells to a greater extent than BNM-III-170 (*Figure 6C and F*).

**Figure 6.**
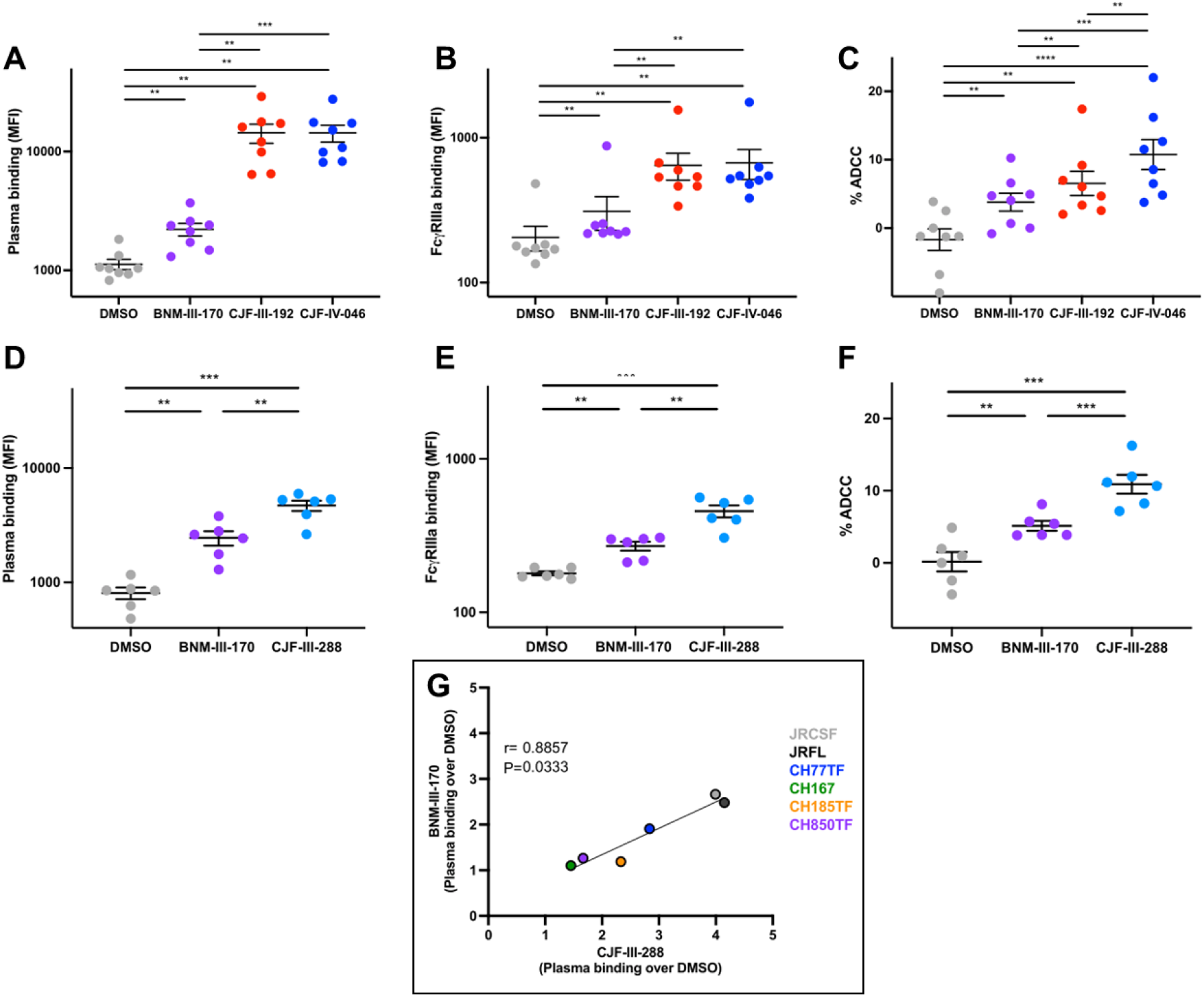
Indoline CD4mcs enhance recognition of HIV-1-infected cells by HIV+ plasma, enabling ADCC. (**A-F**) Primary CD4+ T cells were infected with HIV-1_JR-FL_ and used as targets for HIV+ plasma and FcγRIIIa binding and for their susceptibility to HIV+ plasma mediated-ADCC. (**A**) HIV+ plasma binding (MFI) to CD4+ T cells infected with HIV-1_JR-FL_ in the presence of DMSO or the indicated CD4mc. (**B**) FcyRIIIa binding (MFI) in the presence of HIV+ plasma and DMSO or a CD4mc. (**C**) HIV+ plasma-mediated ADCC (%) in the presence of DMSO or the indicated CD4mc. (**D**) HIV+ plasma binding (MFI) in the presence of DMSO or a CD4mc. (**E**) FcyRIIIa binding (MFI) in the presence of HIV+ plasma and DMSO or a CD4mc. (**F**) HIV+ plasma-mediated ADCC (%) in the presence of DMSO or a CD4mc. (**G**) Comparison of BNM-III-170 and CJF-III-288 stimulation of HIV+ plasma binding to primary CD4+ T cells infected with the indicated HIV-1 strain. Statistical significance was tested using (**A-F**) paired *t* test or Wilcoxon test based on statistical normality (**P<0.01; ***P<0.001 ****P<0.0001) and (**G**) Spearman rank correlation test. Concentrations of CD4mc used in each assay were: BNM-III-170 (50 μM), CJF-III-192 (50 μM), CJF-IV-046 (25 μM), or CJF-III-288 (50 μM).

In addition to enabling the recognition of HIV-1_JR-FL_-infected CD4+ T-cells by HIV+ plasma, indoline CD4mc CJF-III-288 (**19**) enhanced HIV+ plasma binding to primary CD4+ T cells infected with a panel of HIV-1 strains ~2-fold more effectively than BNM-III-170 (*Figure 6G*).

## Discussion

CD4mcs are unique small-organic-molecule HIV-1 entry inhibitors that competitively block CD4 binding, trigger premature conformational changes in the Env leading to functional inactivation, and “open” Env, sensitizing virions to antibody neutralization and infected cells to ADCC. Here, we report that certain indoline CD4mc analogs exhibit 10-20-fold increases in specific anti-HIV-1 potency, compared with the earlier lead indane CD4mc, BNM-III-170. Accompanying this increase in potency, we observed greater breadth of activity against a range of primary HIV-1 strains. Natural HIV-1 variants exhibit approximately 1000-fold differences in sensitivity to indane CD4mcs, with a substantial fraction of variants requiring greater than 100 μM concentrations of BNM-III-170 to achieve 50% inhibition of infection. Remarkably, regardless of their phylogenetic classification into clade A, B, C, D, AG, BC or CD, diverse HIV-1 strains exhibited increased sensitivity to the more potent indoline CD4mc analogs. Only clade AE recombinant HIV-1, which has the imidazole side chain of His 375 occupying the Phe 43 cavity, were resistant to potent indoline CD4mcs such as CJF-III-288.

Likewise, an HIV-1_AD8_ variant (130-C) selected for resistance to BNM-III-170, as well as recombinant HIV-1 containing key resistance-associated changes in Env, were found to be inhibited more efficiently by the indoline CD4mcs than by BNM-III-170. Apparently, Env changes near or within the Phe 43 cavity (such as S375N and I424T in the 130-C virus) that are unfavorable for CD4mc binding can be compensated by contacts made by the indoline CD4mcs with the gp120 vestibule. These novel contacts of the indoline CD4mcs also appear to compensate for the more distant E64G change, which may decrease the propensity of Env to stabilize the CD4-bound conformation favored by the CD4mc (39). Thus, although the indoline CD4mcs are still affected by the resistance-associated gp120 changes, the overall increase in the relative antiviral potency of the indoline CD4mcs allows inhibition of the resistant viruses at achievable concentrations.

Although further study is needed, the increased antiviral potency of the indoline CD4mcs, compared with BNM-III-170, likely derives from novel contacts with the gp120 surface made by the substituents on the N3 nitrogen. Crystal structures suggest that these contacts involve a conserved hydrophobic pocket at the junction of the gp120 inner and outer domains (*Figure 1B*). This hydrophobic pocket is flanked by the polymorphic residues His/Gln 105 in the α1 helix and Arg/Lys 476 in the α5 helix. These natural polymorphisms apparently exert only minimal effects on HIV-1 sensitivity to the more potent indoline CD4mcs, because several susceptible natural variants exhibit polymorphic changes in these residues. These observations suggest that indoline CD4mc potency may benefit from interactions of the N3 substituent that are tolerant of some natural HIV-1 diversity. A consequence of the additional gp120 contacts made by the N3 substituent may thus be better stabilization of the binding pose shared by the indoline and indane cores. This pose promotes hydrogen bonding with multiple mainchain atoms (Asn 425, Met 426, Trp 427, Gly 473) that contribute to the ability of CD4mcs to recognize gp120 glycoproteins from multiple strains.

Strong correlations were also observed between the sensitivities of HIV-1 variants to the indane CD4mc BNM-III-170 and to potent indoline CD4mcs, CJF-III-288 and CJF-IV-046. As the pseudotyped recombinant viruses used in the assays differ only in Env, the observed correlations indicate that Env-specific variables determine sensitivity to CD4mcs in general. The nature of the Env variables accounting for the wide range of sensitivities to CD4mcs is clearly worthy of further investigation. We note that polymorphisms in residues 105 and 476, which can be contacted by some indoline analogs, but not by BNM-III-170, do not result in deviations from the linear correlation in *Figure 3B*, supporting the assertion that these natural changes exert minimal impact on the efficacy of the more potent indoline CD4mcs.

Finally, three of the indoline CD4mcs, CJF-III-192, CJF-III-288 and CJF-IV-046, were more effective than BNM-III-170 in exposing vulnerable Env epitopes on the surface of HIV-1-infected cells relevant to ADCC. These results justify ongoing studies in animal models to evaluate the ability of indoline CD4mc to sensitize HIV-1-infected cells to ADCC.

Indoline CD4mc with improved potency should expedite exploration of the therapeutic and prophylactic potential of these interesting small molecules.

## Materials and Methods

### Ethics statement

Written informed consent was obtained from all study participants [the Montreal Primary HIV Infection Cohort (47, 48) and the Canadian Cohort of HIV Infected Slow Progressors (49–51) and research adhered to the ethical guidelines of CRCHUM and was reviewed and approved by the CRCHUM Institutional Review Board (Ethics Committee, approval number CE 16.164 - CA). Research adhered to the standards indicated by the Declaration of Helsinki. All participants were adult and provided informed written consent prior to enrollment in accordance with Institutional Review Board approval.

### Cell lines and primary cells

293T cells and TZM-bl cells were grown in Dulbecco’s modified Eagle’s medium (DMEM) (Life Technologies, Wisent Inc.) supplemented with 10% fetal bovine serum (FBS) (Life Technologies, VWR) and 100 μg/ml of penicillin-streptomycin (Life Technologies, Wisent Inc.). Cf2Th cells stably expressing the human CD4 and CCR5 coreceptors for HIV-1 were grown in the same medium supplemented with 0.4 mg/ml of G418 and 0.2 mg/ml of hygromycin.

Human peripheral blood mononuclear cells (PBMCs) from healthy HIV-1-negative individuals were obtained by leukapheresis and Ficoll-Paque density gradient isolation and were cryopreserved in liquid nitrogen until use. CD4+ T lymphocytes were purified from resting PBMCs by negative selection using immunomagnetic beads according to the manufacturer’s instructions (StemCell Technologies). The cells were activated with phytohemagglutinin-L (10 μg/mL) for 48 h and then maintained in RPMI 1640 complete medium supplemented with recombinant interleukin-2 (rIL-2) (100 U/mL).

#### Viral production and infection

Vesicular stomatitis virus G (VSVG)-pseudotyped viruses were produced and titrated as previously described (52). Viruses were used to infect activated primary CD4+ T cells from healthy HIV-1-negative donors by spin infection at 800 x g for 1 h in 96-well plates at 25°C.

### CD4-mimetic compounds (CD4mcs)

The CD4mc analogs were synthesized as described in detail in the Supporting Information. The compounds were dissolved in DMSO at a stock concentration of 10 mM and stored at −20°C until use. The CD4mcs were diluted to 1 mM in DMEM for pseudovirus inhibition assays, 50 or 25 μM in PBS for cell-surface staining and 50 or 25 μM in RPMI 1640 complete medium for ADCC assays.

### Antibodies and plasma

The sCD4-Ig used in the virus inhibition assays is described in reference 39. The anti-coreceptor binding site 17b monoclonal antibody (mAb) (NIH HIV Reagent Program) and plasma from HIV-1-infected individuals were used to assess cell-surface Env conformation. Plasma from the Montreal Primary HIV Infection Cohort (47, 48) and the Canadian Cohort of HIV Infected Slow Progressors (49–51) were collected, heat-inactivated and conserved at −80 °C until use. The conformation-independent anti-gp120 outer-domain 2G12 mAb (NIH HIV Reagent Program) was used to normalize Env expression. Goat anti-human IgG (H+L) (Thermo Fisher Scientific) antibodies pre-coupled to Alexa Fluor 647 and streptavidin conjugated to Alexa Fluor 647 were used as a secondary antibody in flow cytometry experiments.

### Inhibition of HIV-1 infection by CD4mcs

HIV-1 inhibition assays were performed as described previously (29,39,43). To produce recombinant luciferase-expressing HIV-1, 293T cells were transfected with plasmids expressing HIV-1 Env variants and Rev, the pCMVΔP1Δenv HIV-1 Gag-Pol packaging construct and the firefly luciferase-expressing HIV-1 vector at a 1:1:3 μg DNA ratio using Effectene transfection reagent (Qiagen). Recombinant luciferase-expressing viruses capable of a single round of replication were released into the cell medium and were harvested 48 h later. The virus-containing supernatants were clarified by low-speed centrifugation (600 x g for 10 min) and used for single-round infections. To measure inhibition of infection, the recombinant viruses were incubated with the CD4mcs for 1 h at 37°C. The mixture was then added to Cf2Th-CD4/CCR5 target cells expressing CD4 and CCR5. After 48 h of culture, the target cells were lysed and the luciferase activity was measured.

The pseudotyped virus inhibition assays used to generate the data in Figure 3B and Table S1 were conducted as previously described (38). Single-round HIV-1 pseudotyped by the Envs from a panel of multiclade HIV-1 were incubated for 1 h at 37°C with 5-fold serial dilutions of CD4mc or sCD4-Ig in duplicate. TZM-bl cells were then added in growth medium containing DEAE-dextran at a final concentration of 11 μg/ml. The assay plates were incubated for 48 h at 37°C, 5% CO_2_, after which the cells were lysed and luciferase reporter gene expression was measured using Bright-Glo luciferase reagent (Promega) and a Victor 3 or GloMax Navigator luminometer (Perkin Elmer/Promega). The 50% and 80% inhibitory concentrations (IC_50_ and IC_80_, respectively) were calculated based on the relative luciferase unit (RLU) activity in the treated sample compared with that in the untreated control, after subtraction of the background RLU in control uninfected TZM-bl cells.

### Cell viability assay

TZM-bl cells were seeded in 96-well assay plates containing 100 μl of culture medium. CD4mcs in stock solutions of 100% DMSO were added at the specified final concentrations in triplicates. For the DMSO control, a volume of DMSO was added that corresponded to the volume of the CD4mc added at each indicated concentration. After seventy-two h, cell viability was measured using the CellTiter 96® Non-Radioactive Cell Proliferation Assay (MTT) kit from Promega. Briefly, 15 μl of dye solution (from the kit) was added to each well followed by incubation of the plate at 37°C for 4 h in a humidified CO_2_ incubator. One hundred μl of solubilization/stop solution (from the kit) was then added to each well. Absorbance at 570 nm and 630 nm was measured using a 96-well plate reader.

### Flow cytometry analysis of cell-surface staining

Cell surface staining of primary infected cells was performed 48 h post-infection as previously described (30). HIV-1-infected cells were incubated for 1 h at 37 °C with HIV+ plasma (1:1000 dilution) in the presence of CD4mcs (25 or 50 μM) or with equivalent volume of vehicle (DMSO). Cells were then washed once with PBS and stained with appropriate Alexa Fluor-647-conjugated (Invitrogen) secondary Abs (2 μg/mL) for 20 minutes at room temperature. Alternatively, the binding of HIV+ plasma was detected using a biotin-tagged dimeric recombinant soluble FcγRIIIa (0.2 μg/mL) followed by the addition of Alexa Fluor 647-conjugated streptavidin (Thermo Fisher Scientific; 2 μg/mL). Infected cells were then stained intracellularly for HIV-1 p24, using the Cytofix/Cytoperm Fixation/ Permeabilization Kit (BD Biosciences, Mississauga, ON, Canada) and the fluorescent anti-p24 mAb (PE or FITC-conjugated anti-p24, clone KC57; Beckman Coulter/Immunotech). The percentage of infected cells (p24+ cells) was determined by gating the living cell population on the basis of AquaVivid viability dye staining. Samples were analyzed on an LSRII cytometer (BD Biosciences), and data analysis was performed using FlowJo vX.0.7 (Tree Star, Ashland, OR, USA).

### ADCC FACS-based assay

Measurement of ADCC using the FACS-based assay was performed at 48 h post-infection as previously described (30, 34). Briefly, HIV-1_JR-FL_**-i**nfected primary CD4+ T cells were stained with viability (AquaVivid; Thermo Fisher Scientific) and cellular (cell proliferation dye eFluor670; eBioscience) markers and used as target cells. Autologous PBMC effectors cells, stained with another cellular marker (cell proliferation dye eFluor450; eBioscience), were added at an effector: target ratio of 10:1 in 96-well V-bottom plates (Corning, Corning, NY). Briefly, infected primary CD4+ T cells were incubated with HIV+ plasma (1:1000), in the presence of CD4mcs (25 or 50 μM) or an equivalent volume of vehicle (DMSO). The plates were subsequently centrifuged for 1 min at 300 g, and incubated at 37°C, 5% CO_2_ for 4 to 6 h before being fixed in a 2% PBS-formaldehyde solution. Cells were then stained intracellularly for HIV-1 p24 as described above. Samples were analyzed on an LSRII cytometer (BD Biosciences). Data analysis was performed using FlowJo vX.0.7 (Tree Star). The percentage of ADCC was calculated with the following formula: (% of p24+ cells in Targets plus Effectors) − (% of p24+ cells in Targets plus Effectors plus plasma) / (% of p24+ cells in Targets) by gating on infected live target cells.

### Statistical analysis

Statistics were analyzed using GraphPad Prism version 9.4.1 (GraphPad, San Diego, CA, USA). Every data set was tested for statistical normality and this information was used to apply the appropriate (parametric or nonparametric) statistical test. P values <0.05 were considered significant; significance values are indicated as * P<0.05, ** P<0.01, *** P<0.001, **** P<0.0001.

### Protein Purification and X-ray Crystallography

The plasmid expressing the gp120 core_e_ protein of the clade C HIV-1_1086_ was donated by Lei Chen from Peter Kwong’s laboratory (National Institutes of Health) (46). An 8-histidine (8His) tag was added to the C-terminus of the gp120 core_e_ protein to assist affinity purification. About 1 mg of the expressor plasmid was transfected into 1 liter of HEK293S GnTI- suspension cells at 2 x 10^6^ cells/mL. After expression of the core_e_ protein, the supernatant was filtered using a 0.45 μm filter and passed through a pre-packed Ni-NTA column. The Ni-NTA column was washed with 40 mM imidazole in 1x phosphate-buffered saline (PBS) and the 8His-tagged gp120 core_e_ protein was eluted with 250 mM imidazole in 1x PBS. The eluted gp120 core_e_ protein was deglycosylated overnight in a 37 °C water bath with Endoglycosidase Hf (New England Biolabs). Afterwards, the deglycosylated gp120 was further purified using Concanavalin A (Con A)-Sepharose (Cytiva) to remove any incompletely deglycosylated proteins. The flowthrough of the Con A-Sepharose column was collected and further purified using a pre-packed Ni-NTA column, which removed Endoglycosidase Hf in the flowthrough. The completely deglycosylated gp120 core_e_ protein was eluted with 250 mM imidazole in 1x PBS and further polished with a Superdex 200 Increase column (Cytiva). The purified deglycosylated gp120 core_e_ protein was concentrated to 10 mg/mL for crystallization.

Crystallization of the unliganded gp120 core_e_ protein was performed at 20 °C using the hanging drop vapor diffusion method. The crystals grew in drops consisting of 1 μL of protein solution and 1 μL of reservoir solution to equilibrate against 600 μL of reservoir solution 14 to 16 %(w/v) PEG 1500, 0.1 M CaCl_2_, 0.1 M imidazole pH 6.5. For each experiment, the compound of interest was dissolved in 100% DMSO. A single crystal was picked from the mother liquor and soaked in 5 μL of a stabilization buffer that contained 26% PEG 1500 (w/v), 0.1 M CaCl_2_, 0.1 M imidazole pH 6.5, 2.5 mM Tris-HCl pH 7.5, 350 mM NaCl, 0.02% NaN_3_, 5% (v/v) DMSO and 2 mM of the compound. The gp120 core_e_ crystals were soaked for 30 to 60 min in the stabilization buffer and then transferred for 5 seconds in cryo-protectant, which is the stabilization buffer but with 30% ethylene glycol. Diffraction data were collected on 24ID-C and 24ID-E beamline at Advanced Photon Source (APS) at Argonne National Laboratory. Crystal structures were solved by the molecular replacement module in PHENIX using the unliganded HIV-1C_1086_ gp120 core_e_ structure (PDB ID: 3TGR) and refined with phenix.refine.

## Data Availability

Data deposition: Atomic coordinates and structure factors have been deposited in the Research Collaboratory for Structural Bioinformatics (RCSB) Protein Data Bank, www.rcsb.org under accession code 8FLY for BNM-III-170 (53), 8FLZ for CJF-III-049-S (54), 8FM0 for CJF-III-214 (55), 8FM2 for CJF-III-289 (56), 8FM3 for CJF-III-288 (57), 8FM4 for CJF-IV-047 (58), 8FM5 for DY-III-065 (59), 8FM7 for CJF-III-192 (60) and 8FM8 for CJF-IV-046 (61).

## Acknowledgments

We thank Irwin Chaiken and all the members of the P01 Consortium Structure-Based Antagonism of HIV-1 Envelope Function in Cell Entry (NIH Grant No. AI150471). We also thank the staff at NE-CAT beamlines at APS for their support during data collection. Dr. Charles Ross, and Dr. Jun Gu (University of Pennsylvania) are also acknowledged for their assistance obtaining mass and NMR spectra, respectively. The authors also thank Dr. Mark Hogarth for kindly providing recombinant dimeric FcgRIIIa and Dr Beatrice Hahn for providing the following infectious molecular clones: HIV_CH77TF_, HIV_CH185TF_, HIV_CH850TF_ and HIV_CH167_. A.F. is the recipient of a Canada Research Chair on Retroviral Entry #RCHS0235 950-232424.

