## Supporting Information for "Indoline CD4-mimetic Compounds Mediate Potent and Broad HIV-1 Inhibition and Sensitization to Antibody-dependent Cellular Cytotoxicity"

[1] Authors contributed equally to the preparation of this manuscript

[2] Corresponding authors

**This PDF file includes:**

Figures S1 to S5

Tables S1 to S3

Experimental Methods

$^{1}$H and $^{13}$C NMR Spectra

### Supporting Information

#### Summary of Crystallographic Data

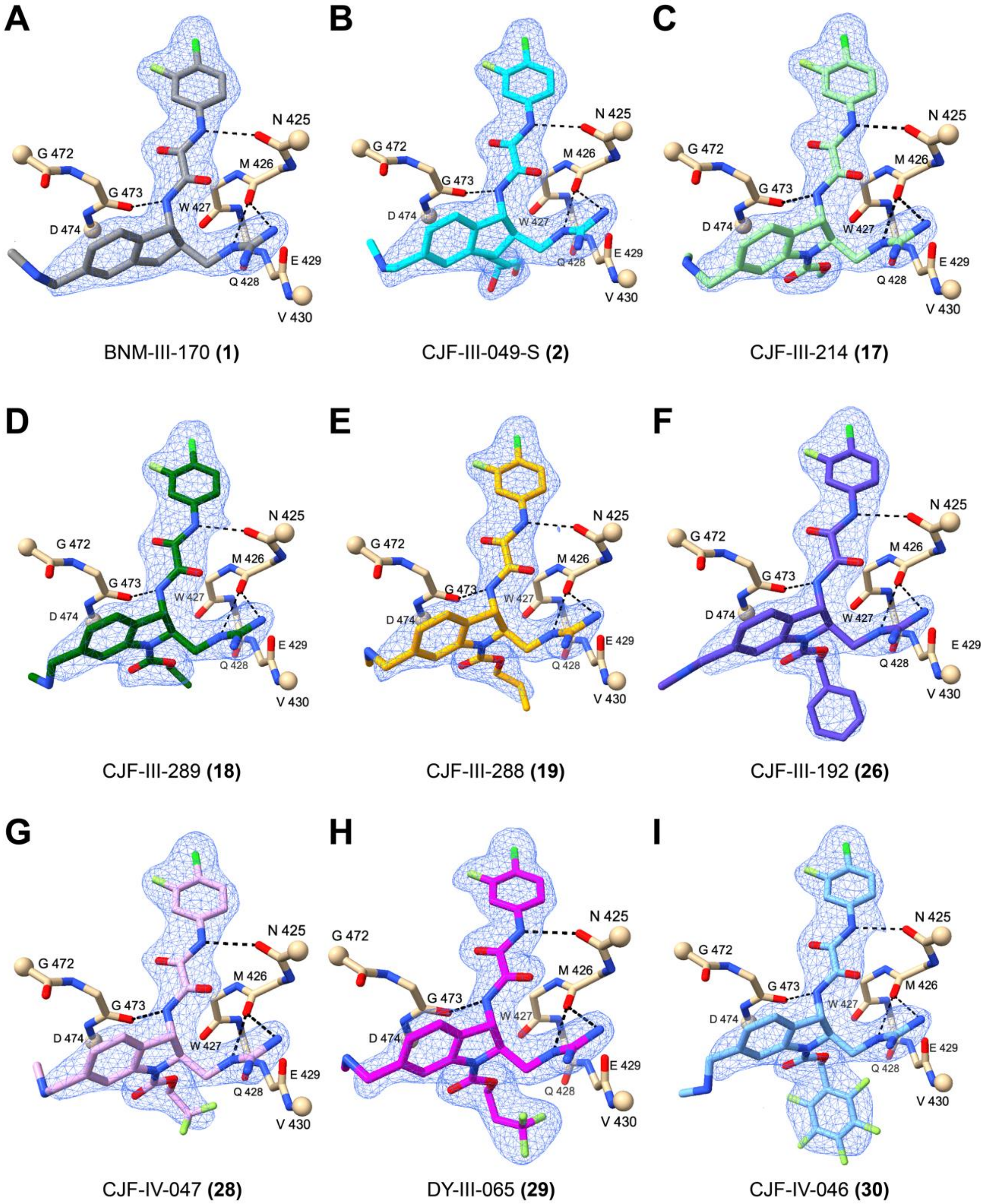

**Figure S1. Electron density distributions are well defined for each CD4mc in its complex with gp120.** The densities are from 2Fo-Fc syntheses of chain A as contoured at about 1.5σ and selected for coverage within 2 Å of ligand atoms. Hydrogen bonds are shown as black dashed lines. A ligand-distinctive color is used for the carbon atoms in each ligand. The gp120 carbon atoms, and oxygen, nitrogen, chlorine and fluorine atoms are shown in beige, red, blue, dark green and light green, respectively. The $C_{\alpha}$ atoms of terminal residues in the gp120 segments are drawn as balls.

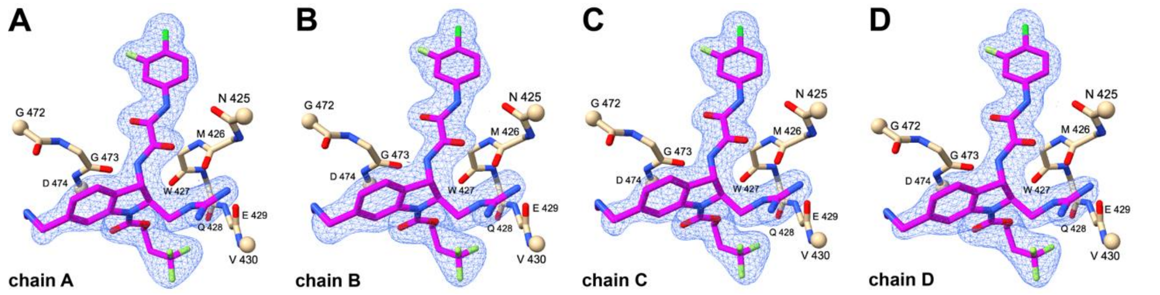

**Figure S2. The electron densities for each of the four copies in the asymmetric unit of $P2_12_12_1$ lattice are highly similar.** Each density is from a 2Fo-Fc synthesis of the gp120 complex with DY-III-065 as contoured at about 1.5σ and selected for coverage within 2 Å of ligand atoms.

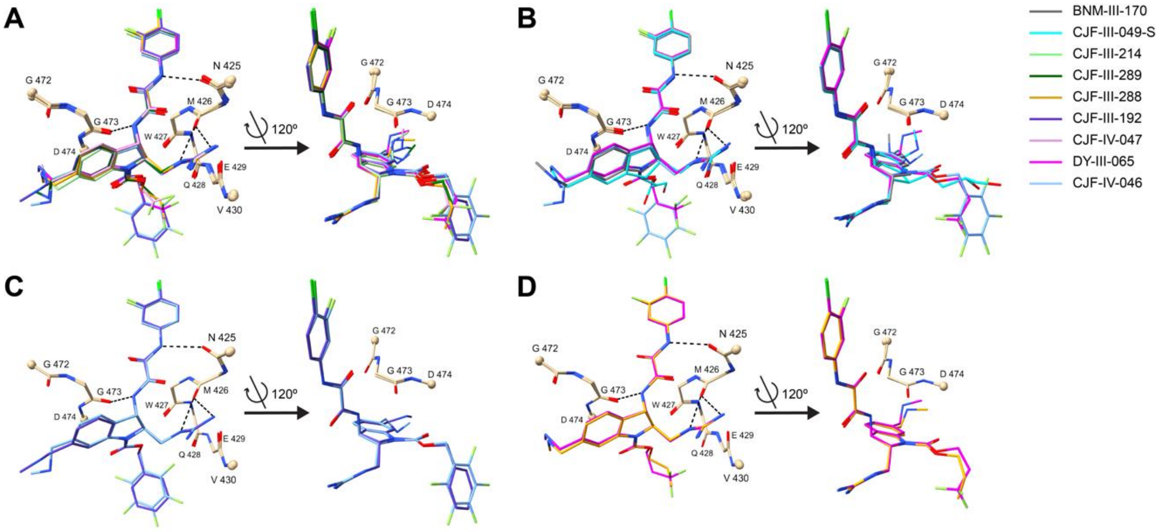

**Figure S3. Comparisons among indoline CD4mcs and indane CD4mcs after superimposition of the corresponding gp120 cores. (A)** Comparison among crystal structures of indoline CD4mcs CJF-III-214 **(17)**, CJF-III-289 **(18)**, CJF-III-288 **(19)**, CJF-III-192 **(26)**, CJF-IV-047 **(28)**, DY-III-065 **(29)** and CJF-IV-046 **(30)**. The color key for each ligand is the same with that in Figure S1 and also shown in the top right corner of this Figure S3. **(B)** Comparison between two of the most potent indoline CD4mcs, DY-III-065 **(29)** and CJF-IV-046 **(30),** and two indane CD4mcs, BNM-III-170 **(1)** and CJF-III-049-S **(2)**. **(C)** Comparison of the benzyl carbamate indoline CD4mc CJF-III-192 **(26)** and its pentafluorinated counterpart CJF-IV-046 **(30)**. **(D)** Comparison of propyl carbamate indoline CD4mc CJF-III-288 **(19)** and its trifluorinated counterpart DY-III-065 **(29).**

#### Table S1. Diffraction Data and Refinement Statistics

| CD4mc dataset | BNM-III-170 **(1)** | CJF-III-049-S **(2)** | CJF-III-214 **(17)** |
|---|---|---|---|
| Beamline | APS 24ID-C | APS 24ID-E | APS 24ID-C |
| Wavelength (Å) | 0.9792 | 0.9792 | 0.9792 |
| Space group | $P2_12_12_1$ | $P2_12_12_1$ | $P2_12_12_1$ |
| Unit cell parameters a, b, c (Å) | 71.79, 120.78, 194.90 | 71.08, 121.93, 195.93 | 71.80, 120.89, 195.16 |
| α, β, γ (°) | 90, 90, 90 | 90, 90, 90 | 90, 90, 90 |
| $Z_a$[a] | 4 | 4 | 4 |
| Bragg spacings (Å)[b] | 48.73-2.04 (2.34-2.04) | 48.98-2.00 (2.26-2.00) | 48.79-2.08 (2.30-2.08) |
| Total reflections | 870028 (49953) | 474362 (23366) | 501236 (27752) |
| Unique reflections | 64404 (3788) | 69947 (3497) | 72754 (3829) |
| Completeness (%) | 94.0 (74.8) | 92.9 (66.9) | 93.5 (74.3) |
| Multiplicity | 13.5 | 6.8 | 6.9 |
| $CC_{1/2}$ (%)[c] | 99.3 (53.4) | 99.7 (59.3) | 99.8 (67.0) |
| $\langle I/\sigma(I) \rangle$ [d] | 9.1 (1.7) | 9.2 (1.6) | 9.9 (1.6) |
| $R_{merge}$[e] | 0.237 (1.965) | 0.138 (1.290) | 0.135 (1.176) |
| $R_{pim}$[f] | 0.067 (0.561) | 0.057 (0.538) | 0.055 (0.469) |
| $R_{work}$[g] | 0.2473 | 0.2631 | 0.2530 |
| $R_{free}$[h] | 0.2761 | 0.2834 | 0.2733 |
| RMS bond deviation (Å) | 0.011 | 0.013 | 0.010 |
| RMS angle deviation (º) | 1.537 | 2.088 | 1.510 |
| Average B factor (Å$^2$) | 39.50 | 42.83 | 44.53 |
| Ramachandran analysis favored/allowed (%) | 95.74/99.09 | 94.83/99.70 | 93.62/99.70 |
| PDB code | 8FLY | 8FLZ | 8FM0 |

| CD4mc dataset | CJF-III-289 **(18)** | CJF-III-288 **(19)** | CJF-III-192 **(26)** |
|---|---|---|---|
| Beamline | APS 24ID-E | APS 24ID-E | APS 24ID-C |
| Wavelength (Å) | 0.9792 | 0.9792 | 0.9792 |
| Space group | $P2_12_12_1$ | $P2_12_12_1$ | $P2_12_12_1$ |
| Unit cell parameters a, b, c (Å) | 71.22, 121.18, 194.63 | 71.46, 121.72, 195.18 | 72.36, 121.10, 195.11 |
| α, β, γ (°) | 90, 90, 90 | 90, 90, 90 | 90, 90, 90 |
| $Z_a$[a] | 4 | 4 | 4 |
| Bragg spacings (Å)[b] | 48.66-2.40 (2.73-2.40) | 48.80-2.11 (2.41-2.11) | 48.78-2.34 (2.63-2.34) |
| Total reflections | 546527 (26865) | 373721 (18355) | 596729 (30176) |
| Unique reflections | 40583 (2029) | 55435 (2771) | 49263 (2738) |
| Completeness (%) | 92.6 (62.1) | 93.3 (69.4) | 94.5 (72.2) |
| Multiplicity | 13.5 | 6.7 | 12.1 |
| $CC_{1/2}$ (%)[c] | 98.4 (34.9) | 99.6 (57.2) | 99.4 (65.7) |
| $\langle I/\sigma(I)\rangle$ [d] | 7.7 (1.6) | 8.6 | 8.5 |
| $R_{merge}$[e] | 0.558 (3.697) | 0.168 (1.298) | 0.256 (1.509) |
| $R_{pim}$[f] | 0.166 (1.046) | 0.070 (0.544) | 0.077 (0.475) |
| $R_{work}$[g] | 0.2188 | 0.2469 | 0.2369 |
| $R_{free}$[h] | 0.2622 | 0.2729 | 0.2713 |
| RMS bond deviation (Å) | 0.013 | 0.009 | 0.013 |
| RMS angle deviation (º) | 1.616 | 1.403 | 1.700 |
| Average B factor (Å$^2$) | 38.39 | 41.10 | 41.78 |
| Ramachandran analysis favored/allowed (%) | 91.79/97.57 | 95.44/99.39 | 94.53/98.78 |
| PDB code | 8FM2 | 8FM3 | 8FM7 |

| CD4mc dataset | CJF-IV-047 **(28)** | DY-III-065 **(29)** | CJF-IV-046 **(30)** |
|---|---|---|---|
| Beamline | APS 24ID-C | APS 24ID-E | APS 24ID-E |
| Wavelength (Å) | 0.9792 | 0.9792 | 0.9792 |
| Space group | $P2_12_12_1$ | $P2_12_12_1$ | $P2_12_12_1$ |
| Unit cell parameters a, b, c (Å) | 71.91, 121.26, 194.60 | 71.52, 121.66, 195.23 | 71.53, 121.79, 194.60 |
| α, β, γ (°) | 90, 90, 90 | 90, 90, 90 | 90, 90, 90 |
| $Z_a$[a] | 4 | 4 | 4 |
| Bragg spacings (Å)[b] | 48.65-2.18 (2.42-2.18) | 48.81-1.88 (2.12-1.88) | 48.65-2.47 (2.74-2.47) |
| Total reflections | 1274520 (81430) | 1050563 (49878) | 583609 (29417) |
| Unique reflections | 65466 (4092) | 88049 (4402) | 42968 (2148) |
| Completeness (%) | 94.9 (64.8) | 93.7 (62.6) | 94.3 (67.9) |
| Multiplicity | 19.5 | 11.9 | 13.6 |
| $CC_{1/2}$ (%)[c] | 99.8 (62.0) | 99.7 (56.7) | 99.6 (59.0) |
| $\langle I/\sigma(I)\rangle$ [d] | 12.0 | 11.9 | 10.3 |
| $R_{merge}$[e] | 0.216 (2.248) | 0.177 (1.666) | 0.250 (1.877) |
| $R_{pim}$[f] | 0.050 (0.515) | 0.052 (0.514) | 0.070 (0.523) |
| $R_{work}$[g] | 0.2369 | 0.2672 | 0.2172 |
| $R_{free}$[h] | 0.2611 | 0.2876 | 0.2625 |
| RMS bond deviation (Å) | 0.013 | 0.012 | 0.013 |
| RMS angle deviation (º) | 1.789 | 2.388 | 1.669 |
| Average B factor (Å$^2$) | 46.40 | 50.13 | 42.08 |
| Ramachandran analysis favored/allowed (%) | 95.14/99.39 | 95.44/99.70 | 93.01/98.48 |
| PDB code | 8FM4 | 8FM5 | 8FM8 |

[a] Za stands for number of subunits per asymmetric unit.

[b] Values in the outermost shell are given in parentheses.
[c] CC1⁄2 is the correlation coefficient of integrated intensities between randomly split two half data sets.
[d] $\langle I/\sigma(I)\rangle = \langle(\langle Ii\rangle) / \langle\sigma(\langle Ii\rangle)\rangle$
[e] Rmerge = $(\Sigma |Ii - \langle Ii \rangle|) / \Sigma |Ii|$, where Ii is the integrated intensity of a given reflection.
[f] Rpim = $(1 / (n - 1))^{1/2} \times (\Sigma |Ii - \langle Ii \rangle|) / \Sigma |Ii|$, where Ii is the integrated intensity of a given reflection.
[g] Rwork = $(\Sigma \,|\, |Fo| - |Fc| \,|) / \Sigma|Fo|$, where Fo and Fc denote observed and calculated structure factors, respectively.
[h] Rfree was calculated using 10% of data excluded from refinement.

**Table S2. H bonds and VDW contacts between CD4mcs and gp120**

| CD4mc complex | C3 or N3 substituent | Non-H atoms | H atoms | All | H bonds | VDW contacts (cutoff -0.3 Å) |
|---|---|---|---|---|---|---|
| CJF-III-049-S **(2)** | $-CH_2CH_2CHOHCH_2OH$ | 6 | 9 | 15 | 2 | 5 |
| CJF-III-214 **(17)** | $-CO_2CH_3$ | 4 | 3 | 7 | 0 | 6 |
| CJF-III-289 **(18)** | $-CO_2CH_2CH_3$ | 5 | 5 | 10 | 0 | 8 |
| CJF-III-288 **(19)** | $-CO_2CH_2CH_2CH_3$ | 6 | 7 | 13 | 0 | 8 |
| CJF-IV-047 **(28)** | $-CO_2CH_2CF_3$ | 8 | 2 | 10 | 0 | 9 |
| DY-III-065 **(29)** | $-CO_2CH_2CH_2CF_3$ | 9 | 4 | 13 | 0 | 9 |
| CJF-III-192 **(26)** | $-CO_2CH_2C_6H_5$ | 10 | 7 | 17 | 0 | 10 |
| CJF-IV-046 **(30)** | $-CO_2CH_2C_6F_5$ | 15 | 2 | 17 | 0 | 12 |

#### Summary of Neutralization Breadth Data

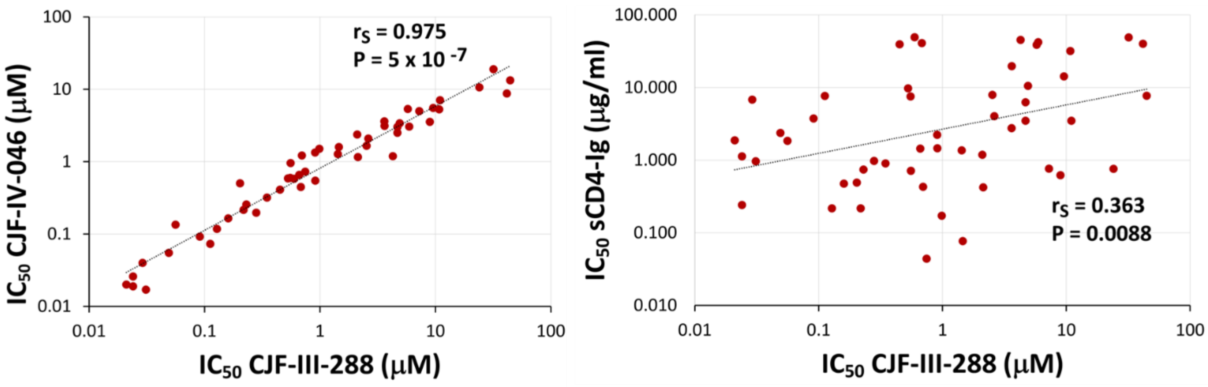

**Figure S4. Comparison of CD4mc and sCD4-Ig inhibition of global HIV-1 strains.** Correlations are shown between indoline CD4mc and sCD4-Ig inhibition of infection by HIV-1 from multiple phylogenetic clades (excluding clade AE recombinants). The IC50 values for these correlations were derived from Table S1. The Spearman rank correlation coefficients and P values are indicated.

#### Summary of neutralization data

**Table S3.** Antiviral activity of the most potent indoline CD4mcs and BNM-III-170[a]

| | BNM-III-170 | CJF-III-288 | CJF-III-192 | CJF-IV-047 | DY-III-065 | CJF-IV-046 |
|---|---|---|---|---|---|---|
| **AD8** | 3.07 ± 1.04 | 0.32 ± 0.21 | 0.15 ± 0.05 | 0.14 ± 0.05 | 0.58 ± 0.29 | 0.22 ± 0.03 |
| **JR-FL** | 25.50 ± 10.07 | 1.38 ± 0.59 | 0.57 ± 0.38 | 0.70 ± 0.30 | 1.60 ± 0.90 | 0.77 ± 0.40 |
| **BG505** | >300 | 46.67 ± 12.91 | 28.33 ± 10.41 | 46.0 ± 18.52 | 60.33 ± 42.78 | 15.67 ± 7.51 |
| | **1** | **19** | **26** | **28** | **29** | **30** |

[a]The antiviral activities of the five most potent indoline CD4mcs are compared with that of the indane CD4mc BNM-III-170. Recombinant luciferase-expressing viruses pseudotyped with the Envs of the indicated HIV-1 strains were incubated with the CD4mcs and then added to Cf2Th-CD4/CCR5 target cells. Forty-eight hours later, the level of infection in the target cells was assessed by a luciferase assay. The means and standard deviations of the IC50 values (in μM) of the CD4mcs are shown.

**Table S4.** CD4mc and sCD4-Ig inhibition of multiclade HIV-1 Env pseudovirus infection of TZM-bl cells[a]

| | | BNM-III-170 (μM) | | CJF-III-288 (μM) | | CJF-IV-046 (μM)[b] | | sCD4-Ig (mg/ml) | |
|---|---|---|---|---|---|---|---|---|---|
| **Virus ID** | **Clade** | **IC50** | **IC80** | **IC50** | **IC80** | **IC50** | **IC80** | **IC50** | **IC80** |
| 6535.3 | B | **0.221** | **0.832** | **0.031** | **0.075** | **0.017** | **0.046** | **0.964** | **8.675** |
| SC422661.8 | B | **0.239** | **1.135** | **0.029** | **0.134** | **0.040** | **0.138** | **6.830** | **66.199** |
| TRO.11 | B | **8.604** | **44.803** | **0.682** | **3.792** | **0.448** | **1.527** | **40.901** | >75 |
| RHPA4259.7 | B | **17.238** | **70.140** | **0.910** | **4.486** | **0.547** | **2.832** | **1.457** | **14.603** |
| REJO4541.67 | B | **0.130** | **0.624** | **0.024** | **0.092** | **0.019** | **0.060** | **0.242** | **3.167** |
| WITO4160.33 | B | **0.432** | **1.592** | **0.024** | **0.107** | **0.026** | **0.112** | **1.127** | **19.257** |
| WEAU_d15_410_787 | B (T/F) | **3.781** | **17.420** | **0.280** | **1.384** | **0.197** | **0.919** | **0.975** | **13.406** |
| 1054_07_TC4_1499 | B (T/F) | **0.171** | **0.776** | **0.021** | **0.073** | **0.020** | **0.104** | **1.873** | **19.359** |
| 1012_11_TC21_3257 | B (T/F) | **28.599** | **72.229** | **4.678** | **10.463** | **2.511** | **9.499** | **6.254** | **43.703** |
| 6244_13_B5_4576 | B (T/F) | **5.314** | **15.049** | **0.552** | **1.648** | **0.599** | **1.164** | **7.525** | **44.813** |
| SC05_8C11_2344 | B (T/F) | **0.612** | **2.214** | **0.049** | **0.207** | **0.055** | **0.218** | **2.368** | **24.107** |
| Du172.17 | C | >100 | >100 | **24.034** | **56.466** | **10.626** | **19.347** | **0.760** | **3.858** |
| ZM197M.PB7 | C | **1.737** | **4.782** | **0.112** | **0.326** | **0.073** | **0.222** | **7.662** | **49.338** |
| ZM233M.PB6 | C | **0.794** | **2.104** | **0.056** | **0.175** | **0.135** | **0.498** | **1.844** | **13.208** |
| ZM53M.PB12 | C | **85.652** | >100 | **3.616** | **7.983** | **3.612** | **7.697** | **2.762** | **14.744** |

| | | | | | | | | | |
|---|---|---|---|---|---|---|---|---|---|
| ZM135M.PL10a | C | **4.535** | **13.074** | **0.528** | **1.573** | **0.587** | **1.643** | **9.758** | **60.786** |
| CAP210.2.00.E8 | C | **8.083** | **28.088** | **0.556** | **2.502** | **0.958** | **3.168** | **0.714** | **3.596** |
| HIV-0013095-2.11 | C | **1.990** | **6.392** | **0.160** | **0.366** | **0.165** | **0.443** | **0.475** | **3.025** |
| HIV-16845-2.22 | C | **1.383** | **6.067** | **0.128** | **0.538** | **0.118** | **0.476** | **0.217** | **1.033** |
| Ce0393_C3 | C (T/F) | **33.121** | **82.653** | **4.907** | **13.401** | **3.401** | **7.713** | **10.512** | **62.071** |
| Ce2010_F5 | C (T/F) | >100 | >100 | **31.882** | **74.455** | **19.000** | **51.393** | **49.003** | >75 |
| Ce0682_E4 | C (T/F) | **30.774** | **87.112** | **5.931** | **16.792** | **3.049** | **11.579** | **42.155** | >75 |
| Ce703010054_2A2 | C (T/F) | >100 | >100 | **10.988** | **25.134** | **7.055** | **13.093** | **3.493** | **21.597** |
| 246F C1G | C (T/F) | **61.001** | >100 | **8.953** | **31.268** | **3.542** | **10.670** | **0.621** | **2.182** |
| ZM247v1(Rev-) | C (T/F) | **32.530** | >100 | **7.234** | **37.474** | **4.999** | **18.167** | **0.764** | **6.606** |
| CNE19 | BC | **30.043** | >100 | **2.099** | **10.554** | **2.376** | **8.186** | **1.187** | **13.824** |
| CNE21 | BC | **12.710** | **44.978** | **1.429** | **4.835** | **1.277** | **3.441** | **1.367** | **10.594** |
| CNE30 | BC | **3.413** | **15.431** | **0.218** | **0.815** | **0.216** | **0.791** | **0.217** | **1.386** |
| CNE53 | BC | **1.371** | **4.478** | **0.091** | **0.261** | **0.092** | **0.261** | **3.760** | **59.427** |
| Q23.17 | A | **38.117** | **82.150** | **9.578** | **21.074** | **5.537** | **12.488** | **14.253** | **48.858** |
| Q461.e2 | A | >100 | >100 | **3.630** | **9.328** | **3.114** | **6.904** | **19.676** | **70.812** |
| Q259.d2.17 | A | **12.638** | **34.870** | **2.622** | **6.830** | **2.085** | **3.822** | **4.042** | **23.262** |
| 3415.v1.c1 | A | **5.185** | **9.679** | **0.451** | **1.480** | **0.410** | **0.914** | **39.312** | >75 |
| 3365.v2.c2 | A | **5.335** | **11.933** | **0.745** | **1.664** | **0.726** | **1.641** | **0.044** | **0.262** |
| 191084 B7-19 | A (T/F) | **12.488** | **36.250** | **2.531** | **6.621** | **1.653** | **4.588** | **7.938** | **37.726** |
| T257-31 | CRF02_AG | **48.630** | >100 | **5.767** | **10.889** | **5.360** | **14.785** | **38.828** | >75 |
| 263-8 | CRF02_AG | **4.295** | **14.567** | **0.347** | **1.575** | **0.320** | **1.129** | **0.905** | **9.666** |
| T251-18 | CRF02_AG | **37.233** | **98.049** | **4.677** | **15.629** | **3.027** | **8.357** | **3.486** | **22.323** |
| T255-34 | CRF02_AG | **14.807** | **58.684** | **2.132** | **9.443** | **1.158** | **3.906** | **0.423** | **2.992** |
| 235-47 | CRF02_AG | **6.643** | **37.207** | **0.596** | **2.272** | **0.583** | **2.524** | **49.139** | >75 |
| 620345.c01 | CRF01_AE | >100 | >100 | >100 | >100 | **21.424** | **55.719** | >75 | >75 |
| C1080.c03 | CRF01_AE | >100 | >100 | >100 | >100 | **36.857** | **69.861** | **1.521** | **20.145** |
| R1166.c01 | CRF01_AE | >100 | >100 | >100 | >100 | **22.979** | **46.037** | >75 | >75 |

| | | | | | | | | | |
|---|---|---|---|---|---|---|---|---|---|
| C2101.c01 | CRF01_AE | >100 | >100 | >100 | >100 | **17.331** | **52.589** | **4.307** | **29.469** |
| C4118.c09 | CRF01_AE | >100 | >100 | >100 | >100 | **19.612** | **49.533** | **11.619** | **60.319** |
| BJOX009000.02.4 | CRF01_AE | >100 | >100 | >100 | >100 | **28.044** | **56.556** | **11.408** | **73.354** |
| BJOX010000.06.2 | CRF01_AE (T/F) | >100 | >100 | >100 | >100 | **20.289** | **46.769** | >75 | >75 |
| X1193_c1 | G | >100 | >100 | **41.647** | **79.588** | **8.735** | **24.353** | **40.083** | >75 |
| X1254_c3 | G | >100 | >100 | **44.510** | >100 | **13.335** | **30.204** | **7.697** | **41.286** |
| X2131_C1_B5 | G | **16.515** | **40.372** | **1.458** | **5.724** | **1.590** | **4.509** | **0.077** | **0.702** |
| X1632_S2_B10 | G | **22.999** | **79.807** | **0.986** | **2.890** | **1.504** | **4.006** | **0.172** | **0.747** |
| 3016.v5.c45 | D | **12.349** | **45.025** | **0.696** | **2.567** | **1.216** | **6.507** | **0.429** | **1.735** |
| 231965.c01 | D | **2.686** | **10.520** | **0.203** | **0.493** | **0.502** | **1.124** | **0.490** | **2.508** |
| 6405.v4.c34 | D | **6.009** | **24.275** | **4.275** | **16.396** | **1.190** | **5.810** | **45.379** | >75 |
| 3817.v2.c59 | CD | >100 | >100 | **10.760** | **99.493** | **5.272** | **20.407** | **31.876** | >75 |
| 6952.v1.c20 | CD | **4.250** | **16.513** | **0.230** | **0.819** | **0.257** | **0.946** | **0.742** | **3.920** |
| 3301.v1.c24 | AC | **8.864** | **32.110** | **0.907** | **6.390** | **1.339** | **5.990** | **2.237** | **11.426** |
| 0815.v3.c3 | ACD | **7.795** | **54.584** | **0.662** | **4.686** | **0.659** | **4.855** | **1.439** | **8.305** |
| A-MuLV | Neg. Control | >100 | >100 | >100 | >100 | **31.994** | **43.371** | >75 | >75 |

a The compound concentrations (IC50 and IC80 values) are reported that inhibit 50% and 80%, respectively, of the infection of TZM-bl cells by recombinant HIV-1 pseudotyped by the Envs from the indicated HIV-1 strains. Viruses pseudotyped with the amphotropic murine leukemia virus (A-MLV) envelope glycoprotein serve as a negative control for specificity.

b The concentrations of CJF-IV-046 highlighted in yellow are in a range that was associated with toxicity for TZM-bl cells. Therefore, the IC50 and IC80 values of CJF-IV-046 are not reliable indicators of specific antiviral activity.

#### Experimental Methods

##### Modeling

Molecular Dynamics: The structure of CD4 mimetic BNM-III-170 in complex with BG505 SOSIP.664 HIV-1 Env trimer (PDB ID: 7LO6) [1] prepared using Protein Preparation Wizard at default settings and energy-minimized using Maestro (Schrödinger Inc., 2022) [2-6]. The prepared system then subjected to solvent-explicit, all-atom molecular dynamics simulation using GPU-accelerated Desmond software (Schrödinger Inc., 2022) [5,6]. The model produced using the OPLS4 force field, and counter-ions were used to modify the system net charge. The full-atom system was immersed in a periodic TIP3P water orthorhombic box. Then molecular dynamic run was carried out for 300 ns. The recording interval was set to 1.0 ps for the trajectory and 1.0 ps for the energy. The NPT ensemble class with a temperature of 300 K and a pressure of 1.01325 bar was used. The obtained trajectory was clustered according to backbone RMSD as structural similarity metric to identify the representative conformation (centroid structure).

Molecular Docking: The target structure was taken from core monomer BG505 gp120 from the representative conformation obtained from the post MD simulation clustering. The target structure then assessed using Protein Preparation Wizard at default settings before subsequent docking calculation. Docking performed using Glide[7,8,9] with the standard precision (SP) scoring function and the number of output poses was increased to 100. The oxalamide torsional angle was restrained to 180° to keep the carbonyl groups in dipole-minimizing trans conformation.

Free energy of binding calculations: Prime MM-GBSA (Molecular Mechanics/Generalized Born Model and Solvent Accessibility) was used to estimate the ligand binding energy of each docked pose using OPLS4 force field, VSGB solvent model[9,10], and rotamer search algorithms. The protein flexibility was included to residues within 10 Å from ligands.

---

1 Jette, Claudia A., Christopher O. Barnes, Sharon M. Kirk, Bruno Melillo, Amos B. Smith, and Pamela J. Bjorkman. "Cryo-EM structures of HIV-1 trimer bound to CD4-mimetics BNM-III-170 and M48U1 adopt a CD4-bound open conformation." Nature communications 12, no. 1 (2021): 1-10.

2 *Maestro*; Schrödinger, LLC, New York, NY, 2022.

3 *Epike*; Schrödinger, LLC, New York, NY, 2022.

4 *Prime*; Schrödinger, LLC, New York, NY, 2022.

5 *Desmond*; Schrödinger, LLC, New York, NY, 2022.

6 Bowers, Kevin J., et al., Scalable algorithms for molecular dynamics simulations on on commodity clusters. ACM/IEEE CS 2006 Conf. 2006, 43-43.

7 *Glide*; Schrödinger, LLC, New York, NY, 2022.

8 Halgren, Thomas A., et al., Glide: a new approach for rapid, accurate docking and scoring. 2. Enrichment factors in database screening. J. Med. Chem. 47.7 (2004): 1750-1759.

9 Friesner, Richard A., et al., Glide: a new approach for rapid, accurate docking and scoring. 1. Method and assessment of docking accuracy. J. Med. Chem. 47.7 (2004): 1739-1749.

10 Li, Jianing, Robert Abel, Kai Zhu, Yixiang Cao, Suwen Zhao, and Richard A. Friesner. "The VSGB 2.0 model: a next generation energy model for high resolution protein structure modeling." Proteins: Structure, Function, and Bioinformatics 79, no. 10 (2011): 2794-2812.

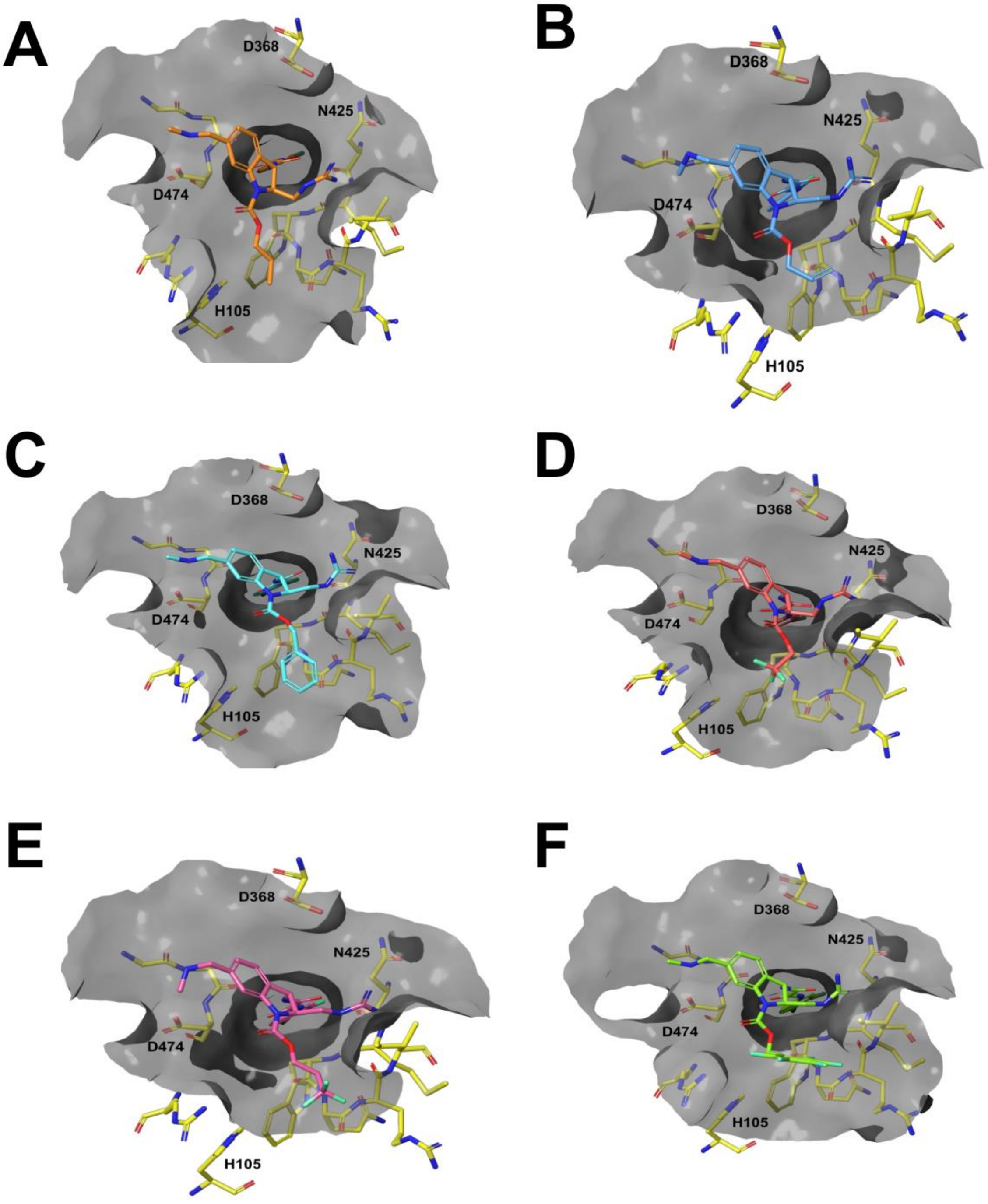

**Figure S5.** *In silico* models of indoline CD4mc's (**A**) **19**, (**B**) **23,** (**C**) **26**, (**D**) **28**, (**E**) **29**, and (**F**) **30** docked in a gp120BG505 monomer.

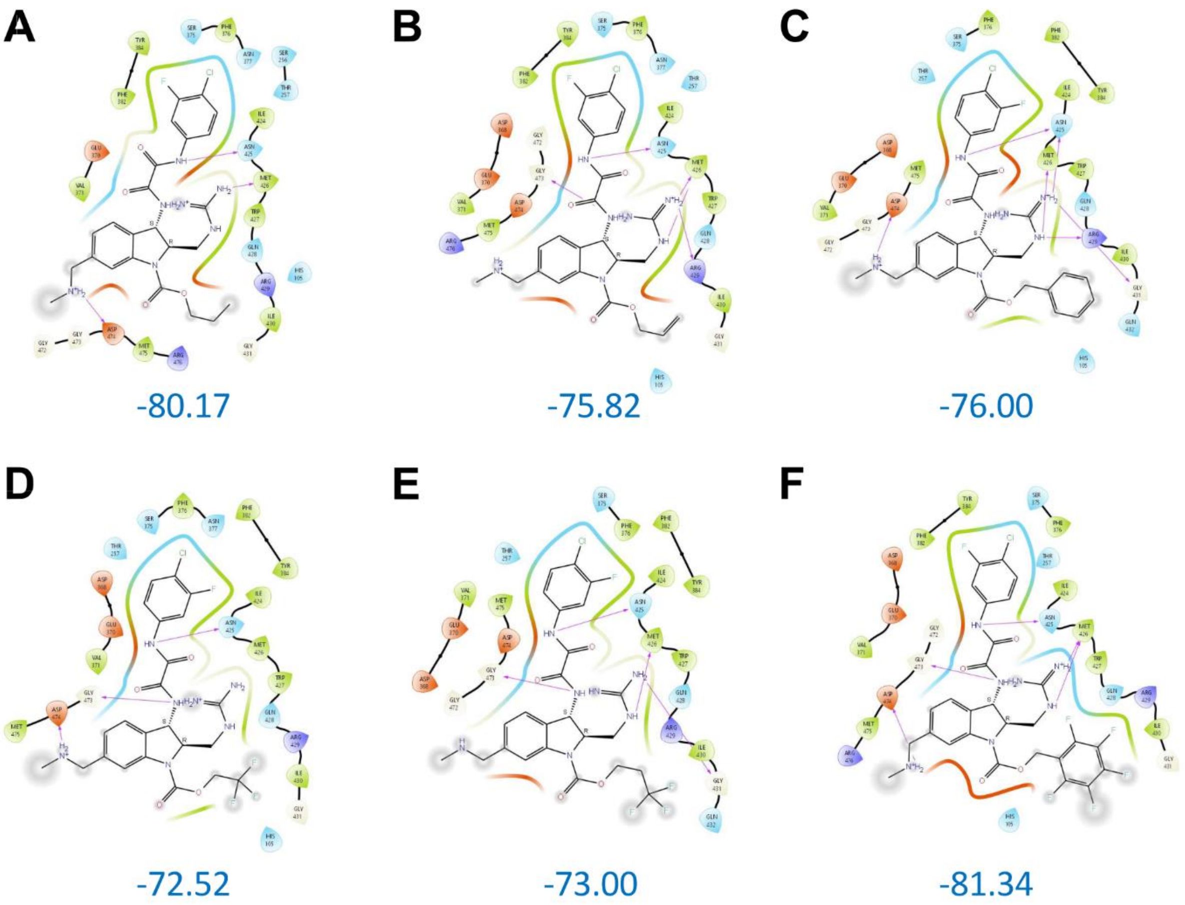

**Figure S6.** 2D representation of indoline CD4mc's interaction for (**A**) **19**, (**B**) **23,** (**C**) **26**, (**D**) **28**, (**E**) **29**, and (**F**) **30** docked in gp120$_{BG505}$ monomer. The values of Prime MM-GBSA are shown in blue (kcal/mol). The gp120 residues are colored as follows: green, nonpolar; blue, polar; indigo, basic; red, acidic.

#### Small Molecule Synthesis

##### General Information

All solvents were reagent or high-performance liquid chromatography (HPLC) grade. Anhydrous $CH_2Cl_2$ and THF were obtained from the Pure SolveTM PS-400 system under an argon atmosphere. All reagents were purchased from commercially available sources and used as received. Reactions were magnetically stirred under a nitrogen or argon atmosphere, unless otherwise noted and reactions were monitored by Thin layer chromatography (TLC) was performed on pre-coated silica gel 60 F-254 plates (40-55 micron, 230-400 mesh) and visualized by UV light. Yields refer to chromatographically and spectroscopically pure compounds. Optical rotations were measured on a JASCO P-2000 polarimeter. Proton ($^1H$) and carbon ($^{13}C$) NMR spectra were recorded on a Bruker Avance III 500-MHz spectrometer or a Bruker NEO600 600-MHz spectrometer. Chemical shifts (δ) are reported in parts per million (ppm) relative to chloroform (δ 7.26), methanol (δ 3.31), or acetone (δ 2.05) for $^1H$ NMR, and chloroform (δ 77.2) methanol (δ 49.15), or acetone (δ 29.92) for $^{13}C$ NMR. High resolution mass spectra (HRMS) were recorded at the University of Pennsylvania Mass Spectroscopy Service Center on either a VG Micromass 70/70H or VG ZAB-E spectrometer. Analytical HPLC was performed with a Waters HPLC-MS system, consisting of a 515 pump and Sunfire C18 reverse phase column (20 µL injection volume, 5 µm packing material, 4.5 × 50 mm column dimensions) with detection accomplished by a Micromass ZQ mass spectrometer and 2996 PDA detector. SFC analyses were performed with a JASCO system equipped with a PU-280-$CO_2$ plus $CO_2$ Delivery System, a CO-2060 plus Intelligent Column Thermostat/Selector, an HC-2068-01 Heater Controller, a BP-2080 plus Automatic Back Pressure Regulator, an MD-2018 plus Photodiode Array Detector (200-648 nm), and PU-2080 plus Intelligent HPLC Pumps. The purity of new compounds was judged by NMR and LCMS (>95%).

##### Synthesis of Indoline CD4mc Intermediates

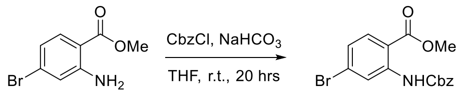

4 88% SI.1

**methyl 2-(((benzyloxy)carbonyl)amino)-4-bromobenzoate (SI.1)** To a 3-neck 1 L round-bottomed flask fitted with a 50 mL addition funnel and magnetic stirring bar was added methyl 2-amino-4-bromobenzoate (**4**) (50.4 g, 219.1 mmol, 1.0 equiv.). The atmosphere was then purged and placed under argon. THF (438.2 mL, 0.50 M) was then added at room temperature, followed by $NaHCO_3$ (55.2 g, 657.4 mmol, 3.0 equiv.) under a positive pressure of argon. Benzyl chloroformate (46.72 mL, 328.7, 1.5 equiv.) was then added to the addition funnel and added dropwise to the heterogenous solution at a drop rate of approximately 1 drop/second. After completion of the addition, the reaction was allowed to stir at room temperature overnight. Distilled water (250 mL) was then added to the reaction mixture, and the aqueous layer was extracted with $CH_2Cl_2$ (3 x 250 mL). The organic layers were then combined, washed with brine, dried over $Na_2SO_4$, and concentrated *in vacuo* to a yellow solid. The solid was then triturated with 500 mL of a 1:3 mixture of $CH_2Cl_2$:hexanes and collected via vacuum filtration to obtain a yellow solid (**SI.1**). The solvent of the filtrate was then concentrated *in vacuo* and resubjected to the same trituration conditions as described above to obtain a second crop of **SI.1** (70.1 g, 88% yield).

**$^1$H NMR** (500 MHz, $CDCl_3$) δ 10.59 (s, 1H), 8.72 (d, J = 1.9 Hz, 1H), 7.85 (d, J = 8.5 Hz, 1H), 7.45 – 7.41 (m, 2H), 7.40 – 7.36 (m, 2H), 7.36 – 7.32 (m, 1H), 7.16 (dd, J = 8.5, 2.0 Hz, 1H), 5.22 (s, 2H), 3.90 (s, 3H); **$^{13}$C NMR** (125 MHz, $CDCl_3$) δ 168.20, 153.37, 142.76, 136.10, 132.15, 129.87, 128.78, 128.53, 128.48, 125.10, 121.99, 113.40, 67.34, 52.62; **HRMS** (EI) *m/z* 363.0109 [calcd for $C_{16}H_{14}BrNO_4$ $(M)^+$ 363.0106].

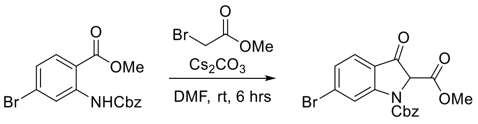

SI.1 85% 6

**1-benzyl 2-methyl 6-bromo-3-oxoindoline-1,2-dicarboxylate (6)** In an open 1 L round bottom flask with oversized magnetic stirring bar (or overhead stirring apparatus depending on scale), **SI.1** (70.1 g, 192.5 mmol, 1.0 equiv.) was dissolved in DMF (385.0 mL, 0.5 M). To this solution was added methyl bromoacetate (19.1 mL, 202.1 mmol, 1.05 equiv.) at room temperature, followed by cesium carbonate (188.1 g, 577.43 mmol, 3.0 equiv.). The heterogenous mixture was then stirred at room temperature for 6 hours, at which time UPLCMS analysis indicated consumption of starting material. The remaining solid was filtered, and the filtrate was diluted with $H_2O$ (400 mL). The mixture was then extracted with EtOAc (3 x 250 mL) and the organic layers were then combined, washed with brine (2 x 250 mL), dried over $Na_2SO_4$, and concentrated *in vacuo*. The product was precipitated by adding $Et_2O$ and collected via vacuum filtration. The crude product was purified by flash column chromatography (20% EtOAc in hexanes) to give the product **6** as an off-white solid (66.1 g, 85% yield).

**$^1$H NMR** (500 MHz, $CD_3OD$) δ 8.50 (brs, 0.72H), 8.04 (brs, 0.14H), 7.60 (d, *J* = 8.2 Hz, 1H), 7.43 – 7.35 (m, 6H), 5.40 (d, *J* = 12.0 Hz, 1H), 5.15 (d, *J* = 12.0 Hz, 1H), 3.60 (brs, 3H); **$^{13}$C NMR** (125 MHz, $CD_3OD$) δ $^{13}$C NMR (126 MHz, MeOD) δ 190.19, 179.46, 166.40, 155.46, 152.16, 136.82, 134.04, 129.85, 129.62, 128.60, 126.77, 123.35, 120.65, 69.64, 53.87, 49.77; **HRMS** (ESI) *m/z* 404.0139 [calcd for $C_{18}H_{15}BrNO_5$ $(M+H)^+$ 404.0134].

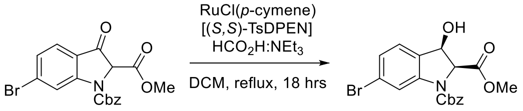

6 82% >20:1 d.r., 94% *ee* 7

**1-benzyl 2-methyl (2S,3R)-6-bromo-3-hydroxyindoline-1,2-dicarboxylate (7)** In a 2 L 2-neck round bottom flask fitted with a reflux condenser and magnetic stirring bar, **6** (66.1 g, 173.4 mmol, 1.0 equiv.) was added and capped with septa. To the flask was then added freshly distilled and sparged (30 minutes with $N_2$ balloon) $CH_2Cl_2$ (694 mL, 0.25 M), followed by addition of RuCl[(*S*,*S*)-TsDPEN](*p*-cymene) (1.10 g, 1.73 mmol, 1 mol %). $NEt_3$ (31.4 mL, 225.5 mmol, 1.3 equiv.) was then added in one portion, followed by $HCO_2H$ (20.9 mL, 555.0 mmol, 3.2 equiv.) dropwise via syringe. The reaction was then heated to reflux and stirred for 16 hours. Upon completion by TLC (30 % EtOAc in hexanes), the reaction was quenched with water (500 mL), and the aqueous layer was extracted with $CH_2Cl_2$ (3 x 300 mL). The organic layers were combined, dried over $Na_2SO_4$, and concentrated *in vacuo* to a black oil. To the oil was added a 1:1 ratio of $Et_2O$:$CH_2Cl_2$ (400 mL) which incited precipitation. The solution was heated until full dissolution was observed. The solvent was allowed to evaporate slowly, forming crystals of **6** that were collected via vacuum filtration. $^1$H NMR confirmed the d.r. to be >20:1 (57.8 g, 82% yield).

**$^1$H NMR** (500 MHz, $CD_3OD$) δ 8.07 (brs, 0.77H), 7.67 (brs, 0.16H), 7.46 – 7.33 (m, 5H), 7.26 – 7.18 (m, 2H), 5.55 (d, *J* = 9.1 Hz, 1H), 5.31 (d, *J* = 12.2 Hz, 1H), 5.10 (d, *J* = 12.3 Hz, 1H), 5.01

(d, $J = 9.1$ Hz, 1H), 3.58 (brs, 3H); **$^{13}$C NMR** (150 MHz, $CD_3OD$) δ 170.32, 153.63, 144.91, 137.29, 131.99, 129.75, 129.58, 129.38, 127.93, 127.46, 124.36, 118.70, 71.50, 68.91, 68.42, 52.65, 49.72; **HRMS** (ESI) *m/z* 406.0301 [calcd for $C_{18}H_{17}BrNO_5$ $(M+H)^+$ 406.0290]; **$[\alpha]_D^{24}$** -96.2 (*c* 0.76, MeOH).

Enantiomeric excess determined by SFC (see below):

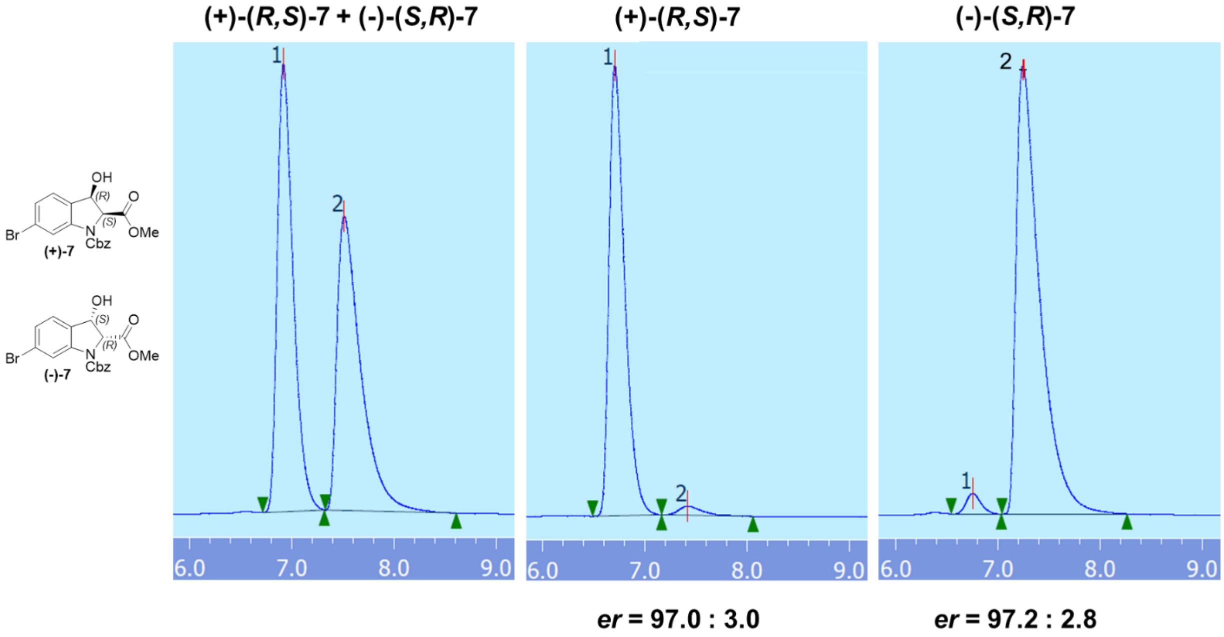

**Method:** column: Chiralpak® IA; eluent: 15% MeOH in supercritical $CO_2$; flow rate: 4 mL/min; pressure: 12 MPa. Retention times: **(+)-(*R*,*S*)-7**: 6.9 min, **(–)-(*S*,*R*)-7**: 7.5 min.

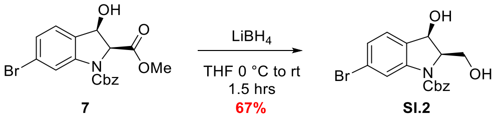

**Benzyl (2R,3R)-6-bromo-3-hydroxy-2-(hydroxymethyl)indoline-1-carboxylate (7)** In a 2 L round-bottomed flask with magnetic stirring bar, **6** (57.8 g, 142.3 mmol, 1.0 equiv.) was added and dissolved in anhydrous THF (569.1 mL, 0.25 M). The solution was then cooled to 0 °C in an ice/water bath. $LiBH_4$ powder (3.87 g, 177.9 mmol, 1.25 equiv.) was added to the reaction flask in 3 portions. The reaction was stirred at 0 °C for 30 minutes and warmed to room temperature. Upon equilibration and stirring for an additional hour, TLC (50% EtOAc in hexanes) indicated complete consumption of starting material. The reaction was cooled to 0 °C quenched with the addition of distilled water (300 mL). The reaction was then warmed to room temperature and the aqueous layer was extracted with $Et_2O$ (3 x 250 mL). The organic layer was dried over $Na_2SO_4$, filtered, and concentrated *in vacuo*. The crude product was purified by flash column chromatography (40% EtOAc/hexanes) to give **SI.2** as a white solid (36.1 g, 67%).

**$^1$H NMR** (500 MHz, $CD_3OD$) δ 7.88 (brs, 1H), 7.45 (d, $J = 7.0$ Hz, 2H), 7.39 (t, $J = 7.2$ Hz, 2H), 7.36 – 7.32 (m, 1H), 7.21 (d, $J = 7.9$ Hz, 1H), 7.17 (dd, $J = 7.9$, 1.7 Hz, 1H), 5.52 (d, $J = 8.8$ Hz, 1H), 5.32 – 5.25 (m, 2H), 4.45 (ddd, $J = 8.4$, 4.6, 3.2 Hz, 1H), 4.00 (dd, $J = 11.8$, 4.7 Hz, 1H), 3.96 (dd, $J = 11.8$, 3.3 Hz, 1H); **$^{13}$C NMR** (125 MHz, $CD_3OD$) δ 154.60, 137.57, 134.44, 129.84, 129.59,

129.52, 127.29, 127.04, 123.46, 119.43, 72.38, 68.96, 66.18, 60.72, 49.78; **HRMS** (ESI) *m/z* 400.0160 [calcd for $C_{17}H_{16}BrNO_4Na$ $(M+Na)^+$ 400.0160]; **[α]$_D$$^{23}$** -22.7 (*c* 1.00, MeOH).

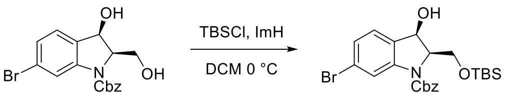

SI.2 93% SI.3

**Benzyl (2R,3R)-6-bromo-2-(((tert-butyldimethylsilyl)oxy)methyl)-3-hydroxyindoline-1-carboxylate (SI.3)** To a suspension of **SI.2** (36.1 g, 95.4 mmol, 1.0 equiv.) in $CH_2Cl_2$ (191 mL, 0.5 M) at 0 ºC was added imidazole (13.0 g, 191.0 mmol, 2.0 equiv.) in one portion and stirred for 10 min. To this mixture was added a solution of tert-butyldimethylsilyl chloride (15.8 g, 105.0 mmol, 1.1 equiv.) in $CH_2Cl_2$ (105 mL, 1.0 M) dropwise over 30 min via dropping funnel. The reaction mixture was stirred at 0 ºC for 30 min. Upon consumption of starting material based on TLC, the resulting mixture was treated with $H_2O$ (150 mL). The aqueous phase was then extracted with $CH_2Cl_2$ (3 x 100 mL), the organic layers were dried over $Na_2SO_4$, filtered, and concentrated *in vacuo*. The resulting oil was further purified by flash column chromatography (10% EtOAc/hexanes) to give **SI.3** as a clear oil (43.7 g, 93%).

**$^1$H NMR** (500 MHz, $CD_3OD$) δ 7.83 (s, 1H), 7.45 (d, *J* = 6.8 Hz, 2H), 7.41 – 7.33 (m, 3H), 7.19 – 7.15 (d, *J* = 7.9 Hz, 1H), 7.13 (d, *J* = 8.0 Hz, 1H), 5.48 (d, *J* = 8.8 Hz, 1H), 5.32 (d, *J* = 12.0 Hz, 1H), 5.22 (d, *J* = 12.2 Hz, 1H), 4.47 (dtd, *J* = 8.8, 2.9, 1.3 Hz, 1H), 4.06 (d, *J* = 2.8 Hz, 2H), 0.63 (s, 9H), -0.11 (s, 3H), -0.22 (s, 3H); **$^{13}$C NMR** (125 MHz, $CD_3OD$) δ 154.32, 144.68, 137.52, 135.31, 129.88, 129.77, 129.69, 127.13, 126.31, 123.13, 119.10, 72.60, 68.85, 66.17, 61.72, 26.11, 18.79, -5.46, -5.49; **HRMS** (ESI) *m/z* 492.1211 [calcd for $C_{23}H_{31}BrNO_4Si$ $(M+H)^+$ 492.1206]; **[α]$_D$$^{23}$** +48.3 (*c* 1.00, MeOH).

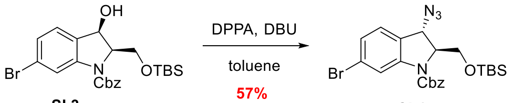

SI.3 57% SI.4

**Benzyl (2S,3S)-3-azido-6-bromo-2-(((tert-butyldimethylsilyl)oxy)methyl)indoline-1-carboxylate (SI.4)** To a solution of **SI.3** (43.7 g, 88.7 mmol, 1.0 equiv.) in toluene (296.0 mL, 0.3 M) was added diphenylphosphoryl azide (28.3 mL, 131.3 mmol, 1.48 equiv.) dropwise for 10 min, followed by dropwise addition of DBU (19.1 mL, 127.8 mmol, 1.44 equiv.) over 10 min. A cloudy mixture was observed upon addition, which was stirred at room temperature overnight. The resulting biphasic red mixture filtered through a pad of silica and washed with $Et_2O$ (500 mL). The filtrate was concentrated *in vacuo*, and the resulting oil was further purified by flash column chromatography (2% $Et_2O$/hexanes) to give **SI.4** as a clear oil (26.2 g, 57%).

**$^1$H NMR** (500 MHz, $CD_3OD$) δ 8.04 (brs, 0.72H), 7.66 (brs, 0.20H), 7.46 (d, *J* = 6.8 Hz, 2H), 7.42 – 7.34 (m, 3H), 7.32 (d, *J* = 8.0 Hz, 1H), 7.23 (d, *J* = 8.0, Hz, 1H), 5.38 (brs, 1H), 5.17 (brs, 1H), 4.89 (s, 1H), 4.30 (s, 1H), 3.85 (d, *J* = 10.1 Hz, 1H), 3.68 (brs, 1H), 0.67 (s, 9H), -0.11 (brs, 3H), -0.20 (brs, 3H); **$^{13}$C NMR** (150 MHz, $CD_3OD$) δ 153.58, 145.89, 137.32, 129.95, 129.86, 129.83, 128.77, 127.96, 127.36, 125.04, 119.53, 69.31, 69.02, 63.90, 60.27, 26.11, 18.87, -5.41, -5.51; **HRMS** (ESI) *m/z* 517.1282 [calcd for $C_{23}H_{30}BrN_4O_3Si$ $(M+H)^+$ 517.1271]; **[α]$_D$$^{23}$** +49.0 (*c* 1.00, MeOH).

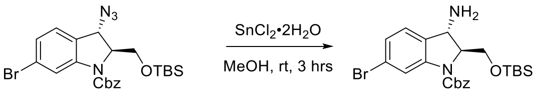

SI.4 72% 8

**Benzyl (2S,3S)-3-amino-6-bromo-2-(((tert-butyldimethylsilyl)oxy)methyl)indoline-1-carboxylate (8)** To a solution **SI.4** (26.2 g, 50.6 mmol, 1.0 equiv.) in MeOH (92.1 mL, 0.55 M) at room temperature, was added $SnCl_2$•$2H_2O$ (17.1 g, 75.9 mmol, 1.5 equiv.) in a solution of MeOH (152 mL, 0.5 M). The solution was stirred for 3 hours at room temperature. Upon completion by TLC, 1M NaOH (100 mL) was added and the resulting precipitate was filtered through a thick pad of Celite ®. The Celite ® pad was washed with copious MeOH (200 mL) and the filtrate was concentrated *in vacuo*. The resulting oil was further purified by flash column chromatography (5% MeOH/$CH_2Cl_2$) to give **8** as a clear oil (17.9 g, 72%).

**$^1$H NMR** (600 MHz, $CD_3OD$) δ 7.99 (brs, 0.70H), 7.62 (s, 0.21H), 7.46 (d, $J$ = 6.9 Hz, 2H), 7.41 – 7.33 (m, 3H), 7.27 – 7.24 (m, 1H), 7.14 (d, $J$ = 7.6 Hz, 1H), 5.35 (d, $J$ = 11.6 Hz, 1H), 5.18 (brs, 1H), 4.25 (d, $J$ = 1.9 Hz, 1H), 4.14 (td, $J$ = 3.4, 1.9 Hz, 1H), 4.04 – 3.71 (br m, 2H), 0.65 (s, 9H), -0.09 (brs, 3H), -0.20 (brs, 3H); **$^{13}$C NMR** (150 MHz, $CD_3OD$) δ 154.20, 145.69, 137.51, 130.87, 129.88, 129.80, 129.70, 127.07, 126.24, 123.22, 119.23, 71.74, 68.73, 64.49, 55.61, 26.13, 18.88, -5.40, -5.46; **HRMS** (ESI) $m/z$ 474.1106 [calcd for $C_{23}H_{29}BrNO_3Si$ $(M+H-NH_3)^+$ 474.1100]; **$[\alpha]_D^{23}$** +16.0 ($c$ 1.00, MeOH).

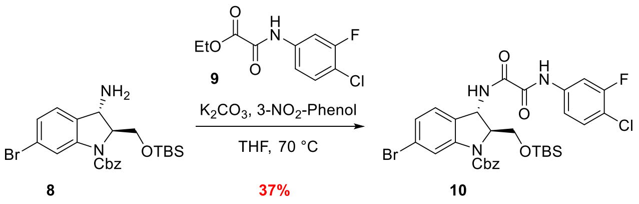

**Benzyl (2S,3S)-6-bromo-2-(((tert-butyldimethylsilyl)oxy)methyl)-3-(2-((4-chloro-3-fluoro phenyl)amino)-2-oxoacetamido)indoline-1-carboxylate (10)** A mixture of **8** (17.9 g, 36.4 mmol, 1.0 equiv.), **9** (11.2 g, 45.5 mmol, 1.25 equiv.), 3-nitrophenol (1.00 g, 7.28 mmol, 0.2 equiv.), and $K_2CO_3$ (1.01 g, 7.28 mmol, 0.2 equiv.) in THF (72.8 mL, 0.5 M) was stirred at reflux for 48 h. The resulting suspension was allowed to cool to room temperature and then treated with 10 wt% $K_2CO_3$ (50 mL) and stirred at room temperature for 1 hr. The mixture was then diluted with EtOAc (100 mL). The insoluble material was removed via vacuum filtration, and the solid was rinsed with EtOAc (2 x 100 mL). After, the filtrates were combined, the organic layer was separated and washed with 10 wt% $K_2CO_3$ (50 mL). The aqueous layers were combined and extracted with EtOAc (2 x 50 mL). The organic layers were combined and washed with brine, dried over $Na_2SO_4$, filtered, and concentrated *in vacuo* to give crude **10** as a brown oil. The resulting oil was further purified by flash column chromatography (20% EtOAc/hexanes) to give **10** as a yellow solid (9.3 g, 37%).

**$^1$H NMR** (600 MHz, $CD_3OD$) δ 8.01 (s, 1H), 7.81 (dd, $J$ = 11.4, 2.4 Hz, 1H), 7.48 – 7.44 (m, 3H), 7.42 – 7.33 (m, 4H), 7.21 (d, $J$ = 8.0 Hz, 1H), 7.14 (d, $J$ = 7.6 Hz, 1H), 5.41 (d, $J$ = 2.4 Hz, 1H), 5.36 (brs, 1H), 5.16 (brs, 1H), 4.37 – 4.35 (m, 1H), 4.08 – 3.90 (br m, 2H), 0.65 (s, 9H), -0.09 (brs, 3H), -0.21 (brs, 3H); **$^{13}$C NMR** (150 MHz, $CDCl_3$) δ 161.08, 159.79, 159.24 (d, $J_{CF}$ = 246.0 Hz), 139.18 (d, $J_{CF}$ = 9.4 Hz), 137.42, 131.79, 129.90, 129.85, 129.75, 127.71, 127.30, 124.01, 119.32, 118.20 (d, $J_{CF}$ = 3.5 Hz), 117.29 (d, $J_{CF}$ = 18.1 Hz), 109.89 (d, $J_{CF}$ = 26.1 Hz), 69.31, 68.84, 64.72, 53.89, 26.10, 18.85, -5.37, -5.44; **HRMS** (ESI) $m/z$ 690.1206 [calcd for $C_{31}H_{35}BrClFN_3O_5Si$ $(M+H)^+$ 690.1202]; **$[\alpha]_D^{24}$** +87.4 ($c$ 0.97, MeOH).

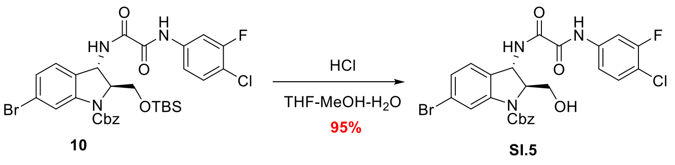

**Benzyl (2S,3S)-6-bromo-2-(((tert-butyldimethylsilyl)oxy)methyl)-3-(2-((4-chloro-3-fluoro phenyl)amino)-2-oxoacetamido)indoline-1-carboxylate compound with benzyl (2S,3S)-6-bromo-3-(2-((4-chloro-3-fluorophenyl)amino)-2-oxoacetamido)-2-(hydroxymethyl)indoline-1-carboxylate (SI.5)** Compound **10** (9.3 g, 13.5 mmol, 1.0 equiv.) was dissolved in THF (40.4 mL), and the resulting solution was diluted with MeOH (13.5 mL) and H2O (13.5 mL), to which conc. HCl (4.86 mL, 0.36 mL/mmol) was added dropwise. The reaction mixture was stirred at room temperature overnight. The mixture was then treated with sat. $NaHCO_3$ (50 mL) and $Et_2O$ (100 mL). The aqueous layer was further extracted with $Et_2O$ (2 x 100 mL). The organic layers were combined, dried over $Na_2SO_4$, filtered, and concentrated *in vacuo* to give crude **SI.5** as an off-white powder, which could be used in the next step without further purification (7.37 g, 95% yield).

**$^1$H NMR** (500 MHz, $CD_3OD$) δ 8.02 (s, 1H), 7.82 (dd, *J* = 11.4, 2.3 Hz, 1H), 7.49 – 7.44 (m, 3H), 7.42 – 7.37 (m, 2H), 7.36 – 7.33 (m, 1H), 7.24 (d, *J* = 8.1 Hz, 1H), 7.18 (d, *J* = 7.9 Hz, 1H), 5.42 (d, *J* = 2.7 Hz, 1H), 5.33 (d, *J* = 11.9 Hz, 1H), 5.28 (brs, 1H), 4.38 – 4.36 (m, 1H), 3.87 – 3.73 (m, 1H); **$^{13}$C NMR** (150 MHz, $CD_3OD$) δ 161.13, 159.84, 159.24 (d, $J_{CF}$ = 244.9 Hz), 139.25 (d, $J_{CF}$ = 9.4 Hz), 137.48, 131.80, 130.88, 129.86, 129.63, 129.57, 128.01, 127.45, 124.11, 118.20 (d, $J_{CF}$ = 3.5 Hz), 117.27 (d, $J_{CF}$ = 18.1 Hz), 109.85 (d, $J_{CF}$ = 26.1 Hz), 69.43, 68.95, 63.09, 49.72; **HRMS** (ESI) *m/z* 576.0348 [calcd for $C_{25}H_{21}BrClFN_3O_5$ $(M+H)^+$ 576.0337]; **$[\alpha]_D^{24}$** +91.7 (*c* 0.22, MeOH).

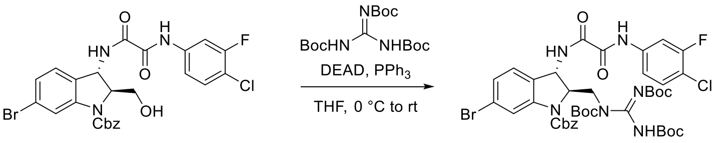

SI.5 65% 11

**Benzyl (2R,3S)-6-bromo-3-(2-((4-chloro-3-fluorophenyl)amino)-2-oxoacetamido)-2-(((Z)-1,2,3-tris(tert-butoxycarbonyl)guanidino)methyl)indoline-1-carboxylate (11)** In a 1 L round-bottomed flask with magnetic stirring bar, a mixture of **SI.5** (7.37 g, 12.8 mmol, 1.0 equiv.), $PPh_3$ (5.36 g, 20.4 mmol, 1.6 equiv.), and N,N′,N″-tri-Boc-guanidine (16.1 g, 44.7 mmol, 3.5 equiv.) in anhydrous THF (320 mL, 0.04 M) was stirred at room temperature until a well-dispersed suspension had formed. This mixture was cooled at 0 ºC and treated with diethyl azodicarboxylate (3.01 mL, 19.2 mmol, 1.5 equiv.) dropwise at such a rate that each drop was only added after the color change resulting from the previous drop had dissipated. After the addition was completed, the reaction was allowed to warm to room temperature and stir for 18 hr. TLC (30% EtOAc/hexanes) indicated complete consumption of starting material at this time. The reaction was then quenched with brine (100 mL) and the aqueous layer was extracted with EtOAc (3 x 100 mL). The organic layers were combined, dried over $Na_2SO_4$, and concentrated *in vacuo* to give a white solid which was purified by flash column chromatography (20% EtOAc/hexanes) to give a mixture of **11** and N,N′,N″-tri-Boc-guanidine. The residue was treated with $Et_2O$ (20 mL) and the insoluble material was removed via vacuum filtration. The filtrate was concentrated *in vacuo* to give **11** as an off-white amorphous solid (7.63 g, 65%).

**$^1$H NMR** (500 MHz, $CD_3OD$) δ 7.99 (brs, 1H), 7.81 (dd, *J* = 11.4, 2.3 Hz, 1H), 7.50 (d, *J* = 7.1 Hz, 2H), 7.48 – 7.41 (m, 2H), 7.40 – 7.29 (m, 3H), 7.27 (d, *J* = 8.0 Hz, 1H), 7.20 (dd, *J* = 8.0, 1.8 Hz, 1H), 5.37 – 5.25 (br m, 3H), 4.72 (s, 1H), 4.15 (dd, *J* = 14.2, 6.3 Hz, 1H), 4.02 (s, 1H), 1.54 – 1.42 (m, 27H); **$^{13}$C NMR** (150 MHz, $CD_3OD$) δ 161.12, 159.70, 159.24 (d, $J_{CF}$ = 246.0 Hz), 155.17, 151.51, 139.24, 139.18, 137.54, 131.81, 130.01, 129.79, 129.72, 129.54, 128.58, 127.79, 124.34, 118.18 (d, $J_{CF}$ = 3.5 Hz), 117.28 (d, $J_{CF}$ = 18.1 Hz), 109.84 (d, $J_{CF}$ = 26.1 Hz), 85.76, 83.72, 67.04, 28.62, 28.60, 28.41; **HRMS** (ESI) *m/z* 917.2286 [calcd for $C_{41}H_{48}BrClFN_6O_{10}$ $(M+H)^+$ 917.2288]; **$[\alpha]_D^{24}$** +54.9 (*c* 0.77, MeOH).

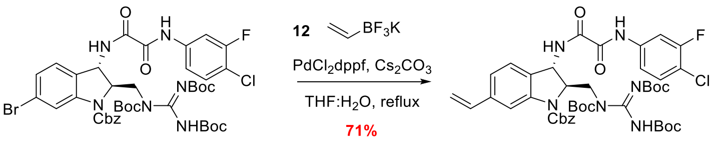

**11** → **SI.6**

**Benzyl (2R,3S)-3-(2-((4-chloro-3-fluorophenyl)amino)-2-oxoacetamido)-2-(((Z)-1,2,3-tris (tert-butoxycarbonyl)guanidino)methyl)-6-vinylindoline-1-carboxylate (14)** A flame dried flask equipped with magnetic stirring bar was charged with **11** (7.63 g, 8.31 mmol, 1.0 equiv.), potassium vinyltrifluoroborate **12** (1.14 g, 9.97 mmol, 1.2 equiv.), [1,1'-Bis(diphenyl-phosphino)ferrocene]palladium(II) dichloride (339 mg, 0.42 mmol, 5 mol%) and cesium carbonate (8.12 g, 24.9 mmol, 3.0 equiv.). The flask was evacuated and backfilled with $N_2$ atmosphere 3 times before the addition of freshly distilled THF (66.5 mL) and distilled $H_2O$ (16.7 mL). The reaction flask was then evacuated and backfilled 4 times before heating to reflux. The reaction mixture was stirred at reflux for 18 hr, at which time UPLCMS analysis indicated full consumption of starting material. The reaction was then cooled to room temperature, diluted with $Et_2O$ and $H_2O$ (100 mL each). The aqueous phase was extracted with $Et_2O$ (2 x 100 mL), and the organic layers were combined, dried over $Na_2SO_4$, filtered, and concentrated *in vacuo* to yield crude **SI.6**. The crude solid was further purified by flash column chromatography (20% EtOAc/hexanes) to give **SI.6** as a white amorphous solid (5.11 mg, 71%).

**$^1$H NMR** (500 MHz, $CD_3OD$) δ 7.93 (s, 1H), 7.81 (dd, *J* = 11.4, 2.3 Hz, 1H), 7.51 (d, *J* = 7.5 Hz, 2H), 7.46 (dd, *J* = 9.3, 2.1 Hz, 1H), 7.42 (d, *J* = 8.2 Hz, 1H), 7.41 – 7.33 (m, 1H), 7.31 (d, *J* = 7.8 Hz, 1H), 7.12 (brs, 1H), 6.71 (brs, 1H), 5.64 (brs, 1H), 5.37 – 5.19 (br m, 2H), 4.74 (s, 1H), 4.16 (dd, *J* = 14.1, 6.4 Hz, 1H), 3.97 (dd, *J* = 14.2, 7.3 Hz, 1H), 1.48 – 1.43 (m, 27H); **$^{13}$C NMR** (150 MHz, $CD_3OD$) δ 161.09, 159.76, 159.24 (d, $J_{CF}$ = 246.0 Hz), 155.26, 140.98, 139.21 (d, $J_{CF}$ = 9.4 Hz), 138.20, 132.55, 131.81, 130.02, 129.78, 129.72, 129.51, 127.26, 123.63, 118.18 (d, $J_{CF}$ = 3.5 Hz), 117.28 (d, $J_{CF}$ = 18.1 Hz), 114.98, 109.86 (d, $J_{CF}$ = 26.1 Hz), 85.75, 83.69, 66.91, 28.61, 28.42; **HRMS** (ESI) *m/z* 865.3354 [calcd for $C_{43}H_{51}ClFN_6O_{10}$ $(M+H)^+$ 865.3339]; **$[\alpha]_D^{24}$** +68.4 (*c* 0.60, MeOH).

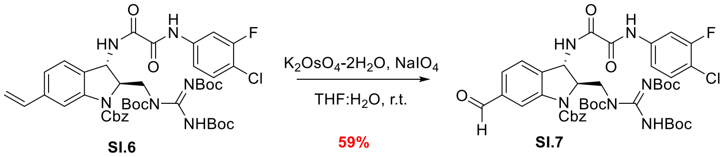

**SI.6** → **SI.7**

**Benzyl (2R,3S)-3-(2-((4-chloro-3-fluorophenyl)amino)-2-oxoacetamido)-6-formyl-2-(((Z)p-1,2,3-tris(tert-butoxycarbonyl)guanidino)methyl)indoline-1-carboxylate (SI.7)** To a solution of **SI.6** (5.11 g, 8.82 mmol, 1.0 equiv.) in a 4:1 mixture of THF/$H_2O$ (44.1 mL, 0.2 M), was added 2,6-lutidine (4.09 mL, 35.3 mmol, 4.0 equiv.) dropwise via syringe, followed by potassium osmate dihydrate (65 mg, 0.18 mmol, 2 mol %) in one portion, and sodium periodate (5.66 g, 26.5 mmol, 3.0 equiv.) in one portion. The cloudy white reaction slurry was stirred for 2 hr at room temperature until completion via TLC. The reaction was quenched by the addition of sat. $NaHCO_3$ (50 mL), diluted with $Et_2O$ (50 mL). The aqueous phase was extracted with $Et_2O$ (2 x 50 mL), and the organic layers were combined, dried over $Na_2SO_4$, filtered, and concentrated *in vacuo* to yield crude **SI.7**. The crude solid was further purified by flash column chromatography (25% EtOAc/hexanes) to give **SI.7** as a white amorphous solid (4.51 g, 59%).

**$^1$H NMR** (500 MHz, $(CD_3)_2CO$) δ 10.52 (brs, 0.5H), 10.12 (s, 0.5H), 10.02 (brs, 1H), 9.17 (d, $J$ = 8.3 Hz, 1H), 8.31 (brs, 1H), 7.98 – 7.93 (m, 1H), 7.71 – 7.69 (m, 1H), 7.62 – 7.57 (m, 4H), 7.51 (t, $J$ = 8.6 Hz, 1H), 7.40 (t, $J$ = 7.4 Hz, 2H), 7.35 (t, $J$ = 7.3 Hz, 1H), 5.69 (brs, 1H), 5.37 (d, $J$ = 12.1 Hz, 1H), 5.33 – 5.28 (br m, 1H), 4.87 (brs, 1H), 4.27 (dd, $J$ = 13.9, 5.2 Hz, 1H), 4.23 (brs, 1H), 1.50 – 1.39 (m, 27H); **$^{13}$C NMR** (150 MHz, $(CD_3)_2CO$) δ 192.65, 170.98, 160.09, 159.07, 158.59 (d, $J_{CF}$ = 246.0 Hz),154.23, 139.03 (d, $J_{CF}$ = 10.2 Hz), 137.33, 131.63, 129.41, 129.08, 126.41, 117.88 (d, $J_{CF}$ = 3.5 Hz), 116.32 (d, $J_{CF}$ = 17.7 Hz), 109.24 (d, $J_{CF}$ = 25.2 Hz), 84.55, 66.11, 60.61, 48.53, 30.43, 28.36, 20.90, 14.58; **HRMS** (ESI) *m/z* 867.3136 [calcd for $C_{42}H_{49}ClFN_6O_{11}$ $(M+H)^+$ 867.3132]; **$[\alpha]_D^{23}$** +83.48 ($c$ 1.00, MeOH).

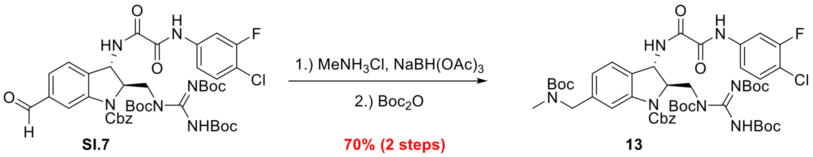

**Benzyl (2R,3S)-6-(((tert-butoxycarbonyl)(methyl)amino)methyl)-3-(2-((4-chloro-3-fluoro phenyl)amino)-2-oxoacetamido)-2-(((Z)-1,2,3-tris(tert-butoxycarbonyl)guanidino)methyl)-indoline-1-carboxylate (13)** To a stirring solution of **SI.7** (4.51 g, 5.20 mmol, 1.0 equiv.) in a 1:1 mixture of EtOH/DCE (52.0 mL, 0.1 M) at 0 ºC was added methylamine hydrochloride (1.76 g, 26.0 mmol, 5.0 equiv.) in one portion. The reaction was stirred for 10 minutes at 0 ºC before the addition of sodium triacetoxyborohydride (2.76 g, 13.0 mmol, 2.5 equiv.) in one portion. The ice bath was then removed, and the reaction mixture was allowed to warm to room temperature. The reaction was allowed to stir overnight, at which time complete consumption of **SI.7** was observed by TLC and confirmed by UPLCMS. The reaction was then diluted with $CH_2Cl_2$ (50 mL), quenched with the slow addition of $H_2O$ (50 mL). The aqueous phase was basified to a pH of 12 with aq. 1 N NaOH solution and extracted with $CH_2Cl_2$ (3 x 50 mL). The organic layers were combined, dried over $Na_2SO_4$, and concentrated *in vacuo* to give an oil. This was dissolved in $CH_2Cl_2$ (10.4 mL, 0.5M) and cooled to 0 °C, to which $Boc_2O$ (1.43 mL, 6.24 mmol, 1.2 equiv) was added dropwise via syringe. The reaction mixture was warmed to room temperature and stirred for 1 hour. The solvent was then concentrated *in vacuo* to give crude **13** as an oil, which was purified by flash column chromatography (20% EtOAc/hexanes) to give the product **13** as an amorphous solid (3.58 g, 70% over 2 steps).

**$^1$H NMR** (500 MHz, $CD_3OD$) δ 7.81 (dd, $J$ = 11.4, 2.3 Hz, 1H), 7.79 (brs, 1H), 7.49 (d, $J$ = 7.1 Hz, 2H), 7.46 (dd, $J$ = 9.1, 2.3 Hz, 1H), 7.44 – 7.30 (m, 5H), 6.96 (dd, $J$ = 7.7, 1.5 Hz, 1H), 5.36 – 5.24 (m, 3H), 4.75 – 4.71 (br m, 1H), 4.42 (brs, 2H), 4.15 (dd, $J$ = 14.1, 6.0 Hz, 1H), 3.97 (dd, $J$ = 14.1, 7.6 Hz, 1H), 2.80 (brs, 3H), 1.49 – 1.44 (m, 36H); **$^{13}$C NMR** (150 MHz, $CD_3OD$) δ 161.07, 159.77, 159.24 (d, $J_{CF}$ = 246.0 Hz), 155.24, 139.22 (d, $J_{CF}$ = 9.4 Hz), 131.81, 129.74, 129.47, 124.17, 118.17 (d, $J_{CF}$ = 3.5 Hz), 117.27 (d, $J_{CF}$ = 18.1 Hz), 109.85 (d, $J_{CF}$ = 26.1 Hz), 85.71, 83.71, 73.41, 66.95, 49.72, 28.88, 28.63, 28.44, 22.07; **HRMS** (ESI) *m/z* 982.4113 [calcd for $C_{48}H_{62}ClFN_7O_{12}$ $(M+H)^+$ 982.4129]; **$[\alpha]_D^{24}$** +57.0 ($c$ 0.85, MeOH).

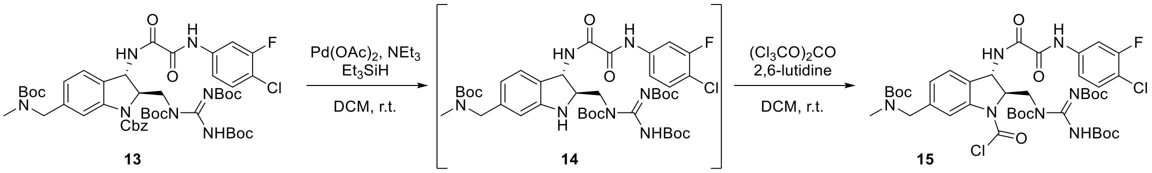

**Carbamoyl Chloride (15)** Compound **13** (500 mg, 0.51 mmol, 1.0 equiv.) was dissolved in $CH_2Cl_2$ (5.09 mL, 0.1 M) and cooled to 0 ºC under argon. In a separate flask, $Pd(OAc)_2$ (8.0 mg, 0.036 mmol, 7 mol %) and $NEt_3$ (17.7 µL, 0.127 mmol, 0.25 equiv.) were dissolved in $CH_2Cl_2$ (1.02 mL). $Et_3SiH$ (139.8 µL, 0.88 mmol, 1.72 equiv.) was then added in one portion to the mixture. Upon turning black, the solution was taken up via syringe and added to the solution of **13** dropwise. The reaction was allowed to warm to r.t. and stirred for 4 hours, at which time UPLCMS indicated complete consumption of starting material. Upon completion, excess $NEt_3$ (1 mL) was added and the solution was filtered through a pad of Celite ®. This was washed with $CH_2Cl_2$ (5 mL) and concentrated *in vacuo*. The crude material was then immediately passed through a plug of silica pre-saturated with 20% EtOAc/hexanes with 1% $NEt_3$ and washed with the same mixture (50 mL) until elution of the desired product **14** was observed by TLC (351.9 mg).

This material was then redissolved in $CH_2Cl_2$ (2.07 mL), followed by addition of 2,6-lutidine (50.5 µL, 0.436 mmol, 1.05 equiv.). The mixture was then cooled to 0 ºC, and a 0.2M stock solution of triphosgene in $CH_2Cl_2$ (0.73 mL, 0.14 mmol, 0.35 equiv.) was added dropwise. The reaction was allowed to stir for 2 hours, at which time UPLCMS analysis indicated consumption of **14**. The reaction was quenched with sat. aq. $NH_4Cl$ (2 mL), and the aqueous layer was extracted 3 x 2 mL $CH_2Cl_2$. The organic layers were separated, dried with $Na_2SO_4$, filtered and concentrated *in vacuo*. The crude carbamoyl chloride **15** was then purified by flash column chromatography (20% EtOAc/hexanes) to give **15** as a red amorphous solid (374.4 mg, 81% over 2 steps).

**$^1$H NMR** (600 MHz, $(CD_3)_2CO$) δ 10.27 (s, 0.5H), 10.13 (s, 0.5H), 9.25 (d, $J$ = 8.2 Hz, 1H), 8.02 – 7.92 (m, 1H), 7.82 (brs, 1H), 7.70 (dt, $J$ = 8.9, 1.3 Hz, 1H), 7.51 (t, $J$ = 8.6 Hz, 1H), 7.47 (brs, 1H), 7.13 (brs, 1H), 5.56 (brs, 1H), 5.02 – 4.98 (m, 1H), 4.55 – 4.42 (m, 2H), 4.30 (dd, $J$ = 14.2, 5.2 Hz, 1H), 4.12 – 4.04 (br m, 1H), 2.79 (s, 3H, obscured by residual $H_2O$), 1.51 (brs, 9H), 1.48 (brs, 9H), 1.46 – 1.45 (br m, 18H); **$^{13}$C NMR** (150 MHz, $(CD_3)_2CO$) δ 160.06, 159.06, 158.59 (d, $J_{CF}$ = 244.9 Hz), 154.13, 145.68, 142.86, 141.79, 139.03 (d, $J_{CF}$ = 9.5 Hz), 131.64, 125.89, 117.90 (d, $J_{CF}$ = 3.5 Hz), 116.34 (d, $J_{CF}$ = 17.7 Hz), 109.24 (d, $J_{CF}$ = 25.2 Hz), 84.85, 69.38, 52.88, 52.78, 48.40, 30.42, 28.66, 28.35, 28.25, 23.36, 22.28, 22.01, 14.42; **HRMS** (ESI) *m/z* 910.3319 [calcd for $C_{41}H_{55}Cl_2FN_7O_{11}$ $(M+H)^+$ 910.3321]; **$[\alpha]_D^{24}$** +71.1 (*c* 0.52, MeOH).

#### General Procedure for the Synthesis of Final Compounds

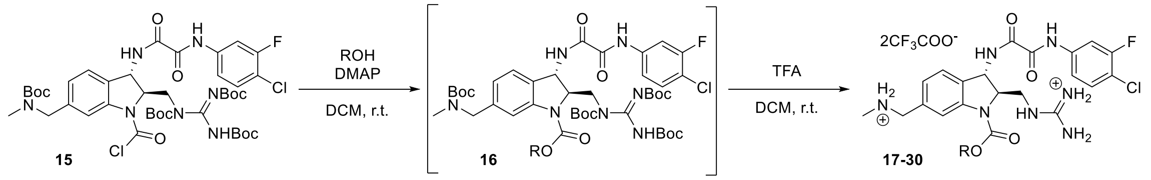

Compound **15** was dissolved in $CH_2Cl_2$ (0.1 M), followed by addition of the alcohol desired for carbamate formation (2.0 equiv.). N,N-dimethylaminopyridine (1.5 equiv.) was then added in one portion, and the mixture was allowed to stir overnight, at which time UPLCMS analysis indicated consumption of **15**. The reaction was quenched with sat. aq. $NH_4Cl$, and the aqueous layer was extracted 3 x $CH_2Cl_2$. The organic layers were separated, dried with $Na_2SO_4$, filtered and concentrated *in vacuo*. The crude **16** (1.0 equiv.) was then taken up in $CH_2Cl_2$ (0.1 M) and cooled to 0 °C in an ice-water bath. Trifluoroacetic acid (40 equiv.) was added and the mixture was allowed to warm to rt. The reaction was allowed to stir for 18 hours. The solution was then concentrated *in vacuo* and the resulting crude residue was purified by flash column chromatography (10% MeOH/$CH_2Cl_2$) to give the products **17-30** as an amorphous white solids.

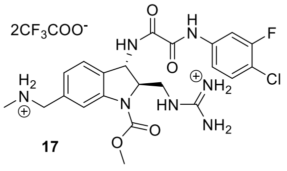

**17**

**Compound 17 $^1$H NMR** (500 MHz, $CD_3OD$) δ 7.89 (brs, 1H), 7.85 – 7.80 (dd, d, *J* = 11.4, 1.9 Hz, 1H), 7.47 – 7.42 (m, 3H), 7.22 (d, *J* = 7.8 Hz, 1H), 5.23 (s, 1H), 4.58 – 4.53 (m, 1H), 4.22 (s, 2H), 3.91 (s, 3H), 3.56 (d, *J* = 5.2 Hz, 2H), 2.73 (s, 3H); **$^{13}$C NMR** (125 MHz, $CD_3OD$) δ 161.44, 159.50, 159.43, 159.15 (d, $J_{CF}$ = 245.5 Hz), 139.11 (d, $J_{CF}$ = 9.8 Hz), 134.74, 131.88, 131.53, 128.04, 126.57, 118.38, 118.22 (d, $J_{CF}$ = 3.6 Hz), 117.46 (d, $J_{CF}$ = 18.3 Hz), 109.92 (d, $J_{CF}$ = 26.4 Hz) 66.87, 54.57, 54.11, 53.70, 49.77, 44.49, 33.25; **HRMS** (ESI) *m/z* 506.1706 [calcd for $C_{22}H_{26}ClFN_7O_4$ $(M+H)^+$ 506.1719]; **$[\alpha]_D^{24}$** +40.8 (*c* 0.57, MeOH).

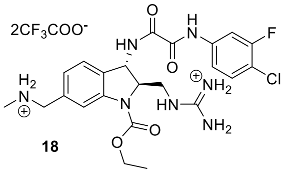

**18**

**Compound 18 $^1$H NMR** (500 MHz, $CD_3OD$) δ 7.90 (brs, 1H), 7.83 (dd, *J* = 11.3, 2.1 Hz, 1H), 7.49 – 7.42 (m, 3H), 7.22 (d, *J* = 7.8 Hz, 1H), 5.23 (d, *J* = 1.8 Hz, 1H), 4.57 – 4.54 (m, 1H), 4.36 (brs, 1H), 4.22 (s, 2H), 3.58 (dd, *J* = 14.4, 5.5 Hz, 1H), 3.54 (dd, *J* = 14.1, 7.0 Hz, 1H), 2.73 (s, 3H), 1.40 (t, *J* = 7.0 Hz, 3H); **$^{13}$C NMR** (150 MHz, $CD_3OD$) δ 163.06 (q, $J_{CF}$ = 35.5 Hz, TFA),161.44, 159.51, 159.40, 159.28 (d, $J_{CF}$ = 246.0 Hz), 139.12 (d, $J_{CF}$ = 10.0 Hz), 134.69, 131.87, 131.54, 128.05, 126.52, 118.49, 118.23 (d, $J_{CF}$ = 3.4 Hz), 117.44 (d, $J_{CF}$ = 17.9 Hz), 109.91 (d, $J_{CF}$ = 26.2 Hz), 66.83, 64.03, 54.55, 53.72, 49.72, 44.53, 33.24, 14.95; **HRMS** (ESI) *m/z* 520.1890 [calcd for $C_{23}H_{28}ClFN_7O_4$ $(M+H)^+$ 520.1875]; **$[\alpha]_D^{24}$** +14.6 (*c* 0.22, MeOH).

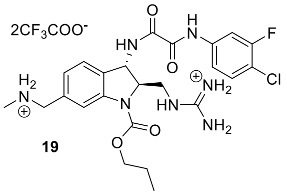

**19**

**Compound 19 $^1$H NMR** (500 MHz, $CD_3OD$) δ 7.89 (brs, 1H), 7.83 (dd, *J* = 11.3, 2.2 Hz, 1H), 7.50 – 7.42 (m, 3H), 7.22 (d, *J* = 7.7 Hz, 1H), 5.24 (d, *J* = 2.0 Hz, 1H), 4.57 – 4.54 (m, 1H), 4.32 – 4.19 (m, 4H), 3.60 (dd, *J* = 14.1, 5.1 Hz, 1H), 3.54 (dd, *J* = 14.0, 7.0 Hz, 1H), 2.73 (s, 3H), 1.81 (sxt, *J* = 7.0 Hz, 2H), 1.03 (t, *J* = 7.4 Hz, 3H); **$^{13}$C NMR** (150 MHz, $CD_3OD$) δ 163.13 (q, $J_{CF}$ = 36.7 Hz, TFA), 161.47, 159.50, 159.40, 159.29 (d, $J_{CF}$ = 244.2 Hz), 139.12 (d, $J_{CF}$ = 9.9 Hz), 134.68, 131.88, 128.07, 126.50, 118.47, 118.22 (d, $J_{CF}$ = 3.5 Hz), 117.45 (d, $J_{CF}$ = 18.0 Hz), 109.91 (d, $J_{CF}$ = 26.4 Hz), 66.90, 54.56, 53.73, 49.72, 44.56, 33.24, 23.39, 10.85; **HRMS** (ESI) *m/z* 534.2027 [calcd for $C_{24}H_{30}ClFN_7O_4$ $(M+H)^+$ 534.2032]; **$[\alpha]_D^{24}$** +20.6 (*c* 1.56, MeOH).

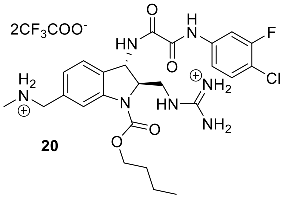

**20**

**Compound 20 $^1$H NMR** (600 MHz, $CD_3OD$) δ 7.89 (s, 1H), 7.83 (dd, *J* = 11.3, 2.4 Hz, 1H), 7.49 – 7.46 (m, 2H), 7.43 (t, *J* = 8.4 Hz, 1H), 7.22 (dd, *J* = 7.8, 1.6 Hz, 1H), 5.23 (d, *J* = 2.4 Hz, 1H), 4.55 (ddd, *J* = 7.4, 5.1, 2.4 Hz, 1H), 4.32 (t, *J* = 8.1 Hz, 2H), 4.21 (s, 2H), 3.59 (dd, *J* = 14.2, 5.1 Hz, 1H), 3.55 (dd, *J* = 14.2, 6.8 Hz, 1H), 2.72 (s, 3H), 1.77 (t, *J* = 7.5 Hz, 2H), 1.47 (h, *J* = 7.5 Hz, 2H), 0.99 (t, *J* = 7.4 Hz, 3H); **$^{13}$C NMR** (150 MHz, $CD_3OD$) δ 162.23 (q, $J_{CF}$ = 35.4 Hz, TFA), 161.43, 159.51, 159.43, 159.24 (d, $J_{CF}$ = 246.7 Hz), 139.12 (d, $J_{CF}$ = 9.8 Hz), 134.65, 131.83, 131.42, 127.99, 126.51, 118.51, 118.23 (d, $J_{CF}$ = 3.5 Hz), 117.38 (d, $J_{CF}$ = 17.9 Hz), 109.90 (d, $J_{CF}$ = 26.1 Hz), 66.93, 54.58, 53.73, 50.00, 44.52, 33.22, 32.11, 20.32, 14.20; **HRMS** (ESI) *m/z* 548.2173 [calcd for $C_{25}H_{31}ClFN_7O_4$ $(M+H)^+$ 548.2188]. **$[\alpha]_D^{24}$** +33.4 (*c* 0.38, MeOH).

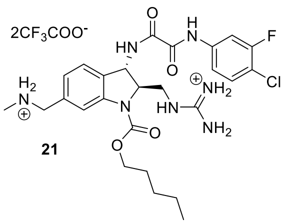

21

**Compound 21** $^{1}$**H NMR** (600 MHz, $CD_3OD$) δ 7.91 (s, 1H), 7.83 (dd, $J$ = 11.3, 2.3 Hz, 1H), 7.51 – 7.42 (m, 3H), 7.22 (dd, $J$ = 7.8, 1.6 Hz, 1H), 5.24 (d, $J$ = 2.5 Hz, 1H), 4.55 (ddd, $J$ = 7.4, 5.0, 2.4 Hz, 1H), 4.36 – 4.27 (m, 1H), 4.22 (s, 2H), 3.60 (dd, $J$ = 14.1, 5.1 Hz, 1H), 3.52 (dd, $J$ = 14.0, 7.2 Hz, 1H), 2.73 (s, 3H), 1.80 (s, 2H), 1.42 (dd, $J$ = 7.0, 3.5 Hz, 4H), 0.97 – 0.93 (m, 3H); $^{13}$**C NMR** (150 MHz, $CD_3OD$) δ 161.49, 159.47, 159.38, 159.29 (d, $J_{CF}$ = 245.7 Hz), 139.10 (d, $J_{CF}$ = 10.0 Hz), 134.72, 131.89, 126.48, 118.21 (d, $J_{CF}$ = 3.4 Hz), 117.48 (d, $J_{CF}$ = 17.9 Hz), 109.91 (d, $J_{CF}$ = 26.2 Hz), 66.90, 54.60, 53.75, 49.72, 44.57, 33.26, 29.79, 29.35, 23.56, 14.49; **HRMS** (ESI) *m/z* 562.2356 [calcd for $C_{26}H_{34}ClFN_7O_4$ $(M+H)^+$ 562.2345]; $[\alpha]_D^{24}$ +40.5 ($c$ 0.18, MeOH).

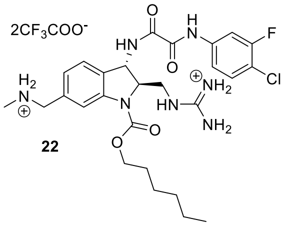

22

**Compound 22** $^{1}$**H NMR** (500 MHz, $CD_3OD$) δ 7.91 (brs, 1H), 7.83 (dd, $J$ = 11.4, 2.3 Hz, 1H), 7.49 – 7.41 (m, 3H), 7.22 (d, $J$ = 7.7 Hz, 1H), 5.23 (d, $J$ = 1.6 Hz, 1H), 4.57 – 4.52 (m, 1H), 4.34 – 4.25 (m, 2H), 4.21 (s, 2H), 3.59 (dd, $J$ = 14.1, 5.1 Hz, 1H), 3.54 (dd, $J$ = 14.1, 6.9 Hz, 1H), 2.72 (s, 3H), 1.81 – 1.75 (m, 2H), 1.46 – 1.40 (m, 2H), 1.39 – 1.27 (m, 4H), 0.93 – 0.89 (m, 3H); $^{13}$**C NMR** (150 MHz, $CD_3OD$) δ 163.31 (q, $J_{CF}$ = 35.5 Hz, TFA), 161.47, 159.50, 159.44, 159.37 (d, $J_{CF}$ = 246.2 Hz), 139.13 (d, $J_{CF}$ = 9.6 Hz), 134.67, 131.85, 128.02, 126.51, 118.50, 118.23 (d, $J_{CF}$ = 3.5 Hz), 117.40 (d, $J_{CF}$ = 17.6 Hz), 109.75 (d, $J_{CF}$ = 26.2 Hz), 70.74, 66.95, 56.18, 54.61, 53.73, 44.53, 33.21, 32.75, 32.24, 30.01, 29.68, 26.83, 23.77, 14.48; **HRMS** (ESI) *m/z* 576.2511 [calcd for $C_{27}H_{36}ClFN_7O_4$ $(M+H)^+$ 576.2501]; $[\alpha]_D^{24}$ +11.0 ($c$ 1.69, MeOH).

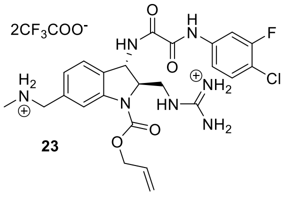

23

**Compound 23** $^{1}$**H NMR** (500 MHz, $CD_3OD$) δ 7.91 (brs, 1H), 7.83 (dd, $J$ = 11.3, 2.1 Hz, 1H), 7.50 – 7.42 (m, 3H), 7.23 (d, $J$ = 7.8, Hz, 1H), 6.12 – 6.04 (m, 1H), 5.42 (d, $J$ = 17.2 Hz, 1H), 5.31 (d, $J$ = 10.4 Hz, 1H), 5.24 (d, $J$ = 1.8 Hz, 1H), 6.12 – 6.04 (m, 1H), 4.22 (s, 2H), 3.59 (dd, $J$ = 14.2, 5.3 Hz, 1H), 3.54 (dd, $J$ = 14.1, 6.9 Hz, 1H), 2.73 (s, 3H); $^{13}$**C NMR** (150 MHz, $CD_3OD$) δ 163.01 (q, $J_{CF}$ = 35.5 Hz, TFA), 161.46, 159.49, 159.38, 159.28 (d, $J_{CF}$ = 246.1 Hz), 139.04 (d, $J_{CF}$ = 9.9 Hz), 134.72, 133.68, 131.88, 128.08, 126.62, 119.54, 118.48, 118.22 (d, $J_{CF}$ = 3.4 Hz), 117.46 (d, $J_{CF}$ = 17.8 Hz), 109.92 (d, $J_{CF}$ = 26.1 Hz), 66.89, 54.56, 53.71, 49.72, 44.51, 33.25; **HRMS** (ESI) *m/z* 532.1870 [calcd for $C_{24}H_{28}ClFN_7O_4$ $(M+H)^+$ 532.1875]; $[\alpha]_D^{24}$ +35.9 ($c$ 0.25, MeOH).

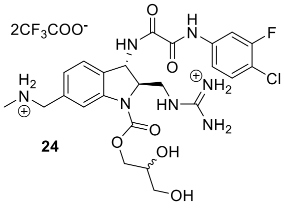

24

**Compound 24** $^{1}$**H NMR** (500 MHz, $CD_3OD$) δ 7.96 (brs, 1H), 7.83 (dd, $J$ = 11.3, 2.1 Hz, 1H), 7.52 – 7.42 (m, 3H), 7.22 (d, $J$ = 7.7 Hz, 1H), 5.27 (d, $J$ = 1.5 Hz, 1H), 4.63 – 4.57 (m, 1H), 4.48 (dd, $J$ = 11.3, 3.4 Hz, 0.5H), 4.43 – 4.25 (m, 1.5H), 4.22 (s, 2H), 4.01 – 3.96 (m, 1H), 3.68 – 3.59 (m, 3H), 3.58 – 3.53 (m, 1H), 2.73 (s, 3H); $^{13}$**C NMR** (150 MHz, $CD_3OD$) δ 161.40, 159.50, 159.30, 159.24 (d, $J_{CF}$ = 245.7 Hz), 139.11 (d, $J_{CF}$ = 9.9 Hz), 134.72, 131.89, 128.06, 126.62, 118.22 (d, $J_{CF}$ = 3.7 Hz), 117.46 (d, $J_{CF}$ = 18.2 Hz), 109.91 (d, $J_{CF}$ = 26.2 Hz), 71.29, 66.95, 64.18, 54.65, 53.69, 49.72, 44.48, 33.24; **HRMS** (ESI) *m/z* 566.1927 [calcd for $C_{24}H_{30}ClFN_7O_6$ $(M+H)^+$ 566.1930]. $[\alpha]_D^{24}$ +15.6 ($c$ 0.26, MeOH).

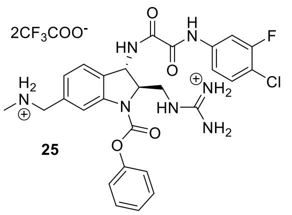

**Compound 25** $^{1}$**H NMR** (600 MHz, $CD_3OD$) δ 7.98 (brs, 1H), 7.84 (dd, *J* = 11.3, 2.3 Hz, 1H), 7.56 (d, *J* = 7.8 Hz, 1H), 7.49 (dd, *J* = 9.0, 2.1 Hz, 1H), 7.47 – 7.43 (m, 3H), 7.32 – 7.27 (m, 4H), 5.29 (d, *J* = 1.6 Hz, 1H), 4.78 – 4.58 (br m, 1H, obscured by residual MeOH), 4.23 (s, 2H), 3.67 (brs, 2H), 2.72 (brs, 3H); $^{13}$**C NMR** (150 MHz, $CD_3OD$) δ 161.44, 159.54, 159.42, 159.15 (d, $J_{CF}$ = 245.5 Hz), 152.16, 139.12 (d, $J_{CF}$ = 9.8 Hz), 131.90, 130.75, 127.17, 123.08, 118.23 (d, $J_{CF}$ = 3.7 Hz), 117.47 (d, $J_{CF}$ = 18.1 Hz), 109.93 (d, $J_{CF}$ = 26.1 Hz), 53.65, 49.72, 33.29; **HRMS** (ESI) *m/z* 568.1885 [calcd for $C_{27}H_{28}ClFN_7O_4$ $(M+H)^+$ 568.1875]; $[\alpha]_D^{24}$ +38.8 (*c* 0.27, MeOH).

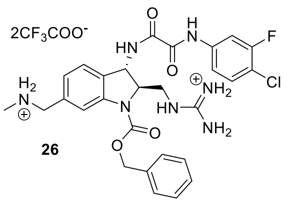

**Compound 26** $^{1}$**H NMR** (500 MHz, $CD_3OD$) δ 7.93 (brs, 1H), 7.82 (dd, *J* = 11.3, 2.0 Hz, 1H), 7.57 – 7.54 (m, 6H), 7.49 – 7.40 (m, 6H), 7.22 (dd, *J* = 7.8, 1.4 Hz, 1H), 5.37 (brs, 2H), 5.25 (d, *J* = 2.2 Hz, 1H), 4.57 (ddd, *J* = 7.4, 4.8, 2.5 Hz, 1H), 4.21 (s, 2H), 3.59 (dd, *J* = 14.0, 4.9 Hz, 1H), 3.51 (dd, *J* = 14.1, 7.3 Hz, 1H), 2.70 (s, 3H); $^{13}$**C NMR** (150 MHz, $CD_3OD$) δ 163.15 (q, $J_{CF}$ = 35.5 Hz, TFA), 161.46, 159.48, 159.36, 159.28 (d, $J_{CF}$ = 245.3 Hz), 139.10 (d, $J_{CF}$ = 10.5 Hz), 137.39, 134.67, 131.87, 129.90, 129.80, 129.74, 128.05, 126.62, 118.51, 118.22 (d, $J_{CF}$ = 3.6 Hz), 117.48 (d, $J_{CF}$ = 18.0 Hz), 109.92 (d, $J_{CF}$ = 26.4 Hz), 69.45, 66.97, 54.57, 53.69, 49.72, 44.49, 33.19; **HRMS** (ESI) *m/z* 582.2029 [calcd for $C_{28}H_{30}ClFN_7O_4$ $(M+H)^+$ 582.2032]; $[\alpha]_D^{24}$ +42.5 (*c* 0.15, MeOH).

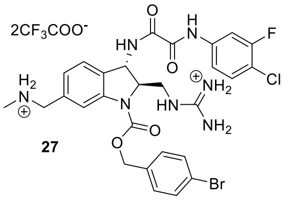

**Compound 27** $^{1}$**H NMR** (500 MHz, $CD_3OD$) δ 7.91 (s, 1H), 7.82 (dd, *J* = 11.3, 2.3 Hz, 1H), 7.56 (d, *J* = 8.4 Hz, 2H), 7.49 (d, *J* = 7.6 Hz, 1H), 7.47 – 7.42 (m, 2H), 7.41 (d, *J* = 8.2 Hz, 2H), 7.22 (dd, *J* = 7.7, 1.6 Hz, 1H), 5.35 (d, *J* = 11.8 Hz, 1H), 5.25 (d, *J* = 2.5 Hz, 1H), 4.57 (d, *J* = 4.5 Hz, 1H), 4.19 (s, 2H), 3.59 (dd, *J* = 14.0, 5.0 Hz, 1H), 3.52 (dd, *J* = 14.1, 7.2 Hz, 1H), 2.70 (s, 3H); $^{13}$**C NMR** (150 MHz, $CD_3OD$) δ 161.47, 159.47, 159.36, 159.28 (d, $J_{CF}$ = 245.3 Hz), 139.10 (d, $J_{CF}$ = 10.5 Hz), 136.74, 133.03, 132.57, 131.89, 131.57, 129.95, 126.66, 123.63, 118.22 (d, $J_{CF}$ = 3.6 Hz), 117.48 (d, $J_{CF}$ = 18.0 Hz), 109.92 (d, $J_{CF}$ = 26.4 Hz), 66.96, 53.75, 49.72, 44.51, 33.28; **HRMS** (ESI) *m/z* 660.1144 [calcd for $C_{28}H_{29}BrClFN_7O_4$ $(M+H)^+$ 660.1137]; $[\alpha]_D^{24}$ +31.2 (*c* 0.19, MeOH).

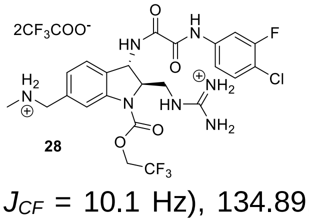

**Compound 28** $^{1}$**H NMR** (500 MHz, $CD_3OD$) δ 8.01 (brs, 1H), 7.83 (dd, *J* = 11.3, 2.3 Hz, 1H), 7.53 (d, *J* = 7.8 Hz, 1H), 7.50 – 7.41 (m, 2H), 7.27 (d, *J* = 7.5 Hz, 1H), 5.28 (d, *J* = 2.0 Hz, 1H), 4.86 (s, 2H, obscured by residual $CH_3OH$), 4.62 – 4.59 (m, 1H), 4.23 (brs, 2H), 3.61 (dd, *J* = 14.1, 5.4 Hz, 1H), 3.56 (dd, *J* = 14.2, 6.7 Hz, 1H), 2.73 (s, 3H); $^{13}$**C NMR** (150 MHz, $CD_3OD$) δ 161.50, 159.45, 159.42, 159.31 (d, $J_{CF}$ = 245.8 Hz), 139.11 (d, $J_{CF}$ = 10.1 Hz), 134.89, 131.90, 127.20, 124.87 (q, $J_{CF}$ = 276.8 Hz), 118.06 (d, $J_{CF}$ = 3.4 Hz), 117.26 (d, $J_{CF}$ = 17.4 Hz), 109.75 (d, $J_{CF}$ = 26.3 Hz), 53.67, 49.72, 44.35, 33.26; **HRMS** (ESI) *m/z* 574.1573 [calcd for $C_{23}H_{25}ClF_4N_7O_4$ $(M+H)^+$ 574.1593]. $[\alpha]_D^{24}$ +31.2 (*c* 0.19, MeOH).

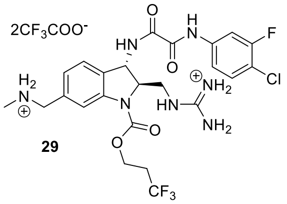

29

**Compound 29** $^1$**H NMR** (500 MHz, $CD_3OD$) δ 7.96 (brs, 1H), 7.82 (dd, *J* = 11.3, 2.2 Hz, 1H), 7.52 – 7.43 (m, 3H), 7.24 (dd, J = 7.9, 1.3 Hz, 1H), 5.25 (d, *J* = 1.8 Hz, 1H), 4.56 - 4.54 (m, 3H), 4.22 (s, 2H), 3.62 (dd, J = 14.1, 5.1 Hz, 1H), 3.52 (dd, J = 14.0, 7.2 Hz, 1H), 3.32 (m, 2H, obscured by residual MeOH) 2.72 (s, 3H); $^{13}$**C NMR** (150 MHz, $CD_3OD$) δ 161.65, 159.58, 159.54, 159.30 (d, $J_{CF}$ = 245.6 Hz), 139.09 (d, $J_{CF}$ = 10.2 Hz), 134.88, 132.04, 131.54, 127.13, 126.88, 118.42, 118.22 (d, $J_{CF}$ = 3.0 Hz), 117.51 (d, $J_{CF}$ = 17.7 Hz), 109.92 (d, $J_{CF}$ = 26.1 Hz), 67.14, 63.2, 54.74, 53.82, 49.87, 44.50, 34.27 (q, $J_{CF}$ = 28.7 Hz), 33.36, 24.51; **HRMS** (ESI) *m/z* 588.1746 [calcd for $C_{24}H_{26}ClF_4N_7O_4$ $(M+H)^+$ 588.1749]. $[\alpha]_D^{23}$ +14.85 (*c* 0.21, $CH_3OH$).

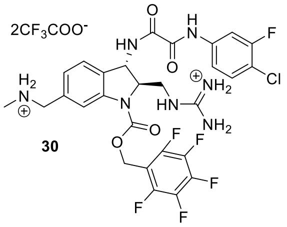

30

**Compound 30** $^1$**H NMR** (500 MHz, $CD_3OD$) δ 7.98 (brs, 1H), 7.82 (dd, *J* = 11.3, 2.2 Hz, 1H), 7.49 – 7.42 (m, 3H), 7.23 (dd, *J* = 7.8, 1.0 Hz, 1H), 5.51 (brs, 2H), 5.23 (d, *J* = 2.4 Hz, 1H), 4.53 (ddd, *J* = 7.1, 5.0, 2.2 Hz, 1H), 4.21 (s, 2H), 3.57 (dd, *J* = 14.1, 5.1 Hz, 1H), 3.51 (dd, *J* = 14.1, 7.0 Hz, 1H), 2.72 (s, 3H); $^{13}$**C NMR** (150 MHz, $CD_3OD$) δ 161.46 (q, $J_{CF}$ = 35.5 Hz, TFA), 159.44, 159.42, 159.30 (d, $J_{CF}$ = 245.8 Hz), 139.13, 139.06, 134.72, 131.88, 128.06, 126.88, 118.51, 118.21 (d, *J* = 3.5 Hz), 117.47 (d, *J* = 18.0 Hz), 116.63, 109.90 (d, *J* = 26.2 Hz), 66.98, 56.31, 54.58, 53.65, 52.02, 49.72, 44.43, 33.20, 29.00, 23.76; **HRMS** (ESI) *m/z* 672.1536 [calcd for $C_{28}H_{25}ClF_6N_7O_4$ $(M+H)^+$ 672.1561]; $[\alpha]_D^{24}$ +34.5 (*c* 0.47, MeOH).

**$^{1}$H and $^{13}$C NMR Spectra**

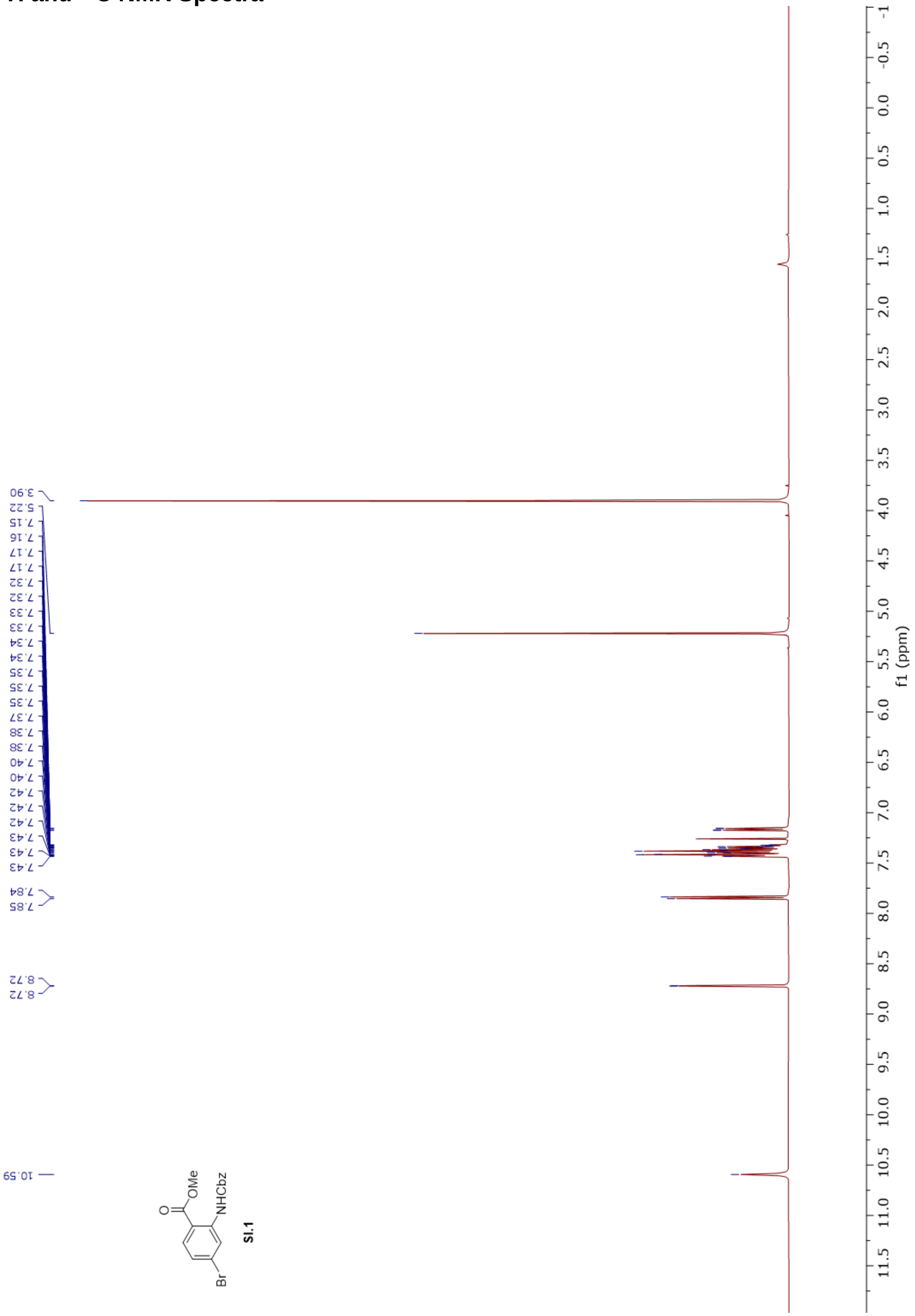

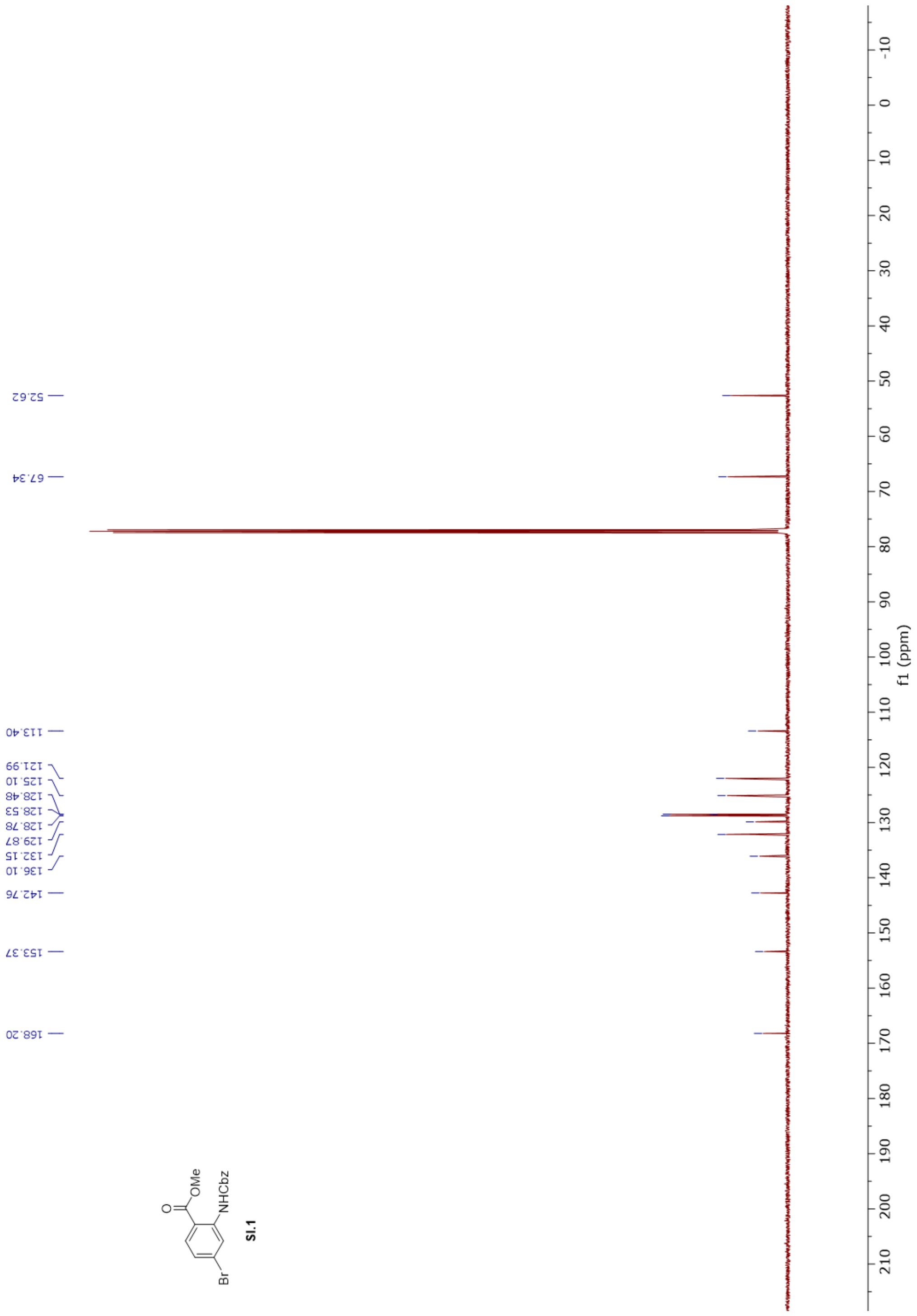
52.62
67.34
113.40
121.99
125.10
128.48
128.53
128.78
129.87
132.15
136.10
142.76
153.37
168.20
O
OMe
NHCbz
Br
SI.1
f1 (ppm)

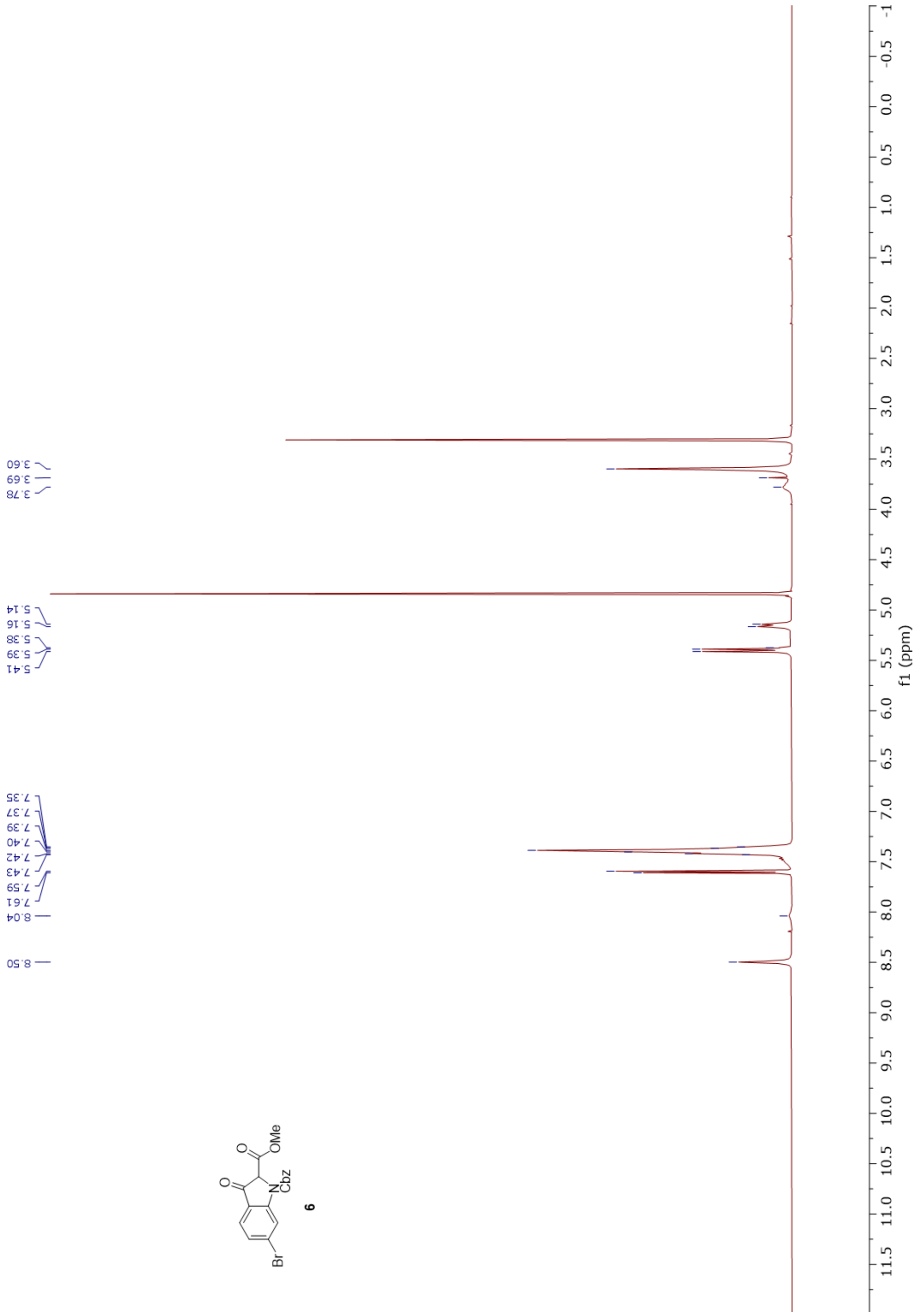
3.60
3.69
3.78
5.14
5.16
5.38
5.39
5.41
7.35
7.37
7.39
7.40
7.42
7.43
7.59
7.61
8.04
8.50
OMe
Cbz
Br
6
f1 (ppm)

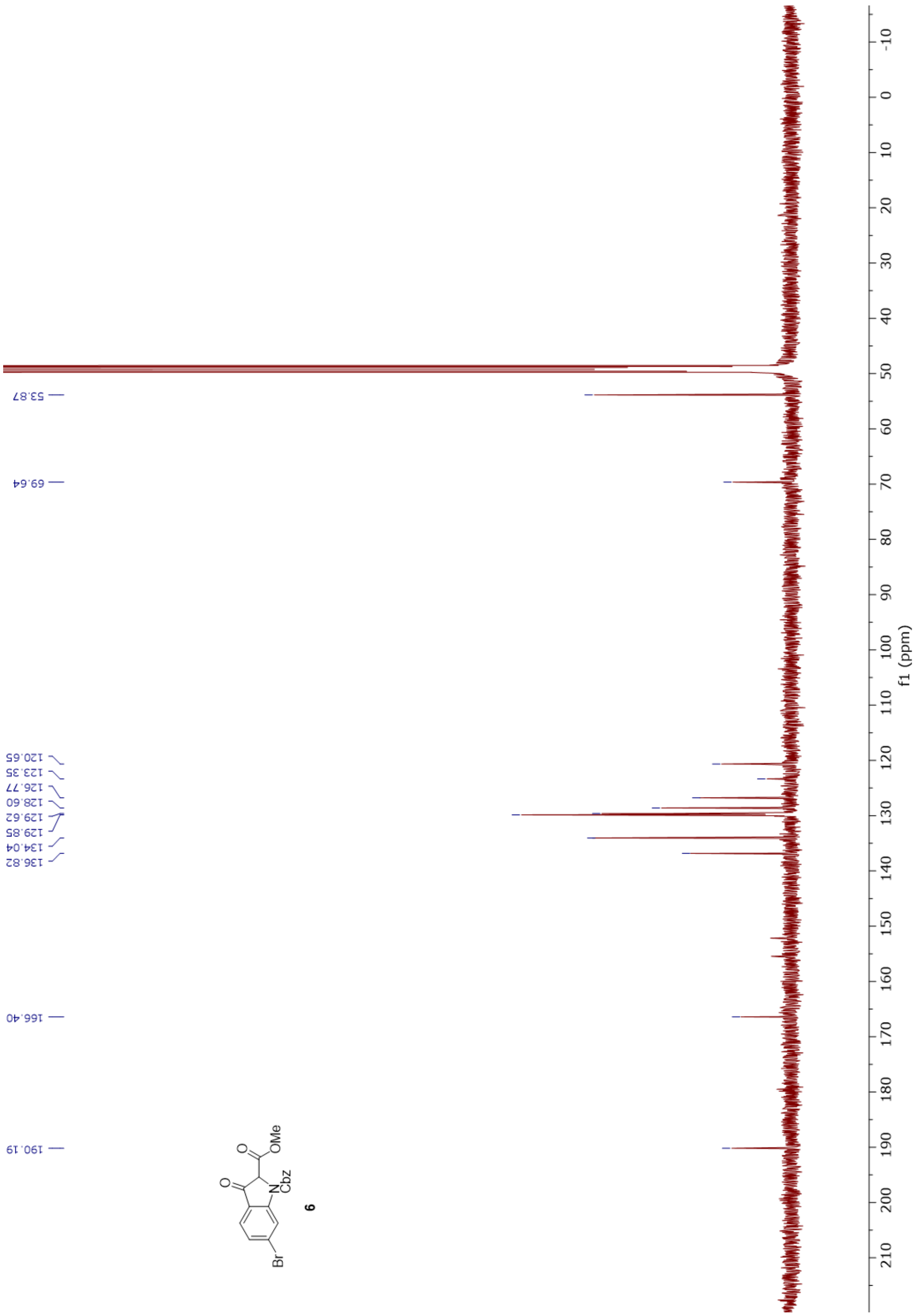
53.87
69.64
120.65
123.35
126.77
128.60
129.62
129.85
134.04
136.82
166.40
190.19
f1 (ppm)
OMe
Cbz
Br
6

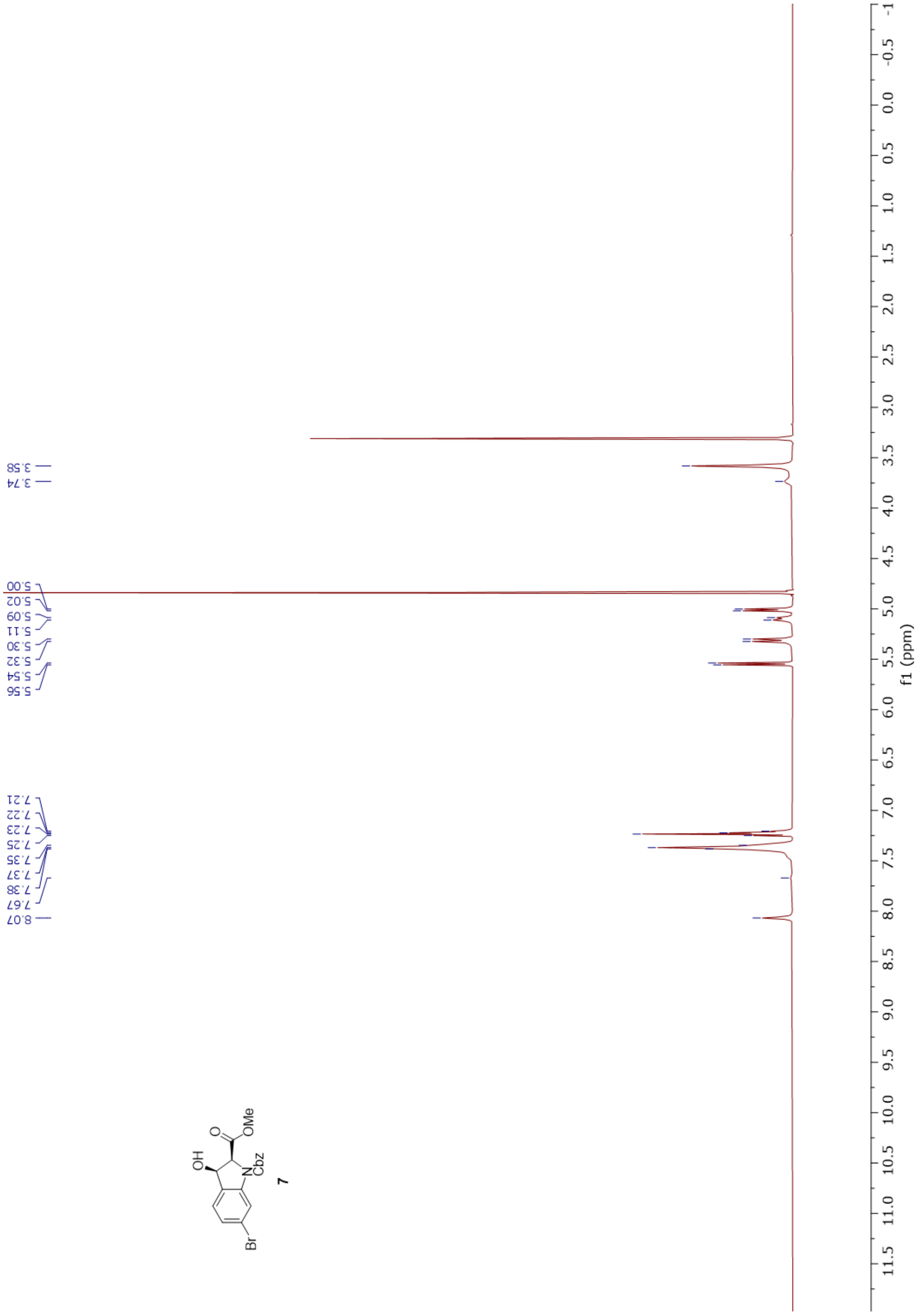
3.58
3.74
5.00
5.02
5.09
5.11
5.30
5.32
5.54
5.56
7.21
7.22
7.23
7.25
7.35
7.37
7.38
7.67
8.07
OH
O
OMe
N
Cbz
Br
7
f1 (ppm)

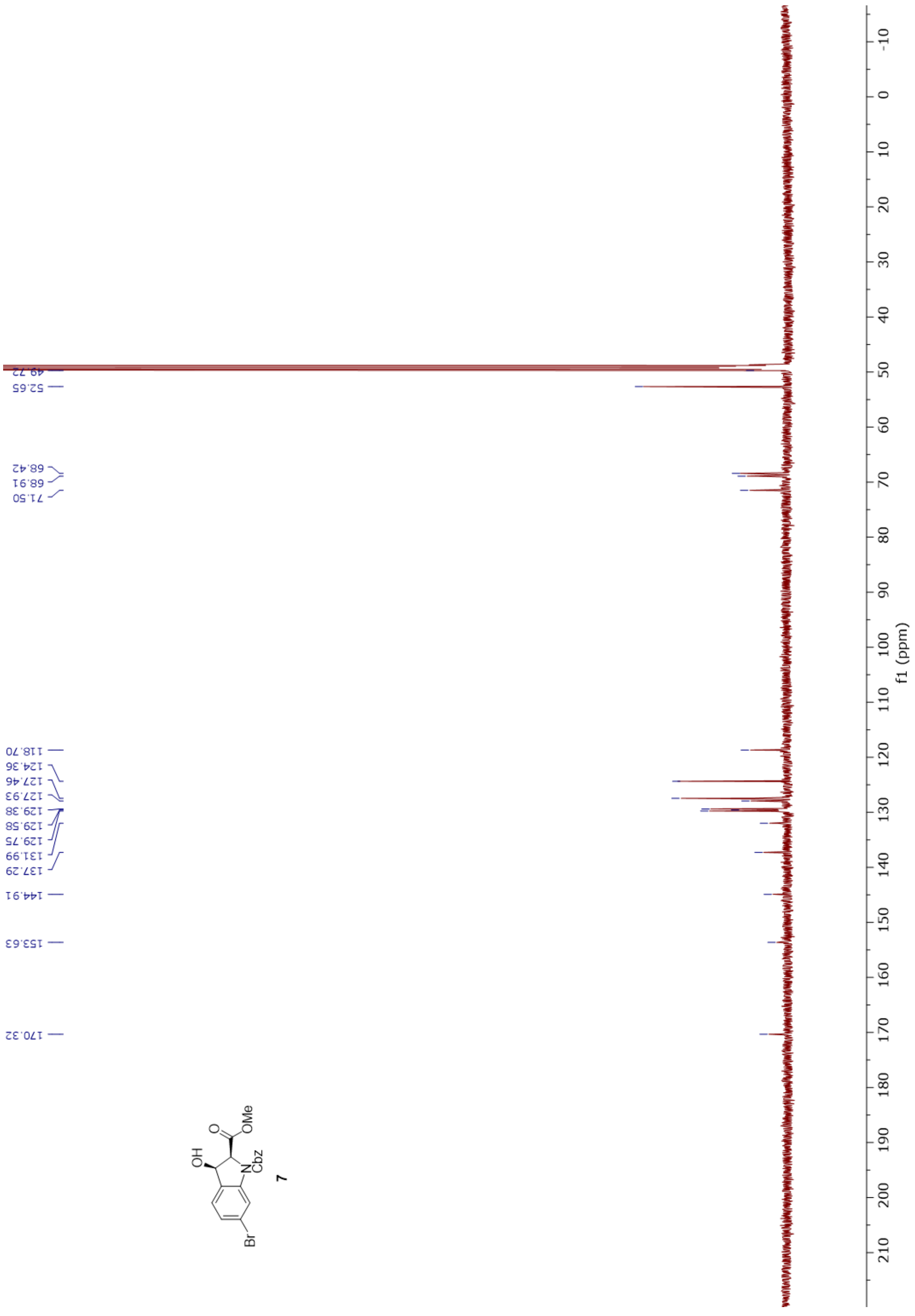

49.72
52.65
68.42
68.91
71.50
118.70
124.36
127.46
127.93
129.38
129.58
129.75
131.99
137.29
144.91
153.63
170.32
f1 (ppm)
OH
O
OMe
N
Cbz
Br
7

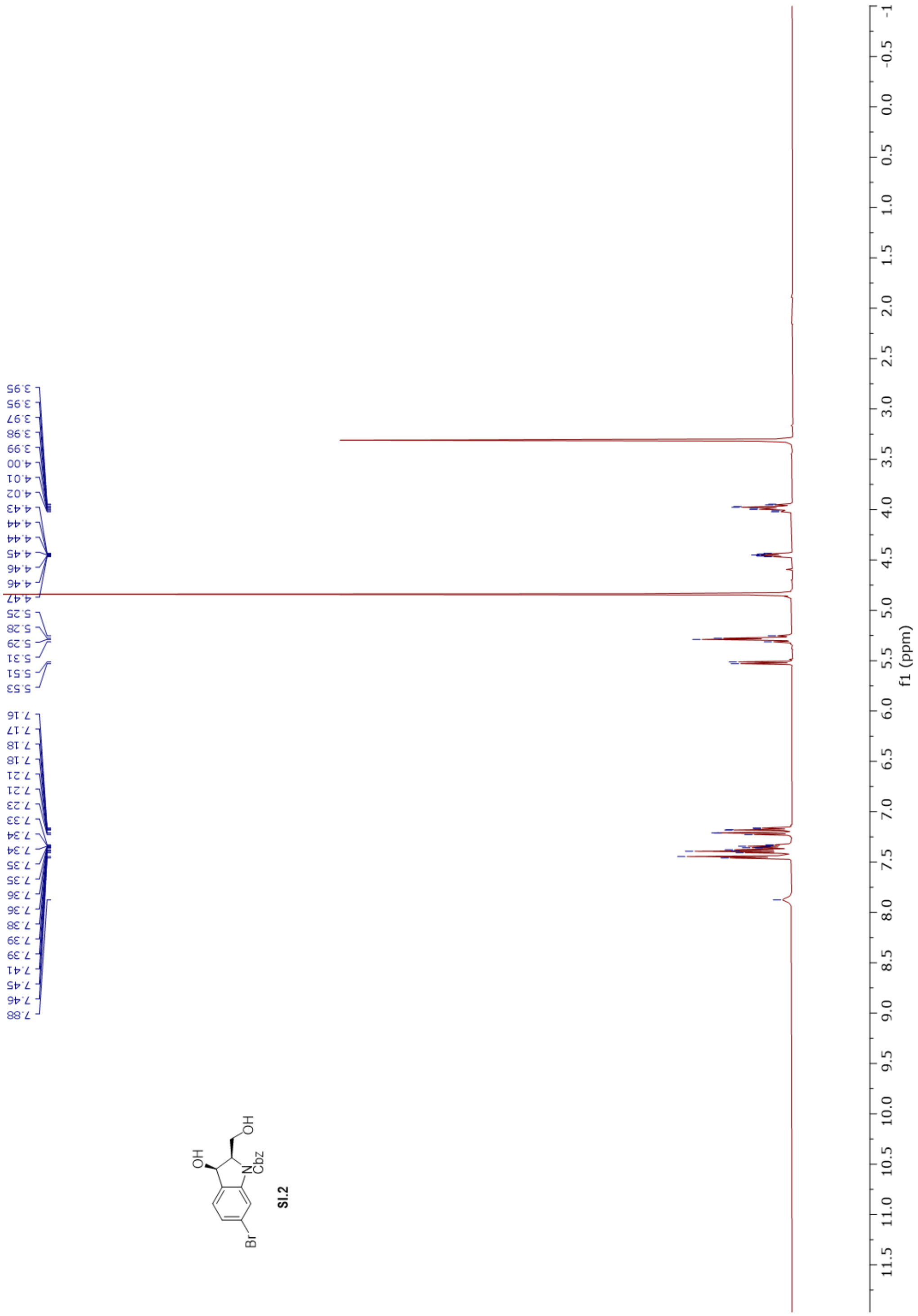
3.95
3.95
3.97
3.98
3.99
4.00
4.01
4.02
4.43
4.44
4.44
4.45
4.46
4.46
4.47
5.25
5.28
5.29
5.31
5.51
5.53
7.16
7.17
7.18
7.18
7.21
7.21
7.23
7.33
7.34
7.34
7.35
7.35
7.36
7.36
7.38
7.39
7.39
7.41
7.45
7.46
7.88
Br
OH
N
Cbz
OH
SI.2
f1 (ppm)

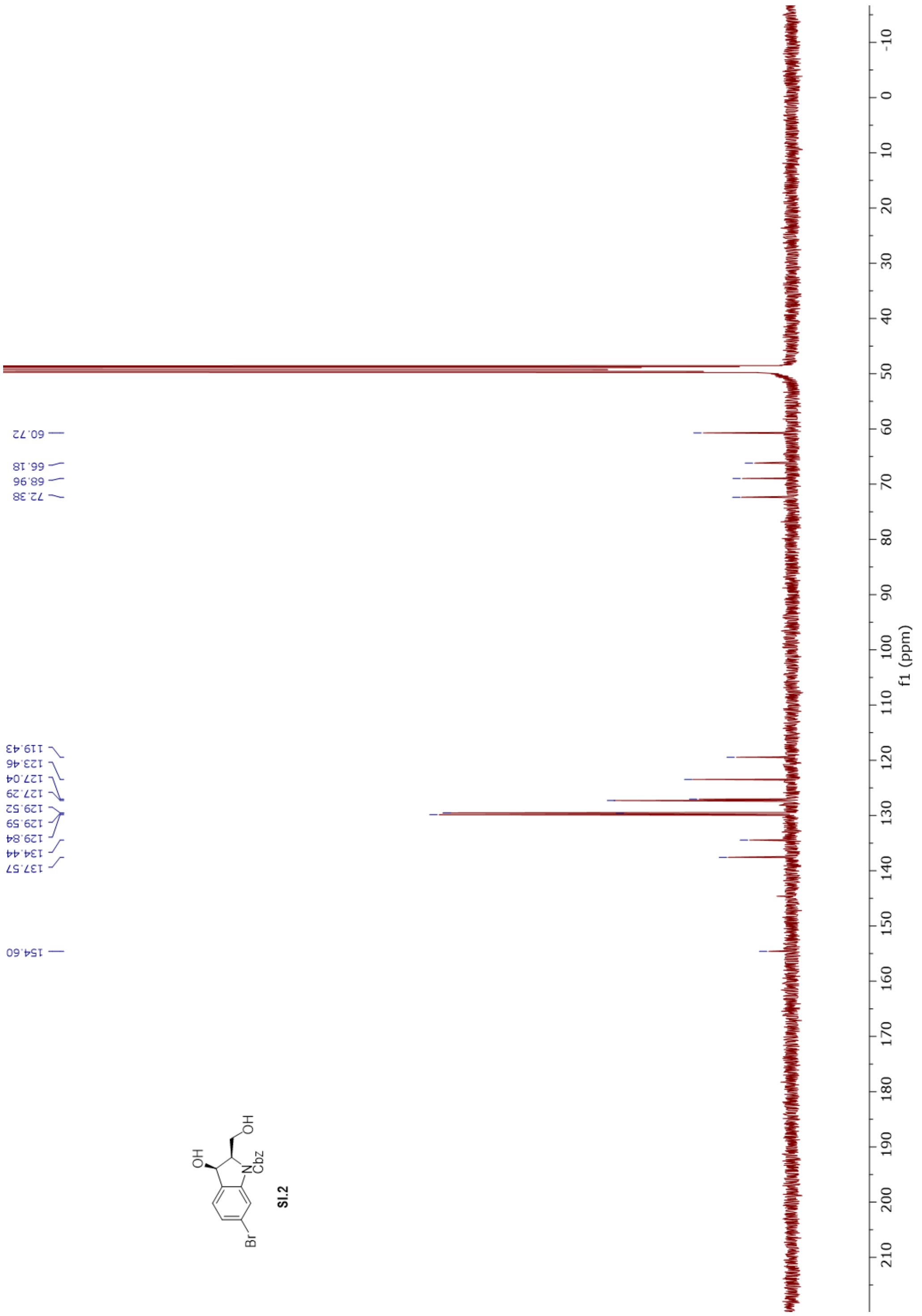
60.72
66.18
68.96
72.38
119.43
123.46
127.04
127.29
129.52
129.59
129.84
134.44
137.57
154.60
OH
OH
N
Cbz
Br
SI.2
f1 (ppm)

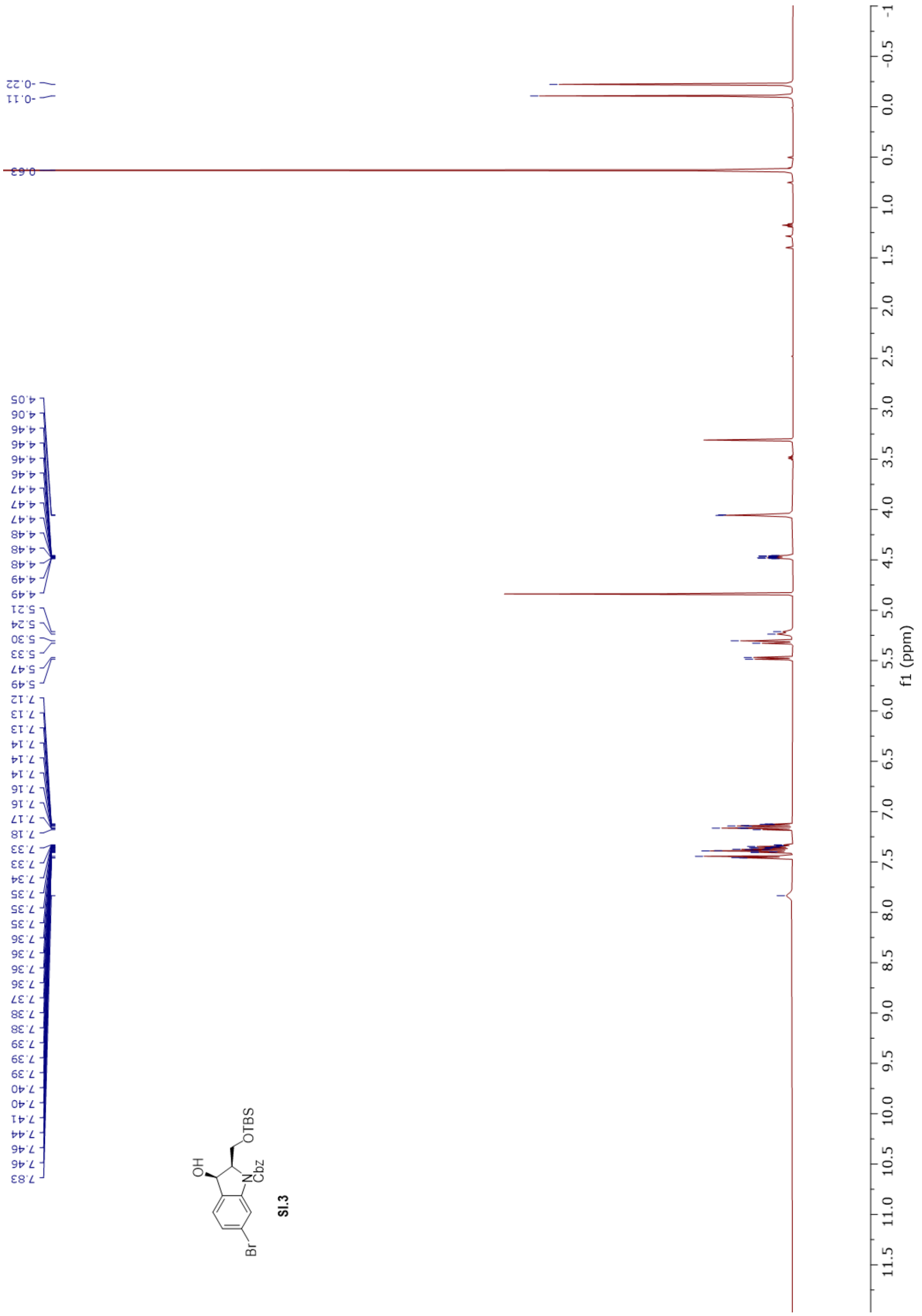

SI.3
f1 (ppm)

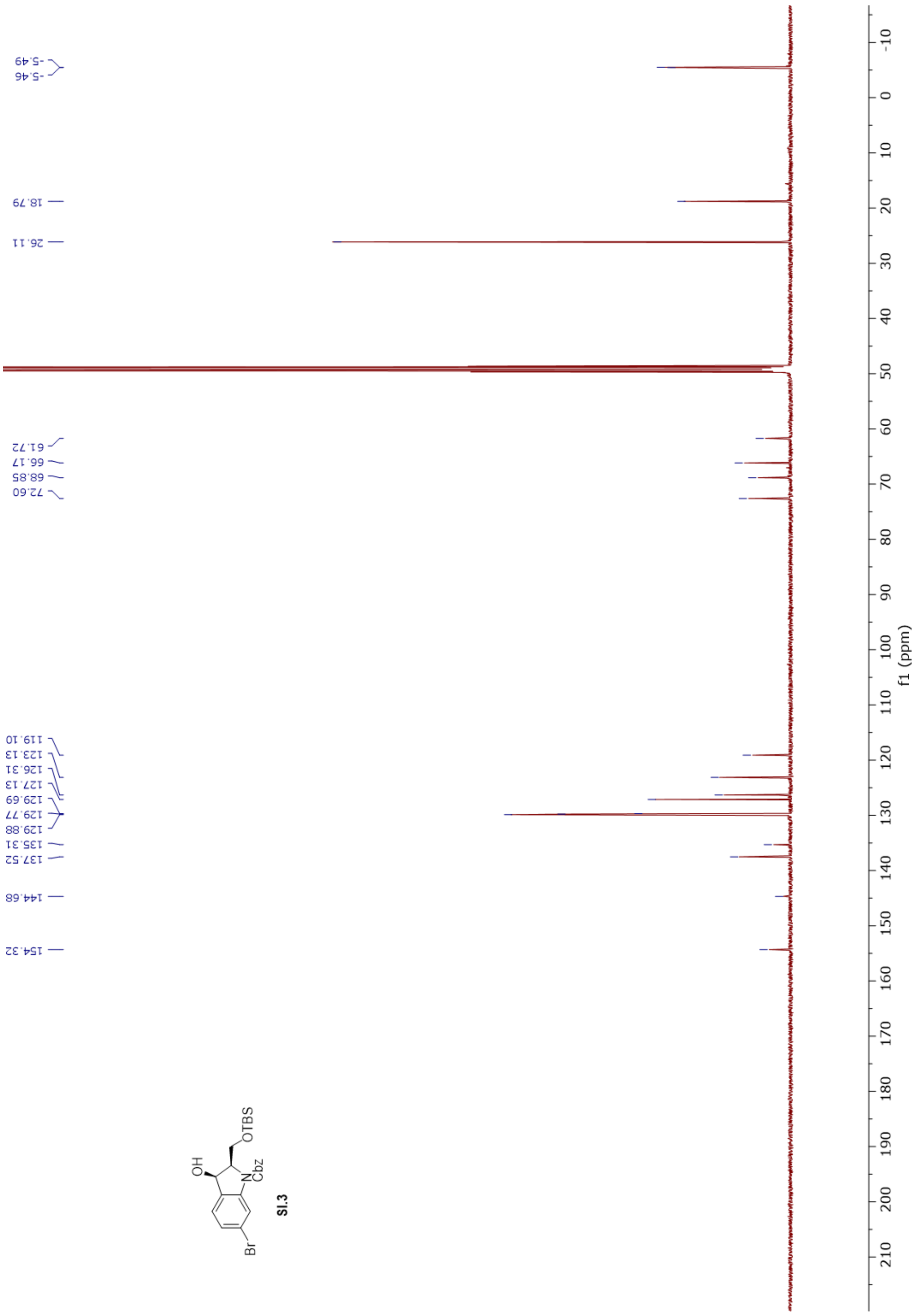
-5.49
-5.46
18.79
26.11
61.72
66.17
68.85
72.60
119.10
123.13
126.31
127.13
129.69
129.77
129.88
135.31
137.52
144.68
154.32
f1 (ppm)
OH
OTBS
N
Cbz
Br
SI.3

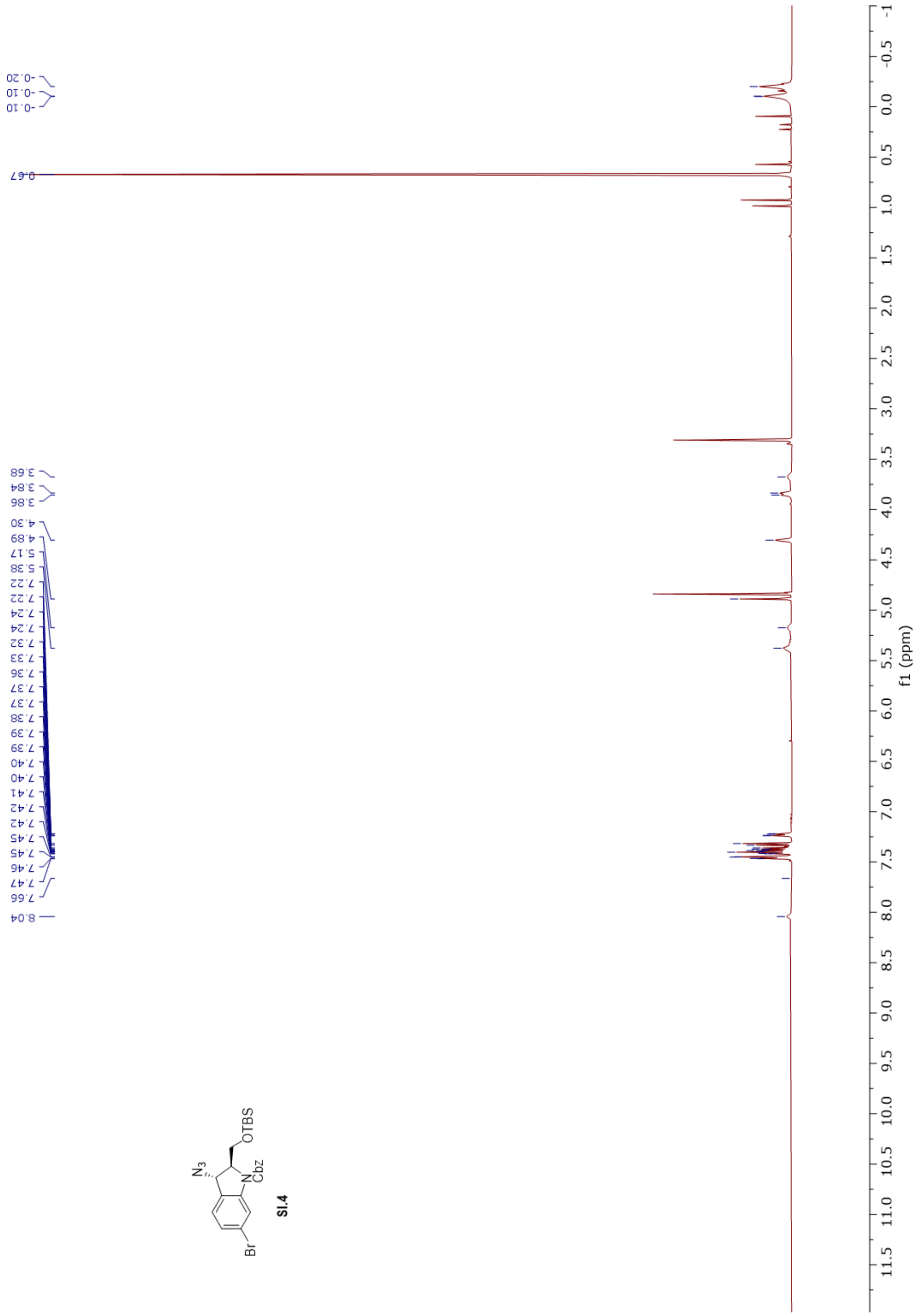
-0.20
-0.10
-0.10
0.67
3.68
3.84
3.86
4.30
4.89
5.17
5.38
7.22
7.22
7.24
7.24
7.32
7.33
7.36
7.37
7.37
7.38
7.39
7.39
7.40
7.40
7.41
7.42
7.42
7.45
7.45
7.46
7.47
7.66
8.04
N3
OTBS
N
Cbz
Br
SI.4
f1 (ppm)

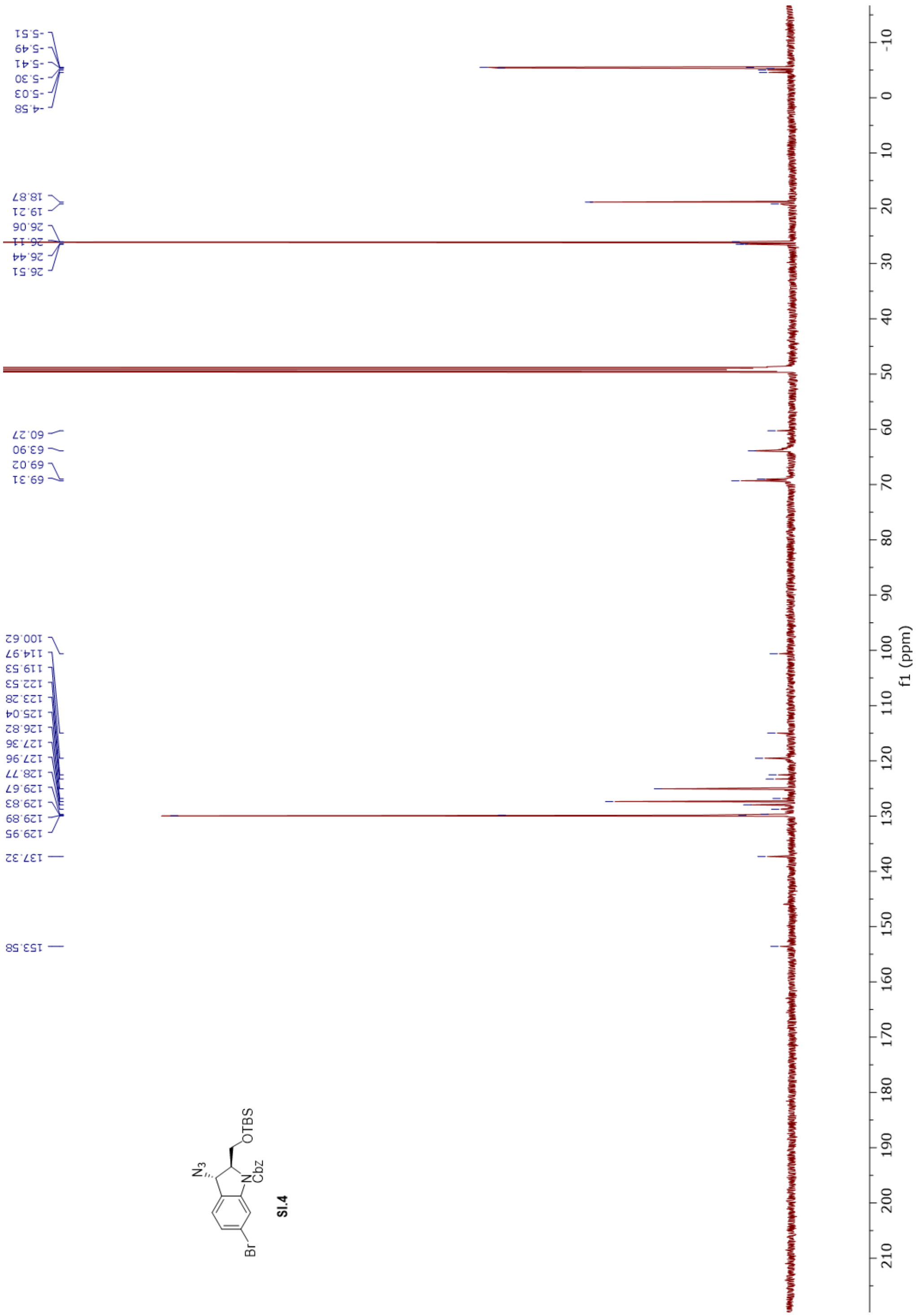
-5.51
-5.49
-5.41
-5.30
-5.03
-4.58
18.87
19.21
26.06
26.11
26.44
26.51
60.27
63.90
69.02
69.31
100.62
114.97
119.53
122.53
123.28
125.04
126.82
127.36
127.96
128.77
129.67
129.83
129.89
129.95
137.32
153.58
N3
OTBS
N
Cbz
Br
f1 (ppm)

SI.4

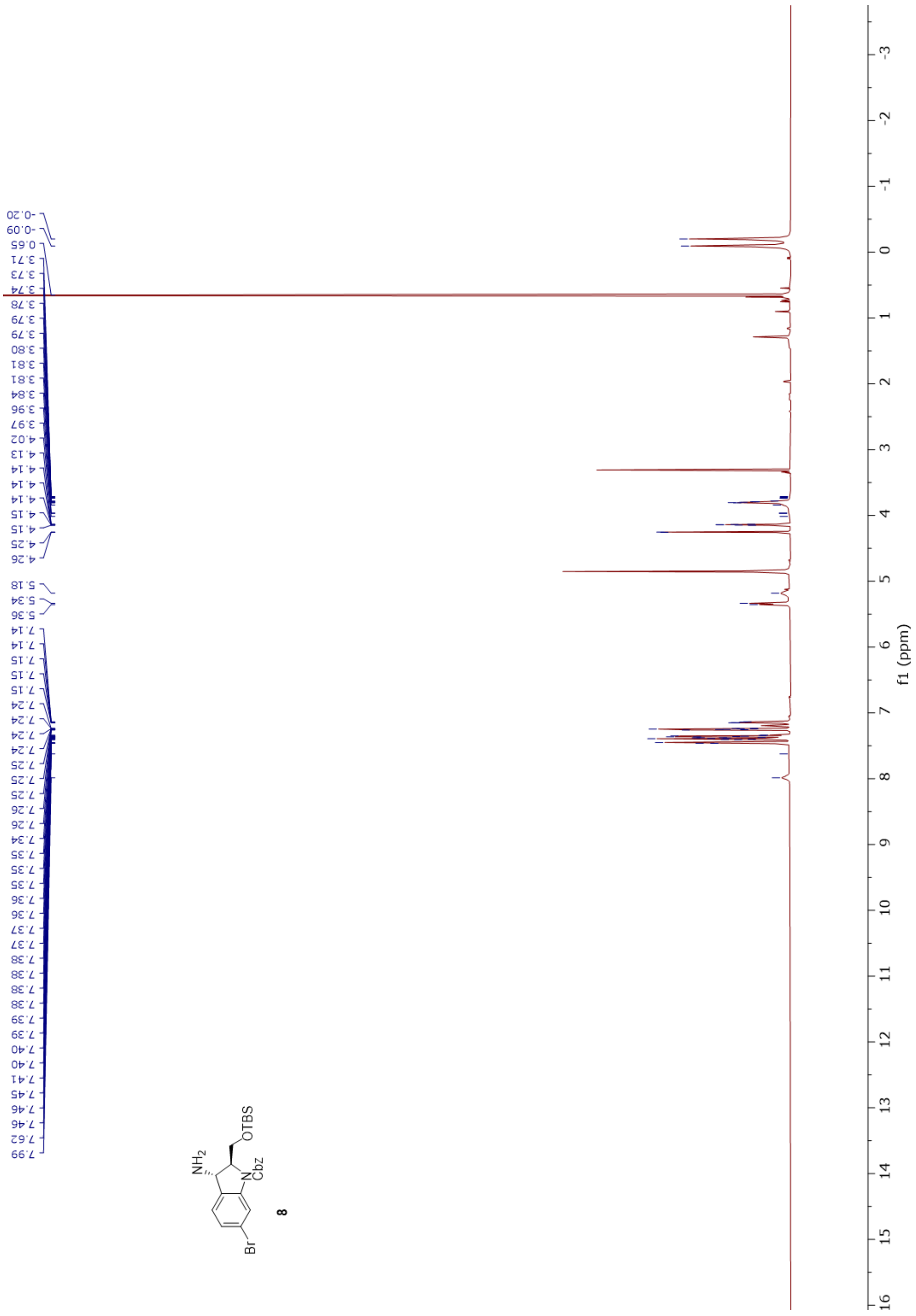

f1 (ppm)

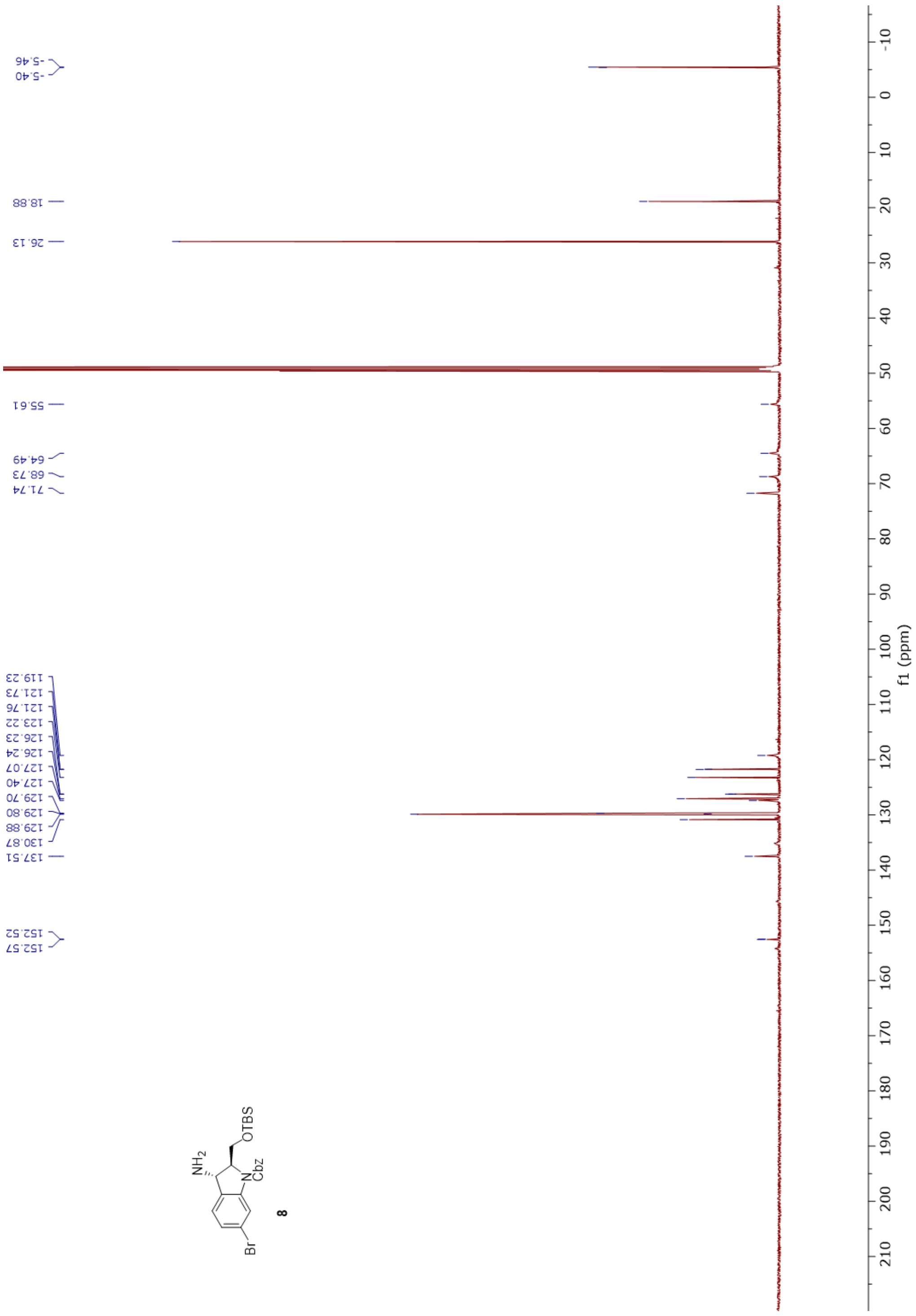
-5.46
-5.40
18.88
26.13
55.61
64.49
68.73
71.74
119.23
121.73
121.76
123.22
126.23
126.24
127.07
127.40
129.70
129.80
129.88
130.87
137.51
152.52
152.57
NH2
OTBS
N
Cbz
Br
8
f1 (ppm)

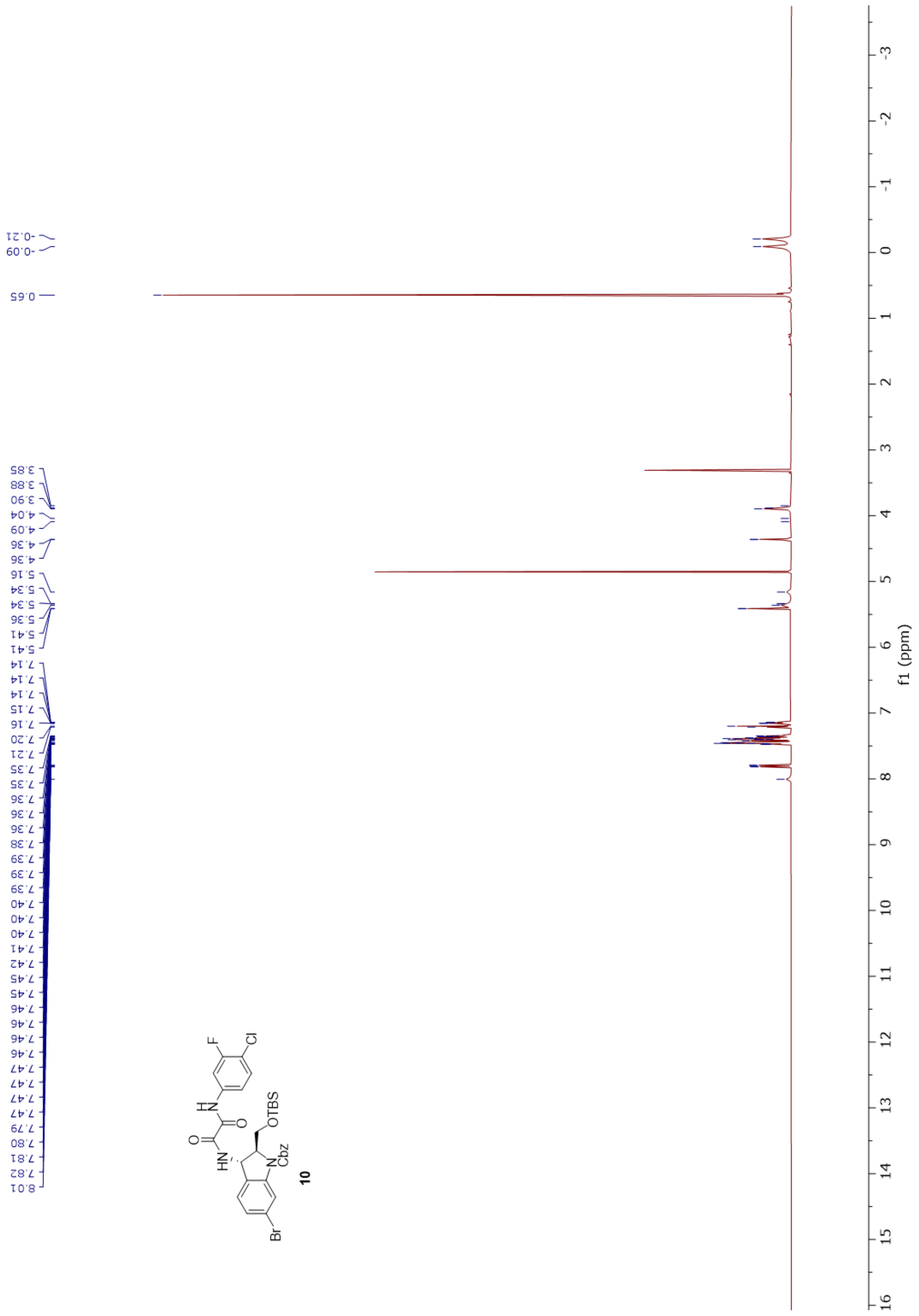

-0.21
-0.09
0.65
3.85
3.88
3.90
4.04
4.09
4.36
4.36
5.16
5.34
5.34
5.36
5.41
5.41
7.14
7.14
7.14
7.15
7.16
7.20
7.21
7.35
7.35
7.36
7.36
7.36
7.38
7.39
7.39
7.39
7.40
7.40
7.40
7.41
7.42
7.45
7.45
7.46
7.46
7.46
7.46
7.47
7.47
7.47
7.47
7.79
7.80
7.81
7.82
8.01
F
Cl
H
N
O
HN
OTBS
N
Cbz
Br
10
f1 (ppm)

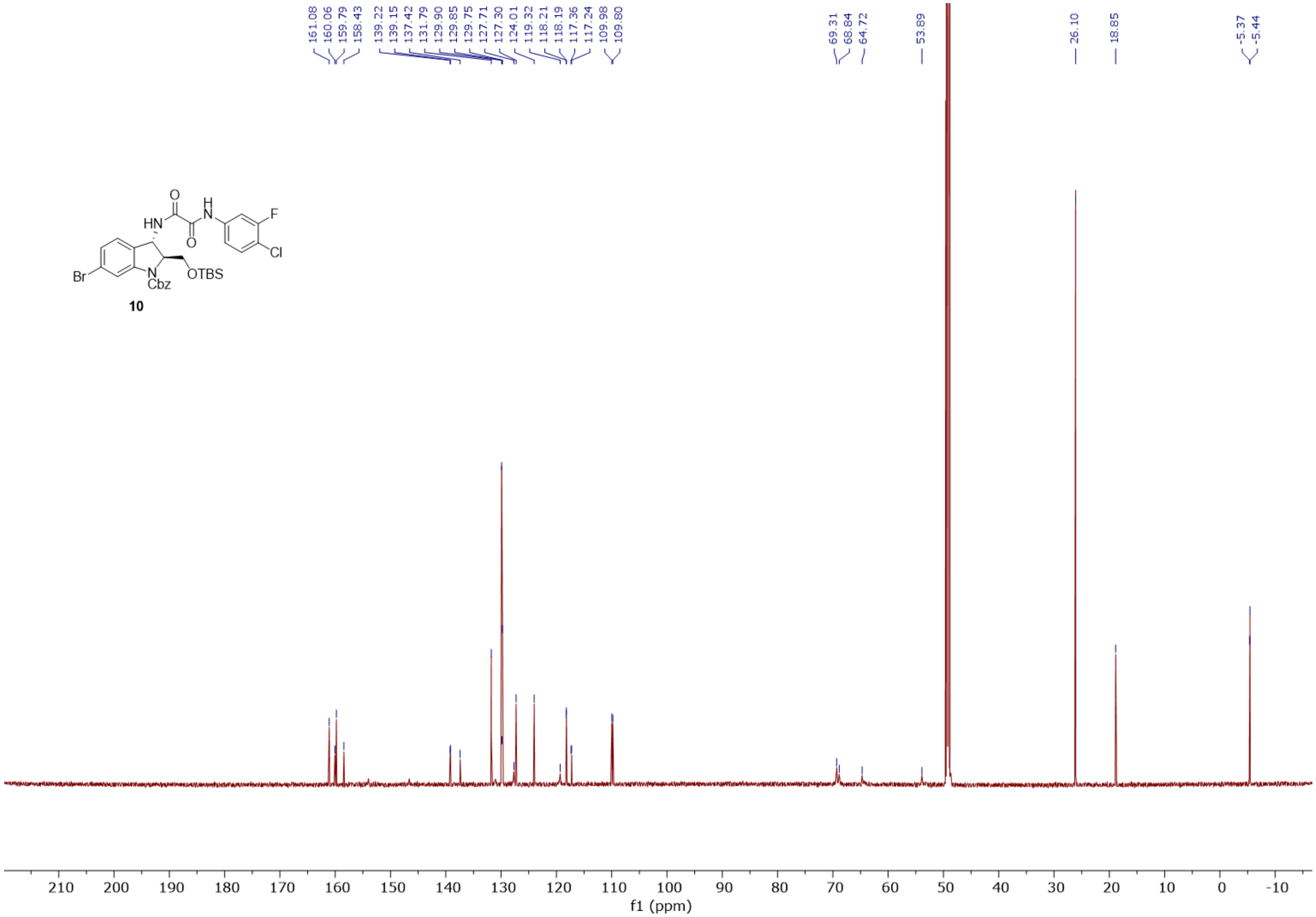

10
161.08
160.06
159.79
158.43
139.22
139.15
137.42
131.79
129.90
129.85
129.75
127.71
127.30
124.01
119.32
118.21
118.19
117.36
117.24
109.98
109.80
69.31
68.84
64.72
53.89
26.10
18.85
-5.37
-5.44
210
200
190
180
170
160
150
140
130
120
110
100
90
80
70
60
50
40
30
20
10
0
-10
f1 (ppm)

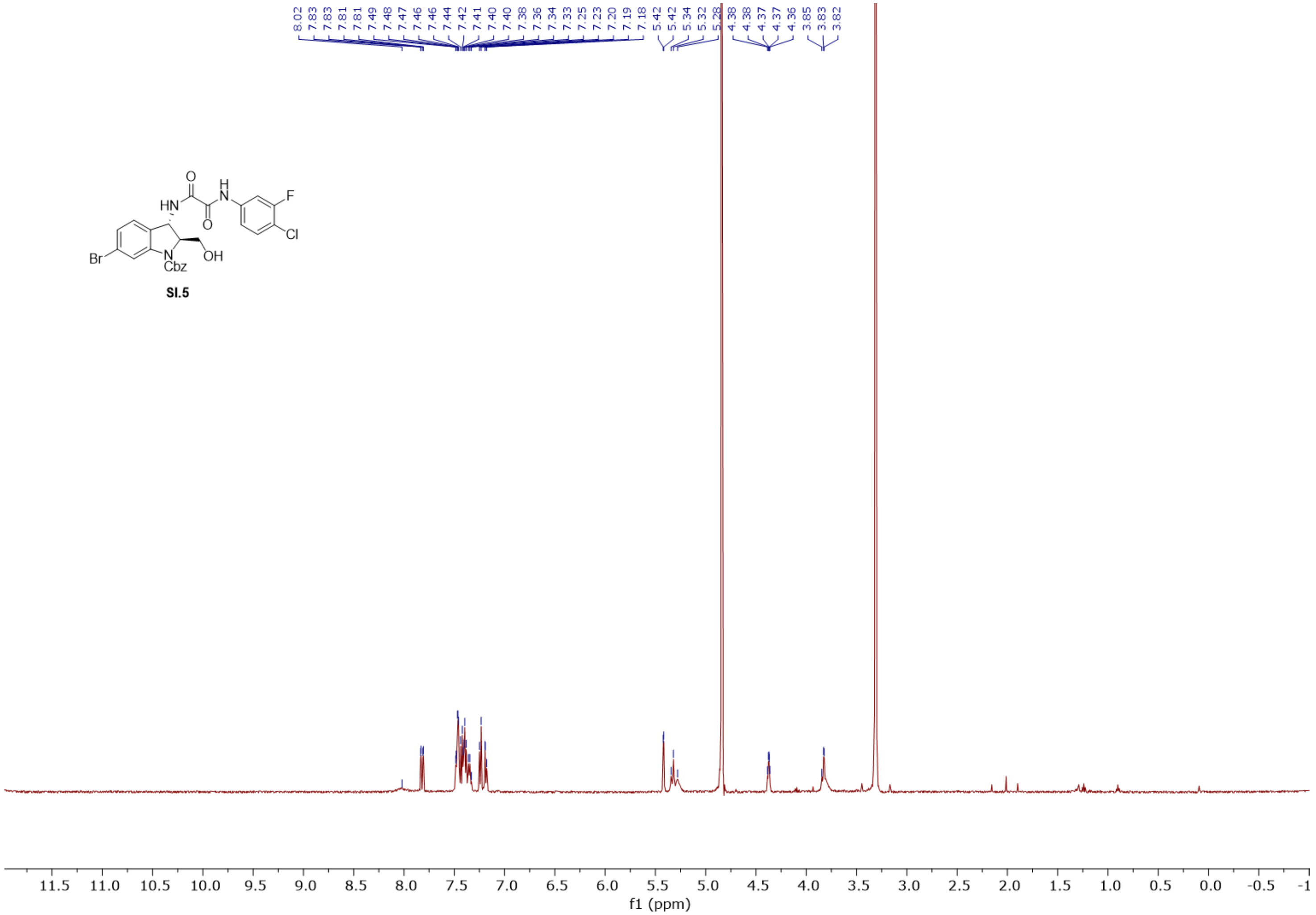
8.02 7.83 7.83 7.81 7.81 7.49 7.48 7.47 7.46 7.46 7.44 7.42 7.41 7.40 7.40 7.38 7.36 7.34 7.33 7.25 7.23 7.20 7.19 7.18
5.42 5.42 5.34 5.32 5.28
4.38 4.38 4.37 4.37 4.36
3.85 3.83 3.82
HN
O
N
H
O
F
Cl
Br
N
Cbz
OH
SI.5
11.5 11.0 10.5 10.0 9.5 9.0 8.5 8.0 7.5 7.0 6.5 6.0 5.5 5.0 4.5 4.0 3.5 3.0 2.5 2.0 1.5 1.0 0.5 0.0 -0.5 -1
f1 (ppm)

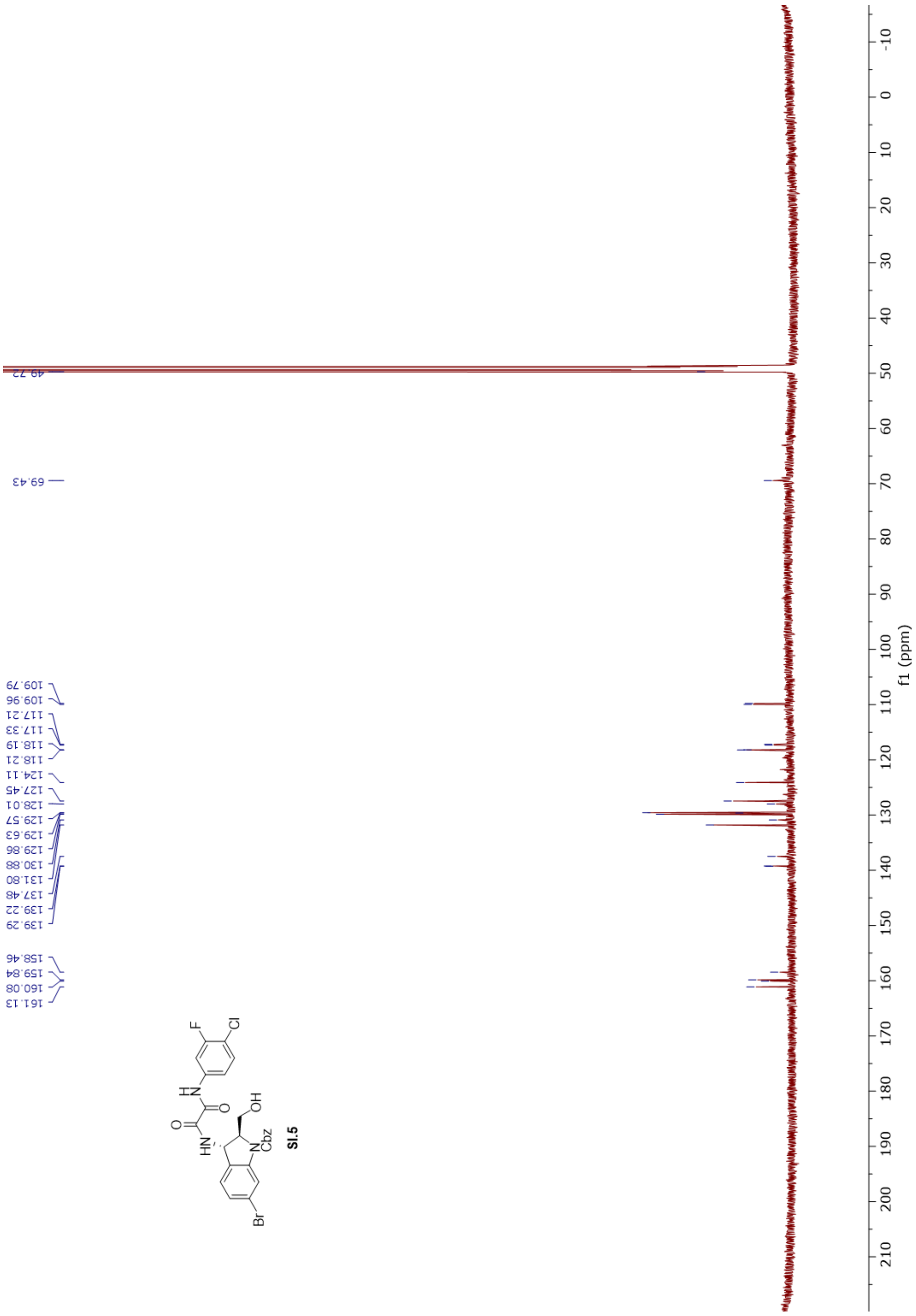
49.72
69.43
109.79
109.96
117.21
117.33
118.19
118.21
124.11
127.45
128.01
129.57
129.63
129.86
130.88
131.80
137.48
139.22
139.29
158.46
159.84
160.08
161.13
F
Cl
H
N
O
HN
OH
N
Cbz
Br
f1 (ppm)

SI.5

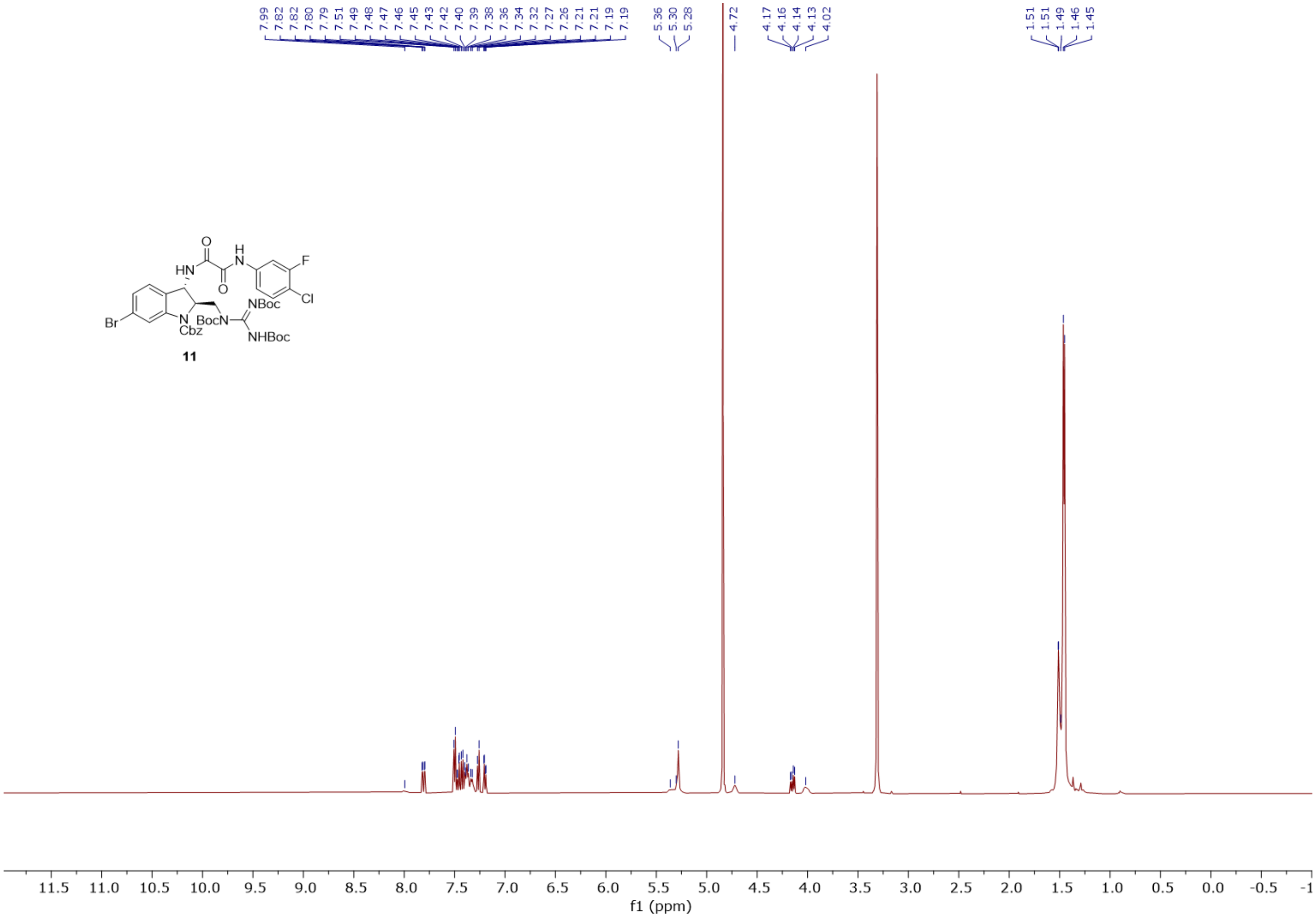
7.99
7.82
7.82
7.80
7.79
7.51
7.49
7.48
7.47
7.46
7.45
7.43
7.42
7.40
7.39
7.38
7.36
7.34
7.32
7.27
7.26
7.21
7.21
7.19
7.19
5.36
5.30
5.28
4.72
4.17
4.16
4.14
4.13
4.02
1.51
1.51
1.49
1.46
1.45
11.5 11.0 10.5 10.0 9.5 9.0 8.5 8.0 7.5 7.0 6.5 6.0 5.5 5.0 4.5 4.0 3.5 3.0 2.5 2.0 1.5 1.0 0.5 0.0 -0.5 -1
f1 (ppm)

**11**

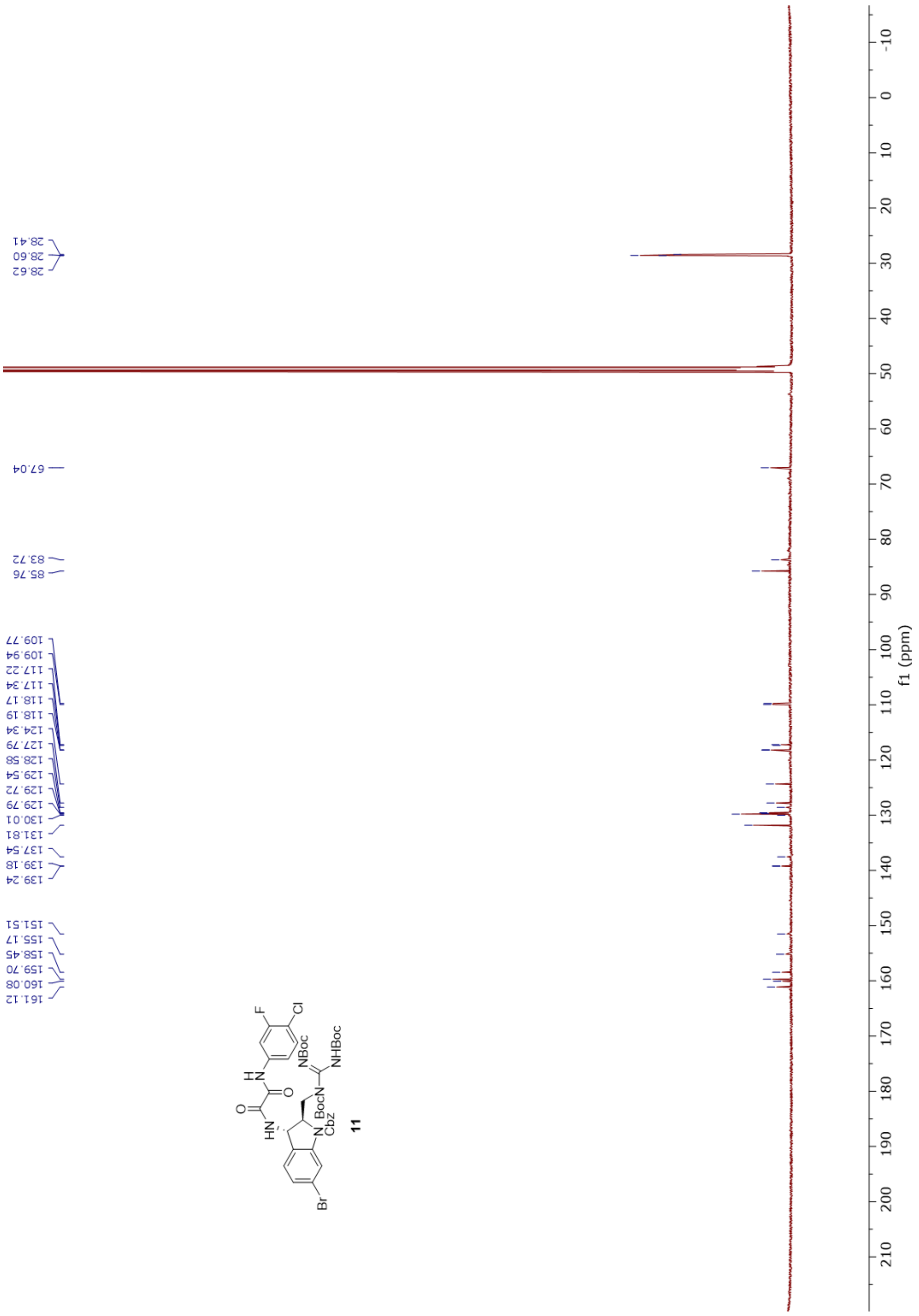
28.41
28.60
28.62
67.04
83.72
85.76
109.77
109.94
117.22
117.34
118.17
118.19
124.34
127.79
128.58
129.54
129.72
129.79
130.01
131.81
137.54
139.18
139.24
151.51
155.17
158.45
159.70
160.08
161.12
f1 (ppm)
F
Cl
NBoc
NHBoc
BocN
Cbz
Br
11

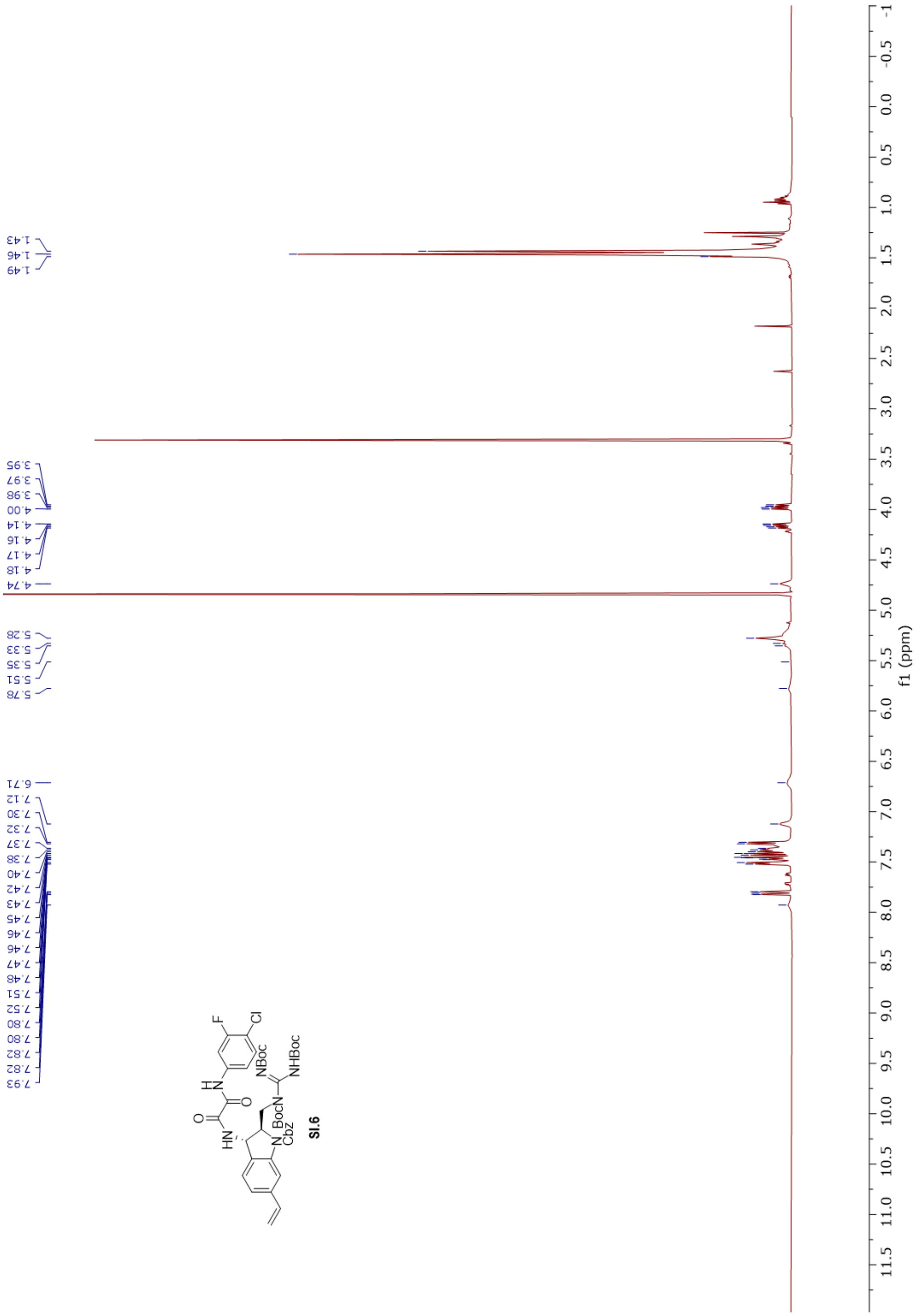

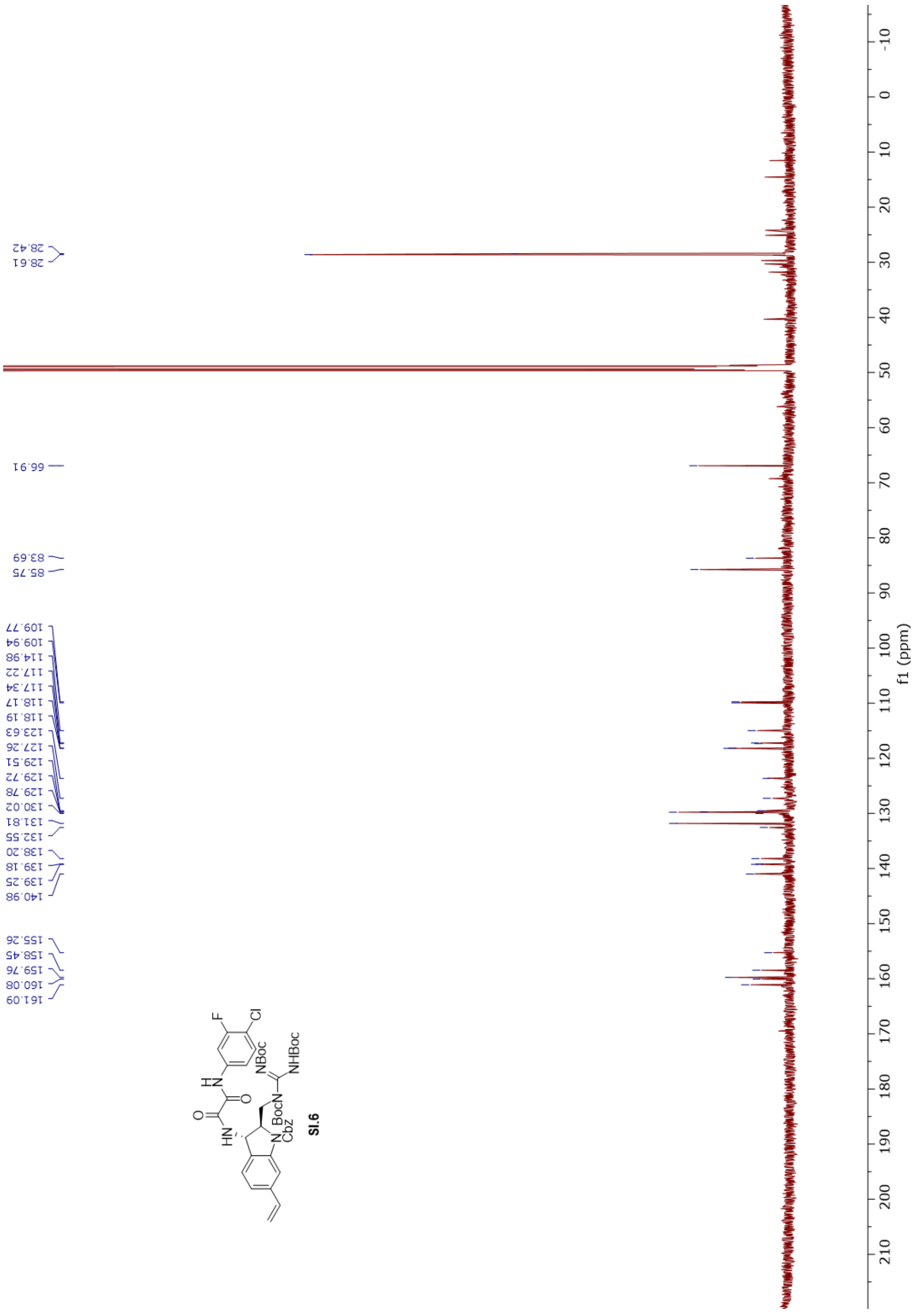
28.42
28.61
66.91
83.69
85.75
109.77
109.94
114.98
117.22
117.34
118.17
118.19
123.63
127.26
129.51
129.72
129.78
130.02
131.81
132.55
138.20
139.18
139.25
140.98
155.26
158.45
159.76
160.08
161.09
F
Cl
H
N
O
HN
NBoc
NHBoc
BocN
N
Cbz
SI.6
f1 (ppm)

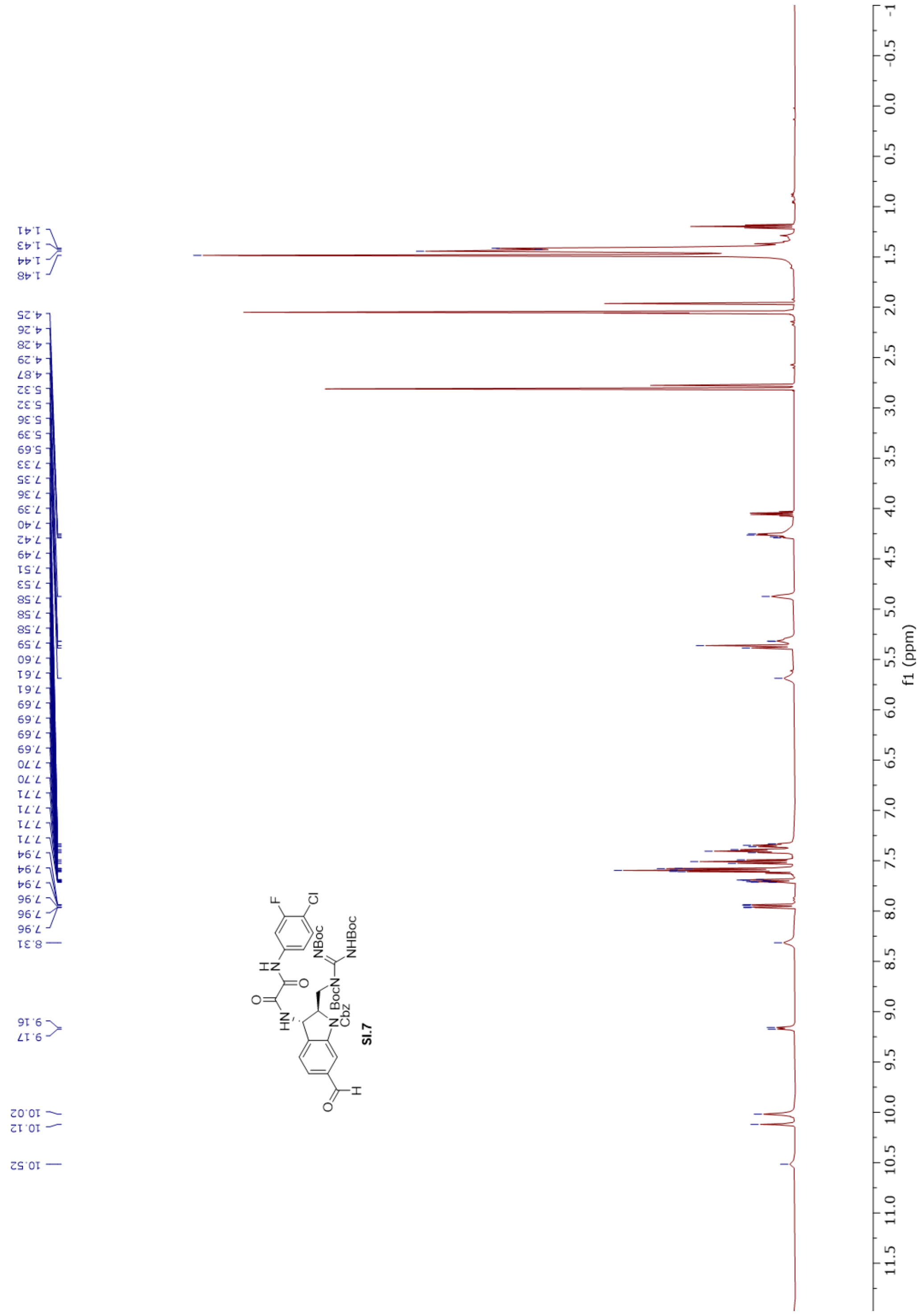
f1 (ppm)
SI.7
NBoc
NHBoc
BocN
Cbz
F
Cl

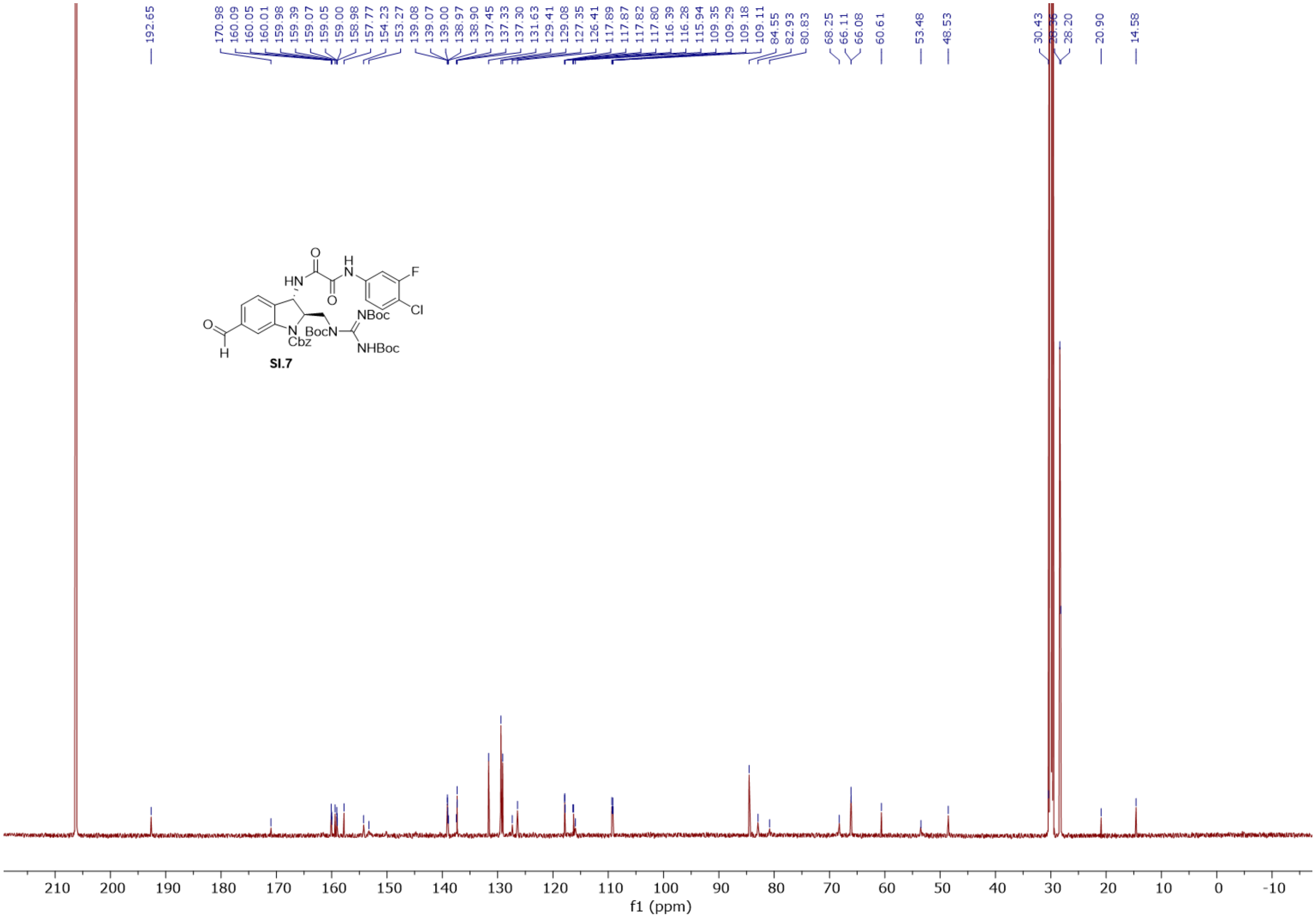
192.65
170.98
160.09
160.05
160.01
159.98
159.39
159.07
159.05
159.00
158.98
157.77
154.23
153.27
139.08
139.07
139.00
138.97
138.90
137.45
137.33
137.30
131.63
129.41
129.08
127.35
126.41
117.89
117.87
117.82
117.80
116.39
116.28
115.94
109.35
109.29
109.18
109.11
84.55
82.93
80.83
68.25
66.11
66.08
60.61
53.48
48.53
30.43
28.36
28.20
20.90
14.58
O
HN
H
N
F
Cl
O
NBoc
O
H
N
Cbz
BocN
NHBoc
SI.7
210 200 190 180 170 160 150 140 130 120 110 100 90 80 70 60 50 40 30 20 10 0 -10
f1 (ppm)

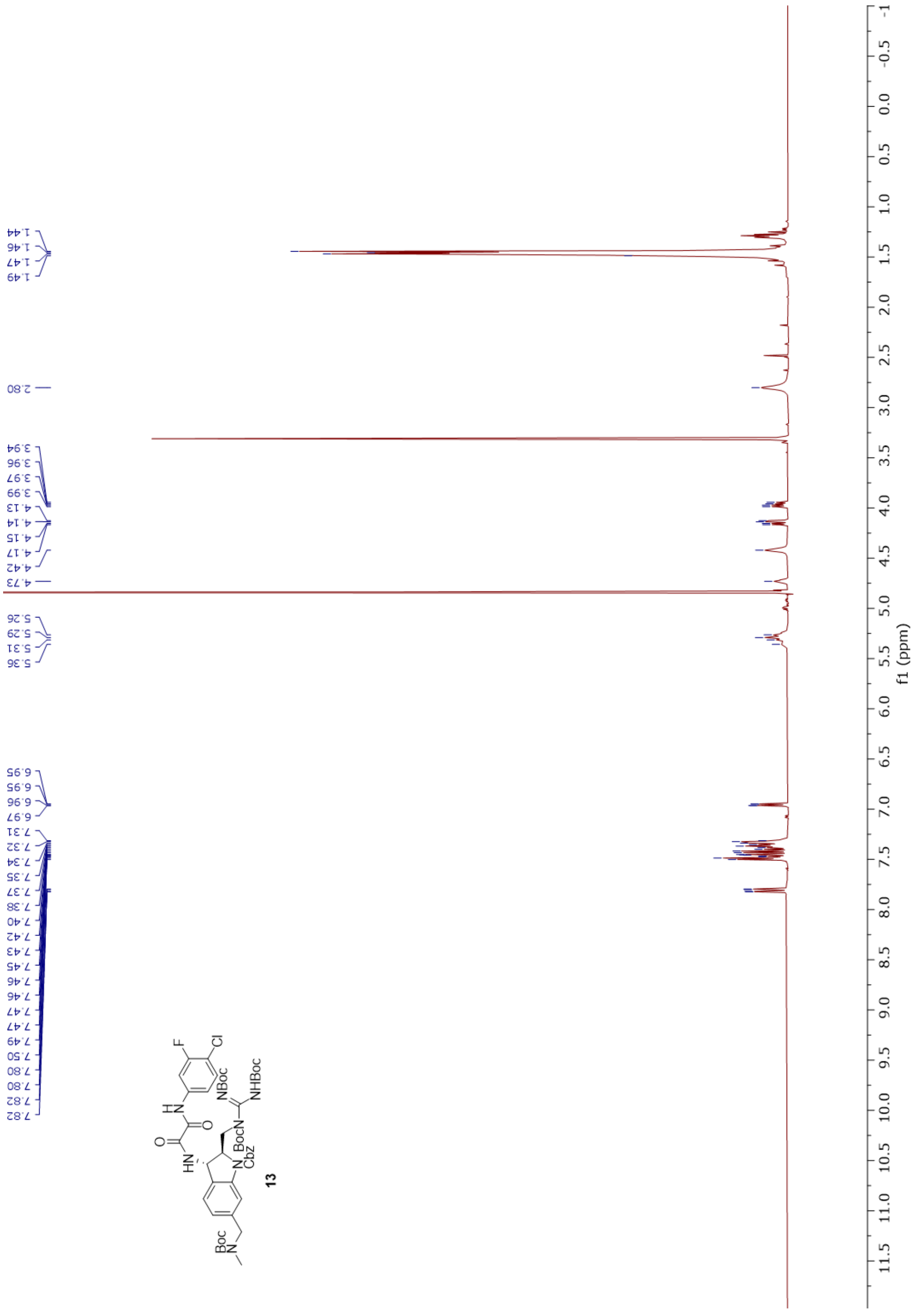

1.44
1.46
1.47
1.49
2.80
3.94
3.96
3.97
3.99
4.13
4.14
4.15
4.17
4.42
4.73
5.26
5.29
5.31
5.36
6.95
6.95
6.96
6.97
7.31
7.32
7.34
7.35
7.37
7.38
7.40
7.42
7.43
7.45
7.46
7.46
7.47
7.47
7.49
7.50
7.80
7.80
7.82
7.82
F
Cl
H
N
O
O
HN
NBoc
NHBoc
BocN
N
Cbz
Boc
N
13
f1 (ppm)

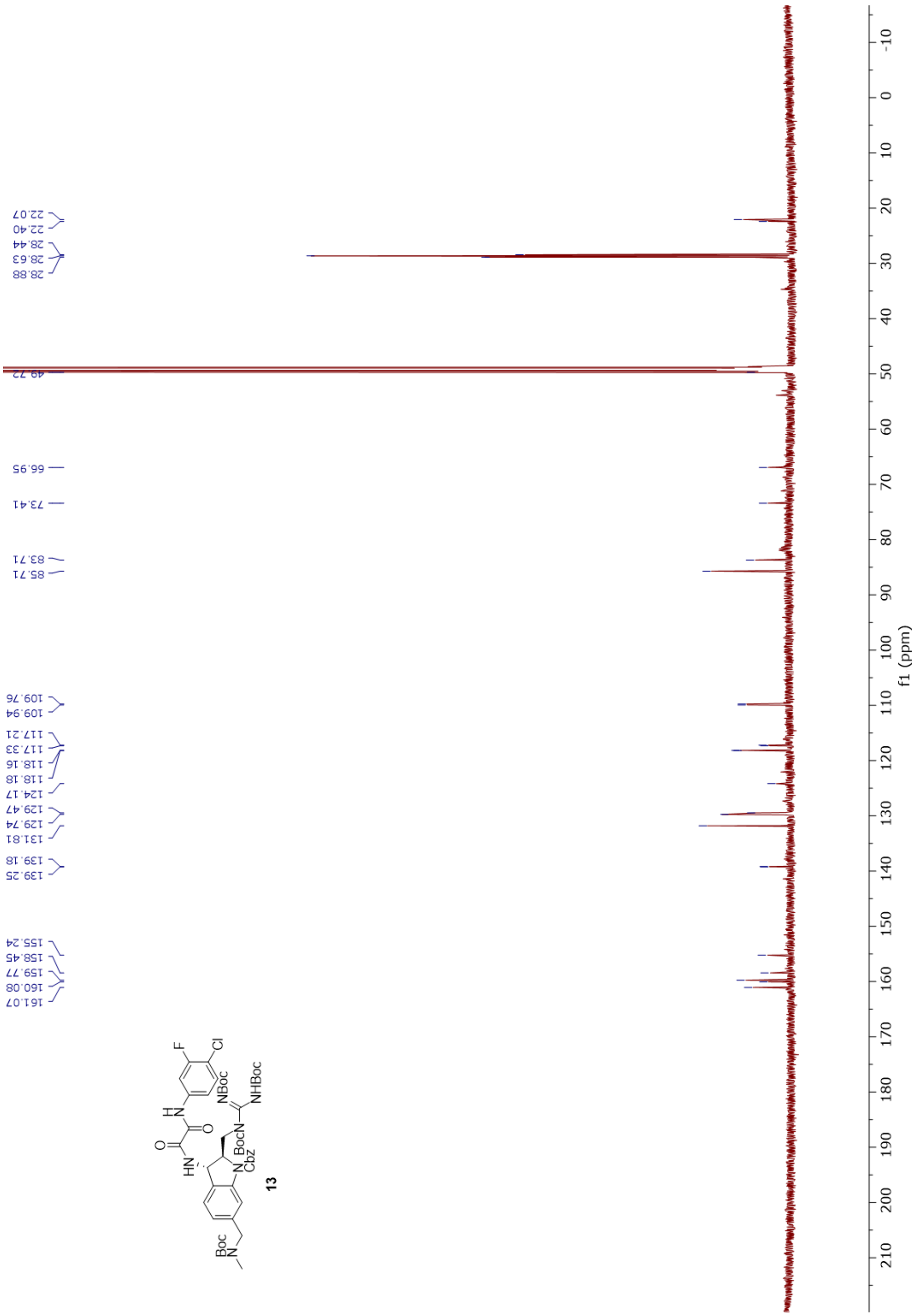

22.07
22.40
28.44
28.63
28.88
49.72
66.95
73.41
83.71
85.71
109.76
109.94
117.21
117.33
118.16
118.18
124.17
129.47
129.74
131.81
139.18
139.25
155.24
158.45
159.77
160.08
161.07
f1 (ppm)
13

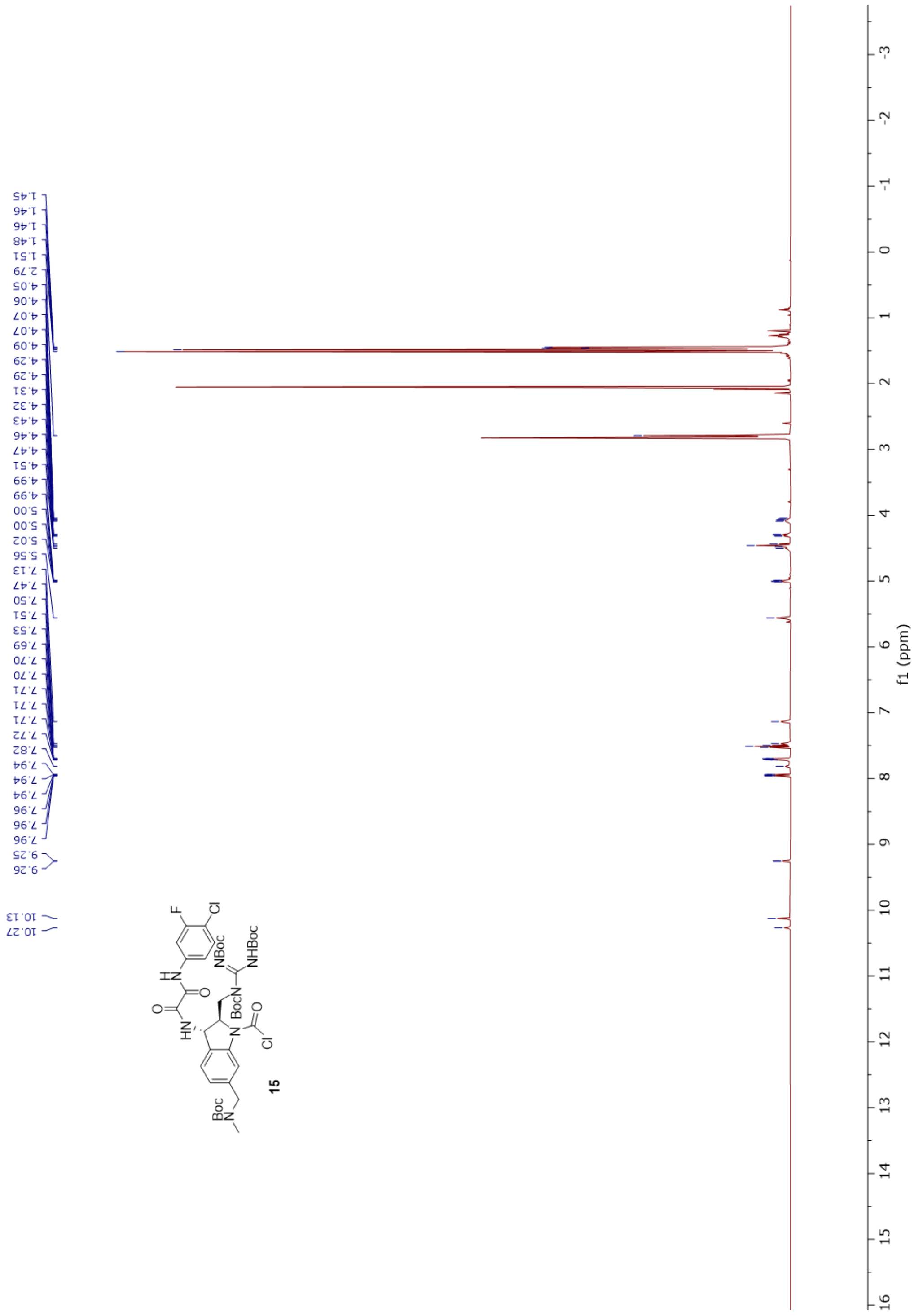
1.45
1.46
1.46
1.48
1.51
2.79
4.05
4.06
4.07
4.07
4.09
4.29
4.29
4.31
4.32
4.43
4.46
4.47
4.51
4.99
4.99
5.00
5.00
5.02
5.56
7.13
7.47
7.50
7.51
7.53
7.69
7.70
7.70
7.71
7.71
7.71
7.72
7.82
7.94
7.94
7.94
7.96
7.96
7.96
9.25
9.26
10.13
10.27
F
Cl
H
N
O
HN
NBoc
NHBoc
BocN
N
Cl
Boc
15
f1 (ppm)

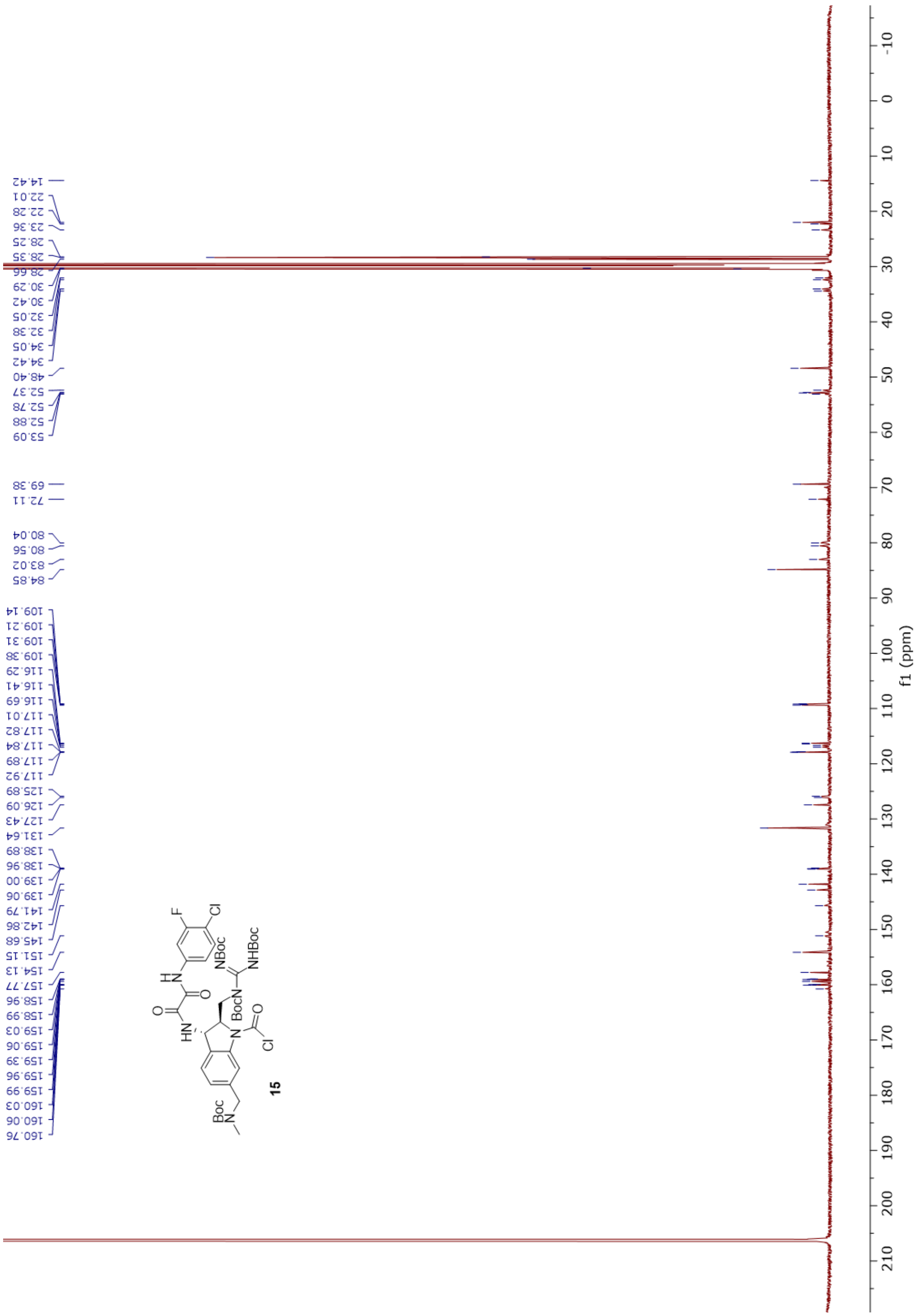
14.42
22.01
22.28
23.36
28.25
28.35
28.66
30.29
30.42
32.05
32.38
34.05
34.42
48.40
52.37
52.78
52.88
53.09
69.38
72.11
80.04
80.56
83.02
84.85
109.14
109.21
109.31
109.38
116.29
116.41
116.69
117.01
117.82
117.84
117.89
117.92
125.89
126.09
127.43
131.64
138.89
138.96
139.00
139.06
141.79
142.86
145.68
151.15
154.13
157.77
158.96
158.99
159.03
159.06
159.39
159.96
159.99
160.03
160.06
160.76
F
Cl
NBoc
NHBoc
BocN
HN
Boc
N
15
f1 (ppm)

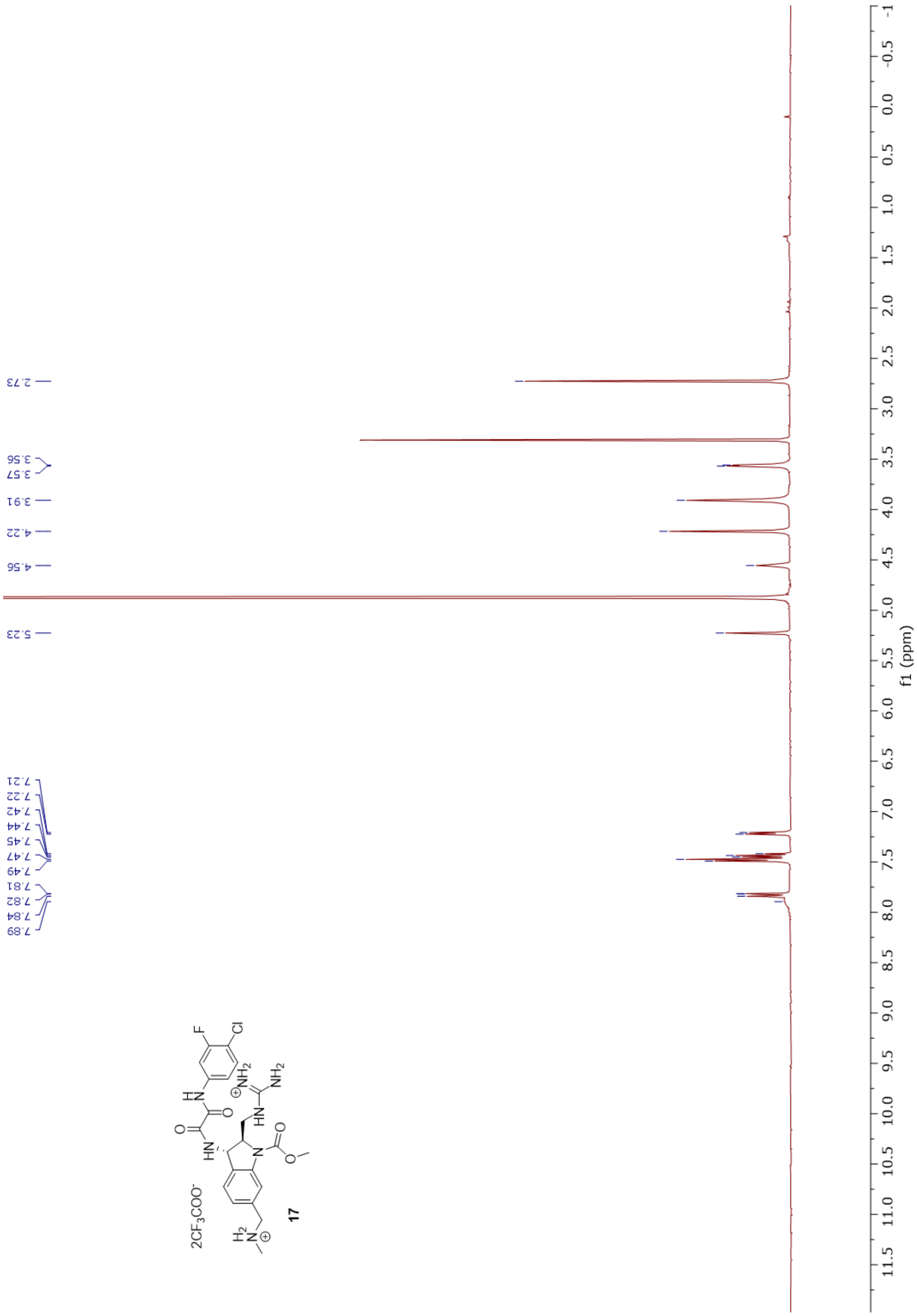
2.73
3.56
3.57
3.91
4.22
4.56
5.23
7.21
7.22
7.42
7.44
7.45
7.47
7.49
7.81
7.82
7.84
7.89
2CF3COO-
17
f1 (ppm)

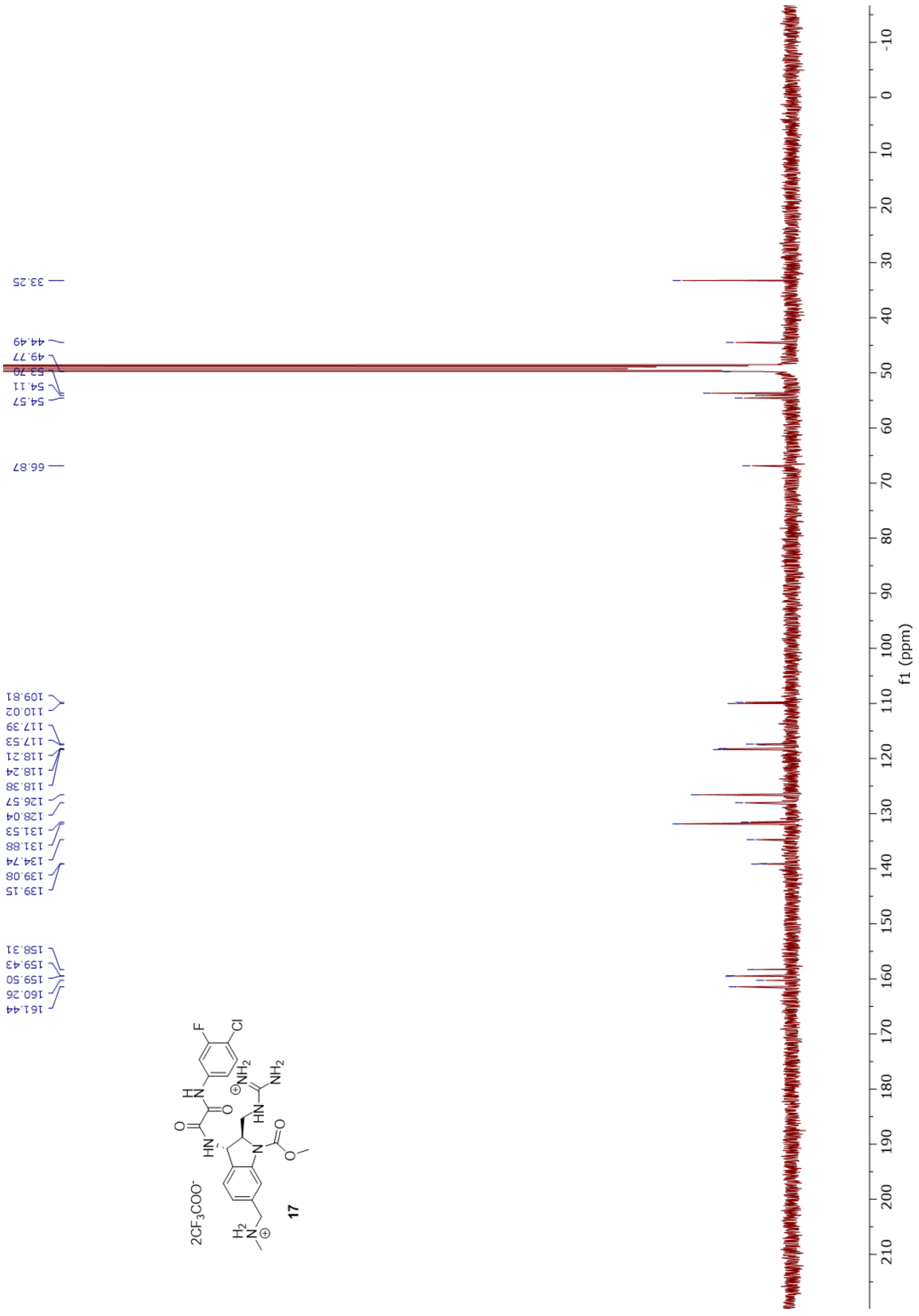

33.25
44.49
49.77
53.70
54.11
54.57
66.87
109.81
110.02
117.39
117.53
118.21
118.24
118.38
126.57
128.04
131.53
131.88
134.74
139.08
139.15
158.31
159.43
159.50
160.26
161.44
f1 (ppm)
2CF3COO-
17

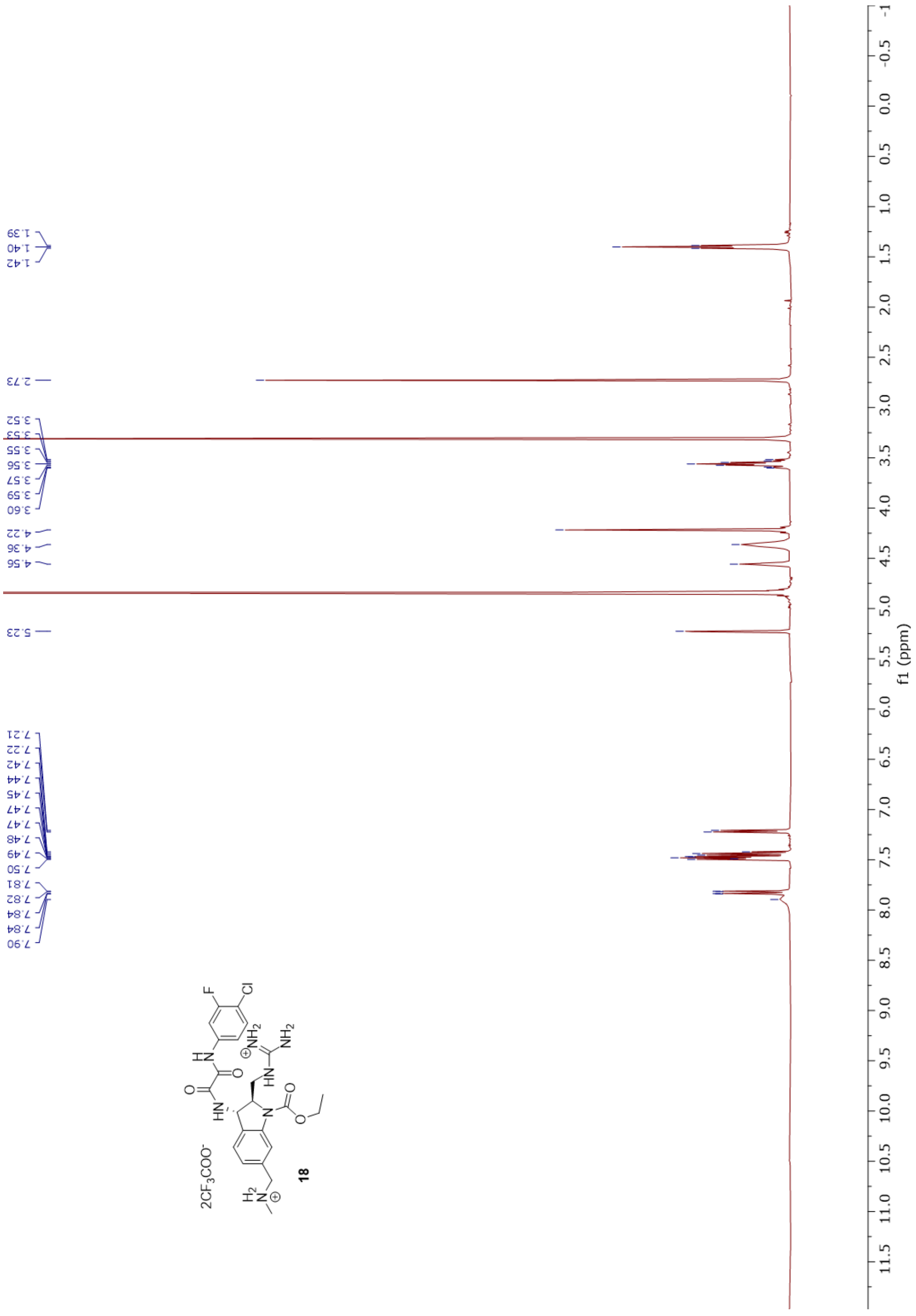
1.39
1.40
1.42
2.73
3.52
3.53
3.55
3.56
3.57
3.59
3.60
4.22
4.36
4.56
5.23
7.21
7.22
7.42
7.44
7.45
7.47
7.47
7.48
7.49
7.50
7.81
7.82
7.84
7.84
7.90
2CF3COO-
18
f1 (ppm)

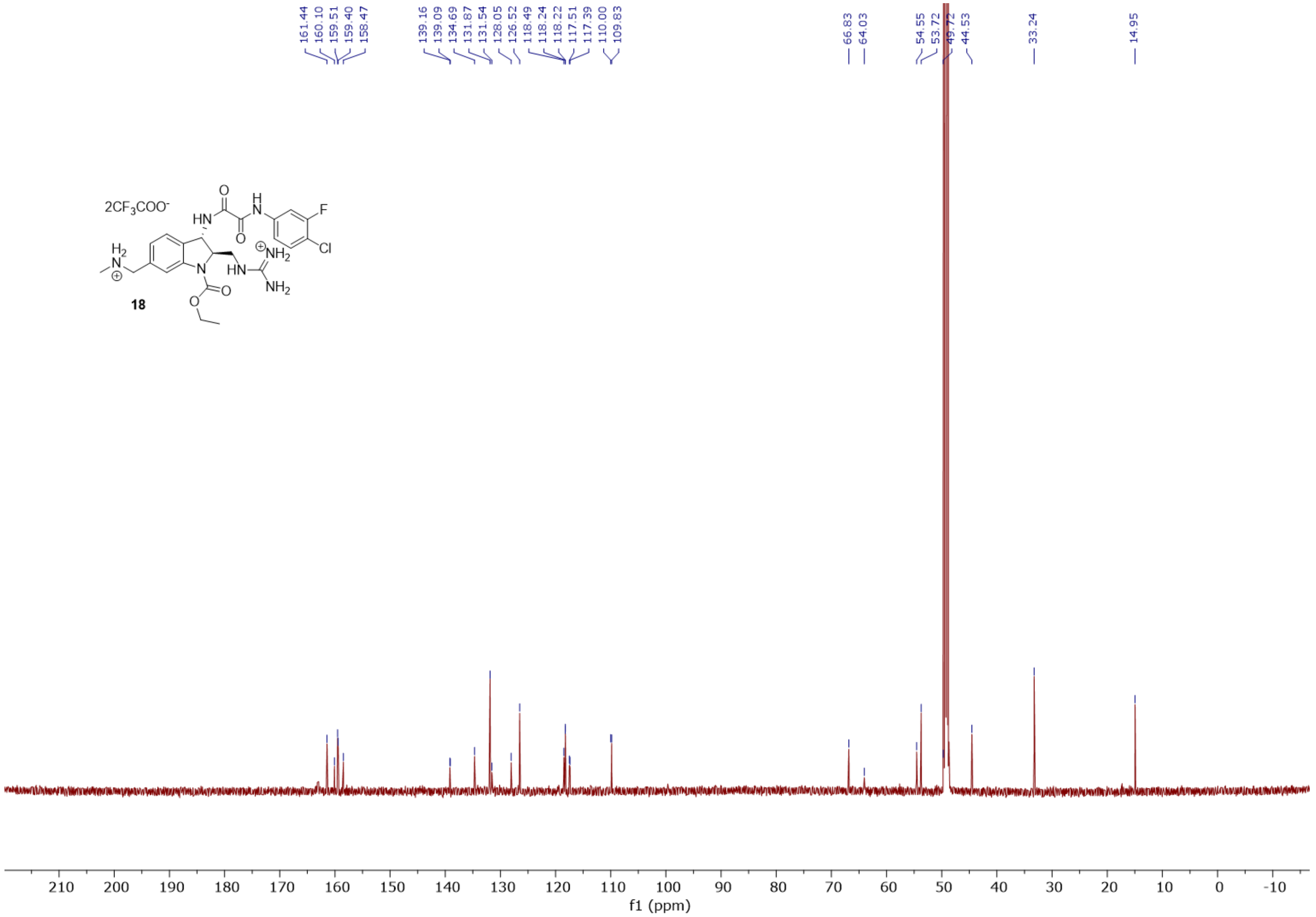
161.44
160.10
159.51
159.40
158.47
139.16
139.09
134.69
131.87
131.54
128.05
126.52
118.49
118.24
118.22
117.51
117.39
110.00
109.83
66.83
64.03
54.55
53.72
49.72
44.53
33.24
14.95
2CF3COO-
18
f1 (ppm)

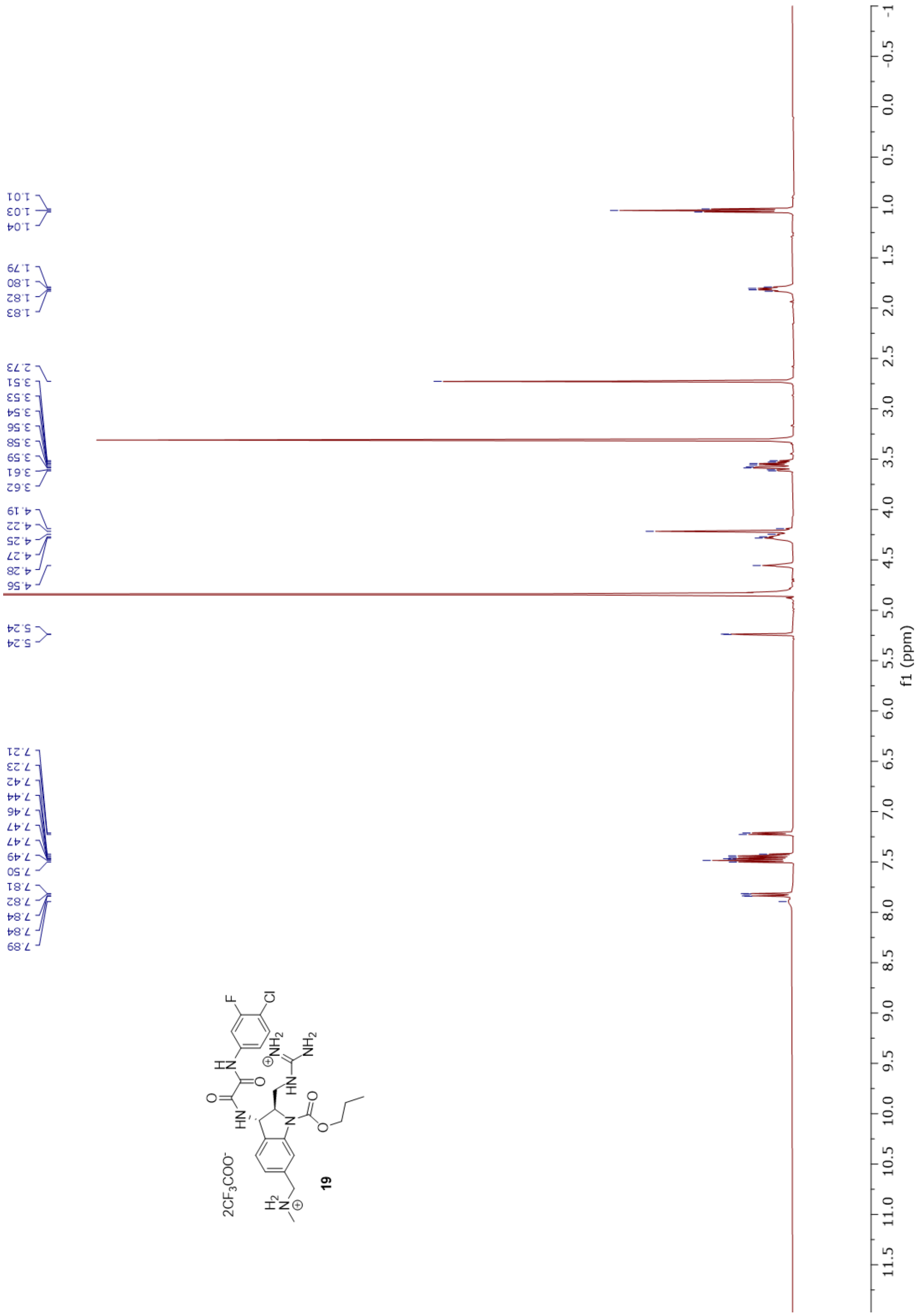
1.01
1.03
1.04
1.79
1.80
1.82
1.83
2.73
3.51
3.53
3.54
3.56
3.58
3.59
3.61
3.62
4.19
4.22
4.25
4.27
4.28
4.56
5.24
5.24
7.21
7.23
7.42
7.44
7.46
7.47
7.47
7.49
7.50
7.81
7.82
7.84
7.84
7.89
2CF3COO-
19
f1 (ppm)

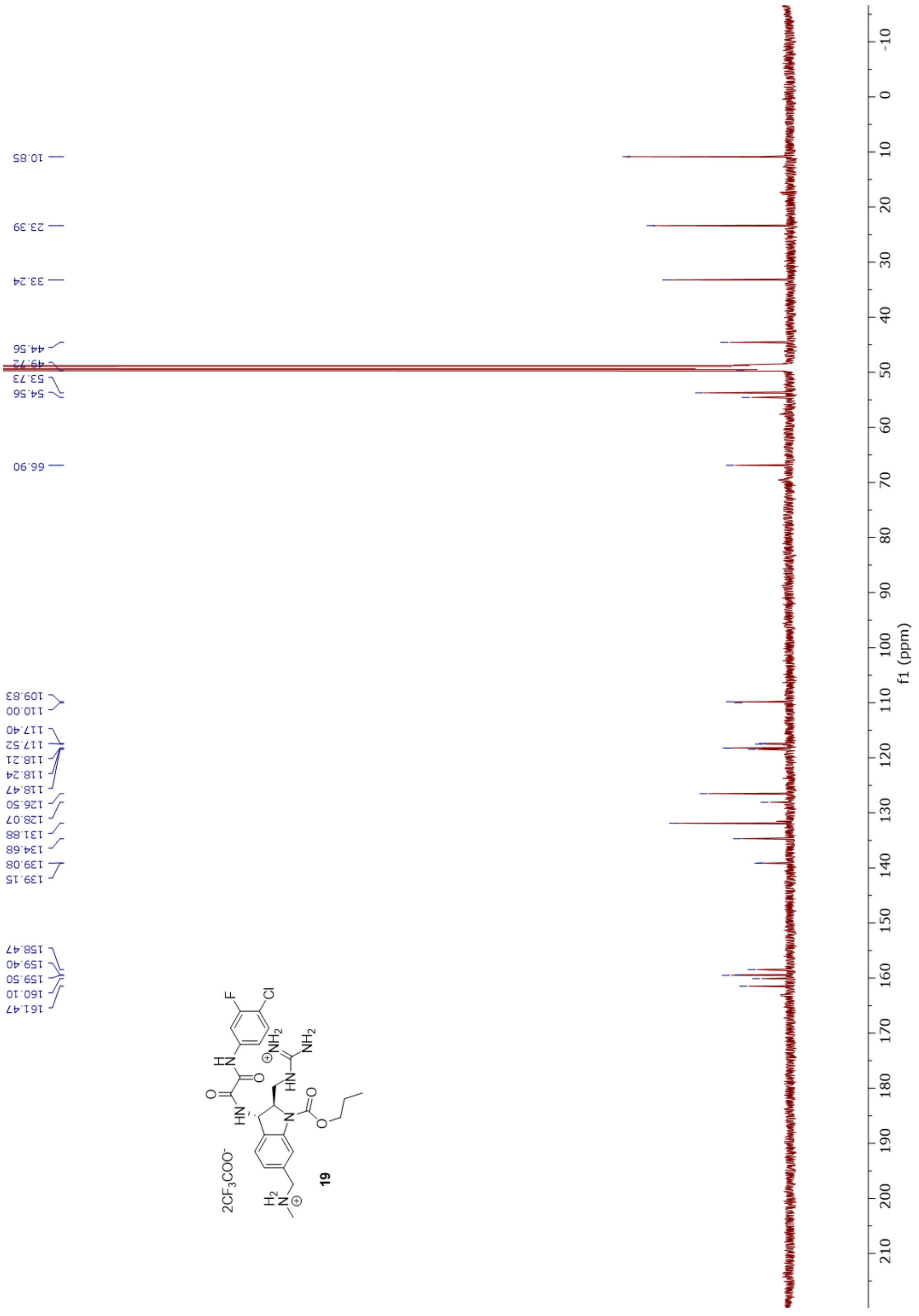
10.85
23.39
33.24
44.56
49.72
53.73
54.56
66.90
109.83
110.00
117.40
117.52
118.21
118.24
118.47
126.50
128.07
131.88
134.68
139.08
139.15
158.47
159.40
159.50
160.10
161.47
f1 (ppm)
2CF3COO⁻
19

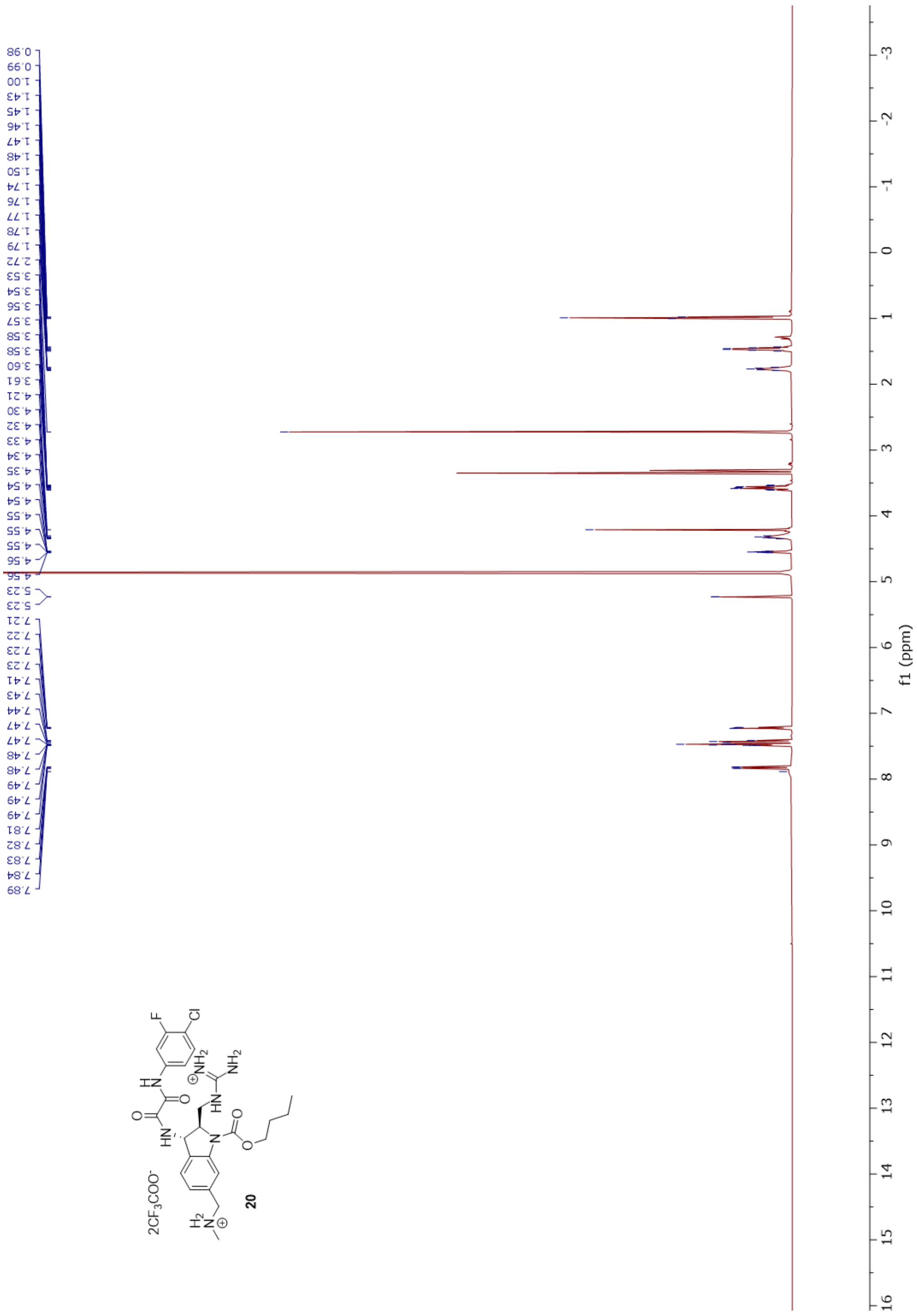
0.98 0.99 1.00 1.43 1.45 1.46 1.47 1.48 1.50 1.74 1.76 1.77 1.78 1.79 2.72 3.53 3.54 3.56 3.57 3.58 3.58 3.60 3.61 4.21 4.30 4.32 4.33 4.34 4.35 4.54 4.54 4.55 4.55 4.55 4.56 4.56 5.23 5.23 7.21 7.22 7.23 7.23 7.41 7.43 7.44 7.47 7.47 7.48 7.48 7.49 7.49 7.49 7.81 7.82 7.83 7.84 7.89
f1 (ppm)
2CF3COO-
20

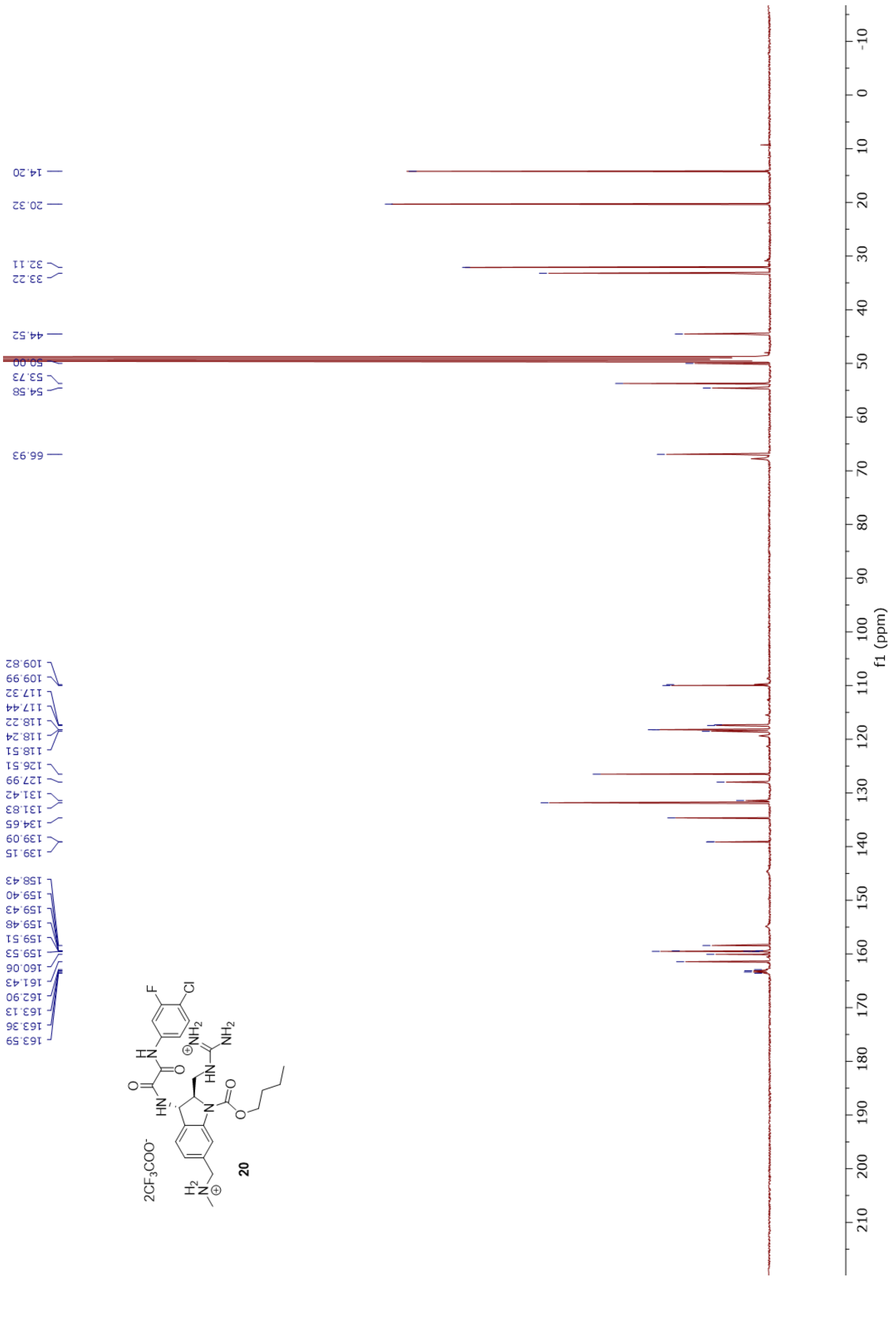

14.20
20.32
32.11
33.22
44.52
50.00
53.73
54.58
66.93
109.82
109.99
117.32
117.44
118.22
118.24
118.51
126.51
127.99
131.42
131.83
134.65
139.09
139.15
158.43
159.40
159.43
159.48
159.51
159.53
160.06
161.43
162.90
163.13
163.36
163.59
f1 (ppm)
20

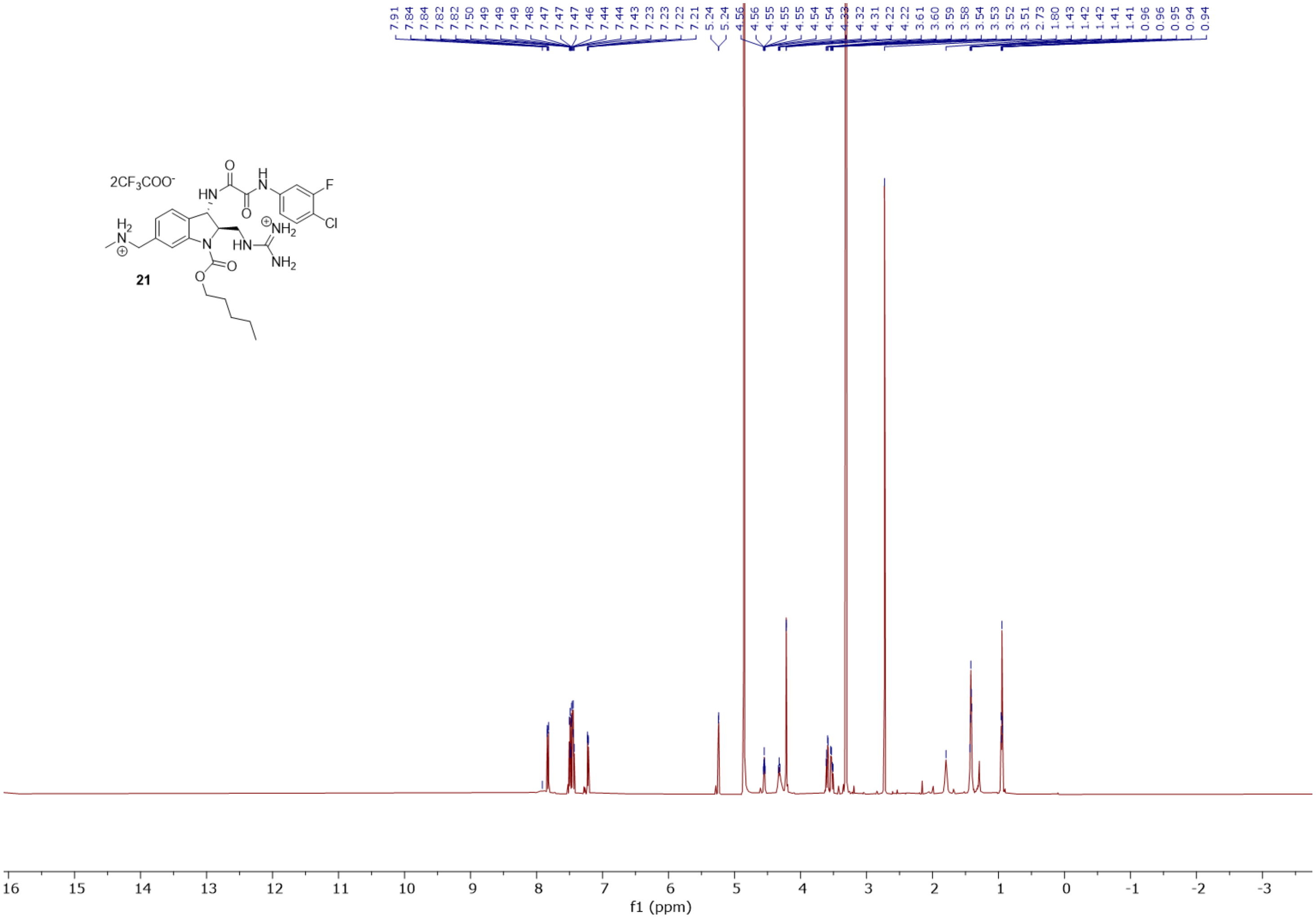
7.91 7.84 7.84 7.82 7.82 7.50 7.49 7.49 7.49 7.48 7.47 7.47 7.47 7.46 7.44 7.44 7.43 7.23 7.23 7.22 7.21 5.24 5.24 4.56 4.56 4.55 4.55 4.55 4.54 4.54 4.33 4.32 4.31 4.22 4.22 3.61 3.60 3.59 3.58 3.54 3.53 3.52 3.51 2.73 1.80 1.43 1.42 1.42 1.41 1.41 0.96 0.96 0.95 0.94 0.94
2CF3COO-
21
16 15 14 13 12 11 10 9 8 7 6 5 4 3 2 1 0 -1 -2 -3
f1 (ppm)

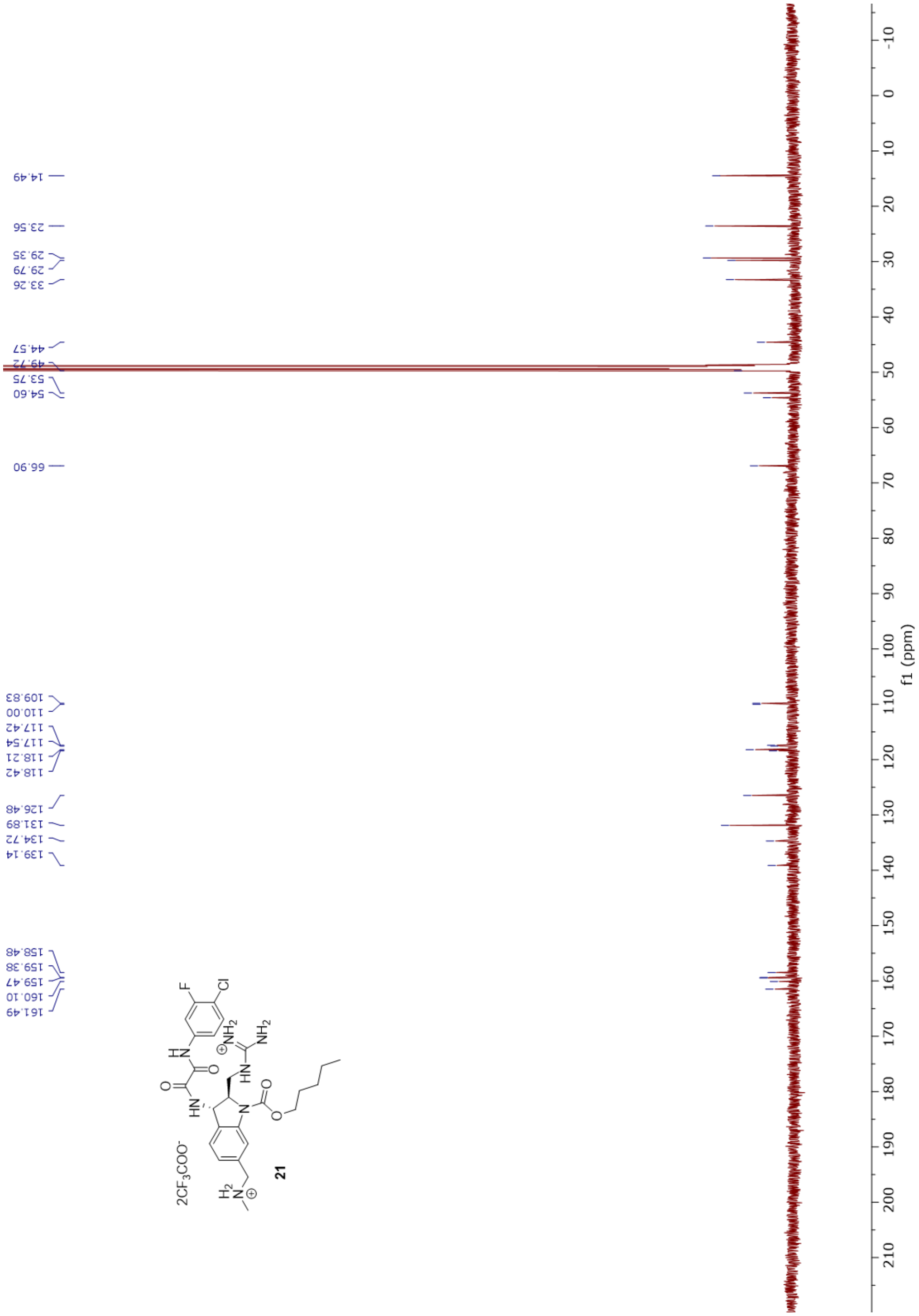
14.49
23.56
29.35
29.79
33.26
44.57
49.72
53.75
54.60
66.90
109.83
110.00
117.42
117.54
118.21
118.42
126.48
131.89
134.72
139.14
158.48
159.38
159.47
160.10
161.49
21
f1 (ppm)

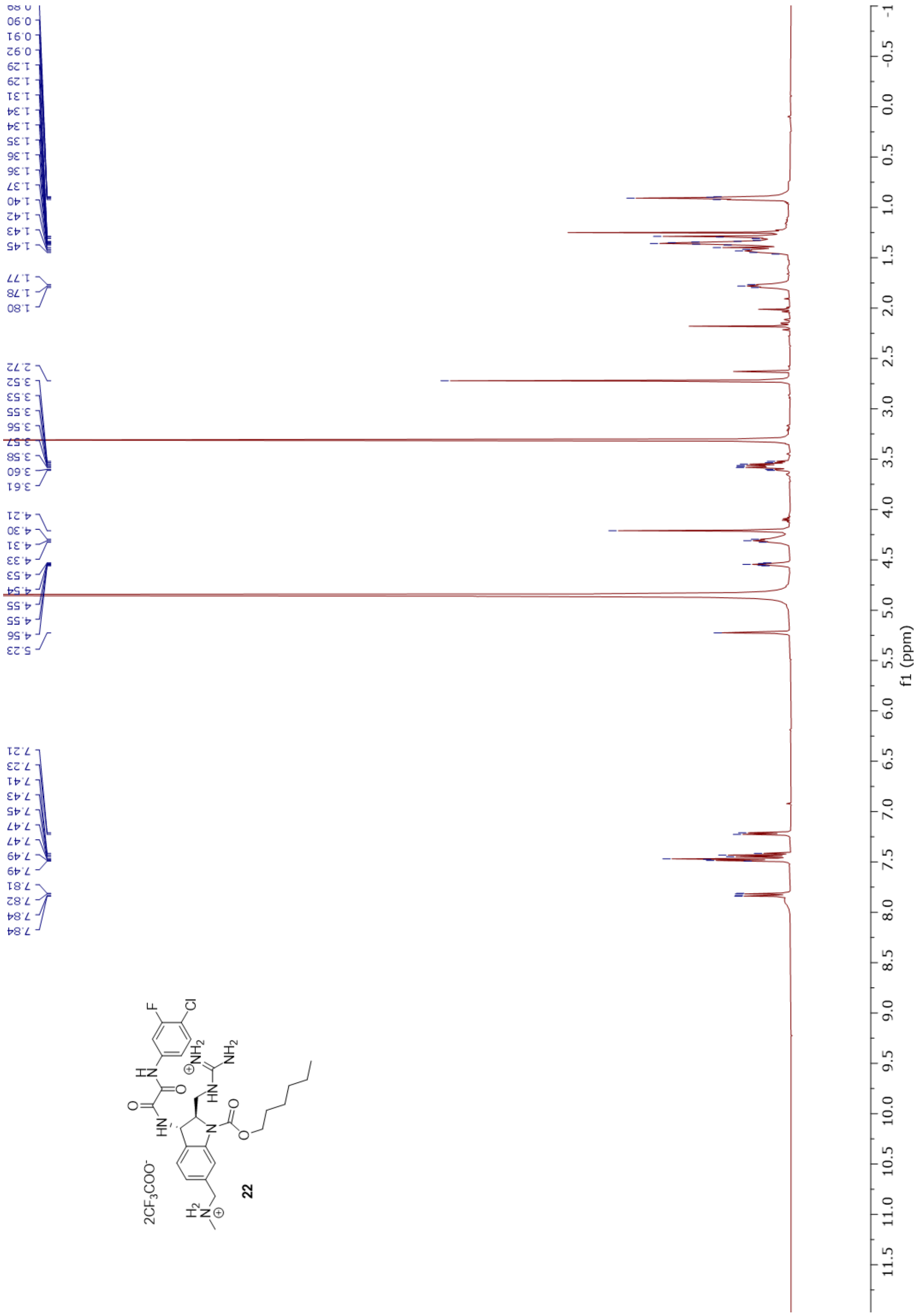
0.89
0.90
0.91
0.92
1.29
1.29
1.31
1.34
1.34
1.35
1.36
1.36
1.37
1.40
1.42
1.43
1.45
1.77
1.78
1.80
2.72
3.52
3.53
3.55
3.56
3.57
3.58
3.60
3.61
4.21
4.30
4.31
4.33
4.53
4.54
4.55
4.55
4.56
5.23
7.21
7.23
7.41
7.43
7.45
7.47
7.47
7.49
7.49
7.81
7.82
7.84
7.84
f1 (ppm)
22

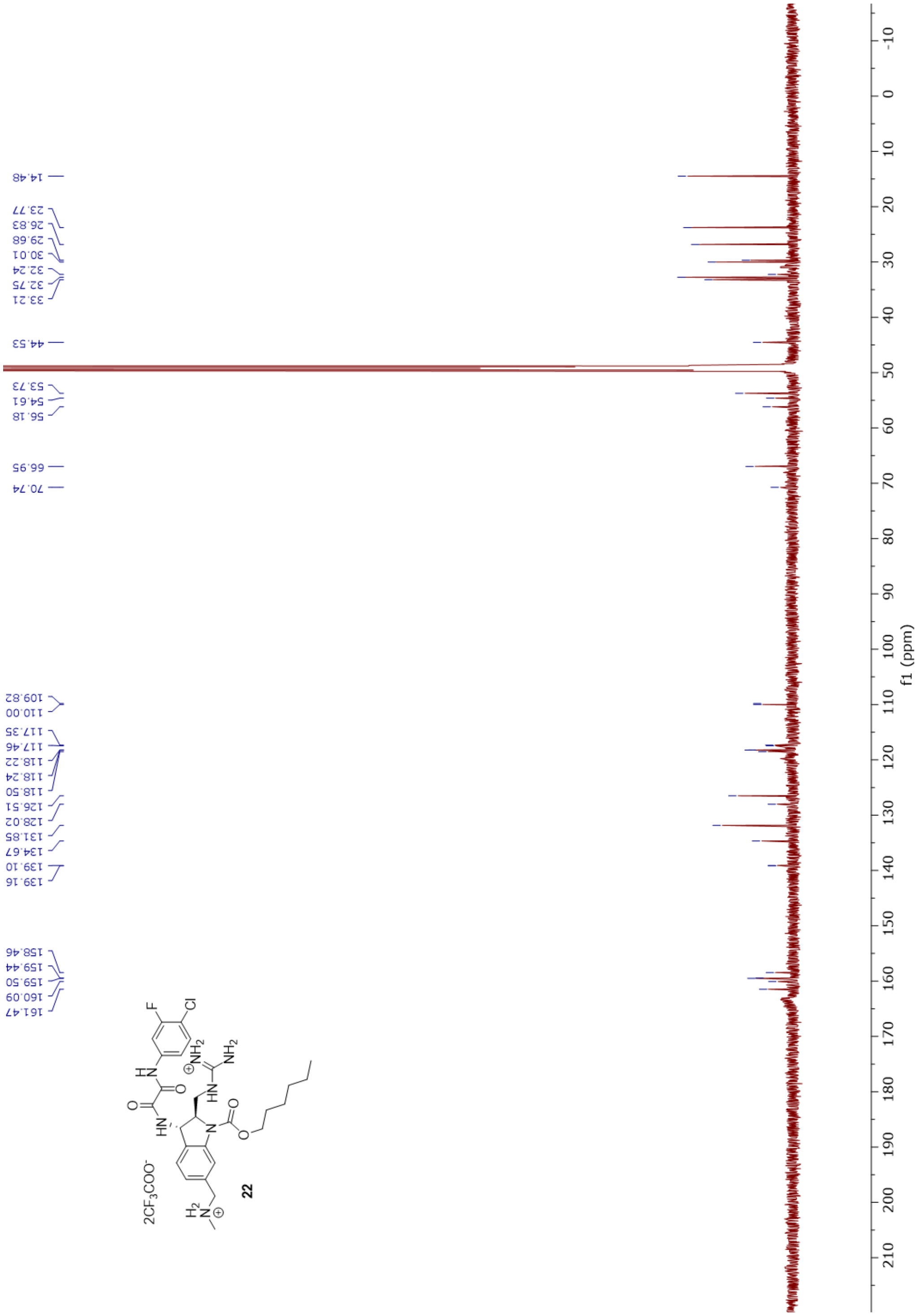
14.48
23.77
26.83
29.68
30.01
32.24
32.75
33.21
44.53
53.73
54.61
56.18
66.95
70.74
109.82
110.00
117.35
117.46
118.22
118.24
118.50
126.51
128.02
131.85
134.67
139.10
139.16
158.46
159.44
159.50
160.09
161.47
f1 (ppm)
2CF3COO-
22

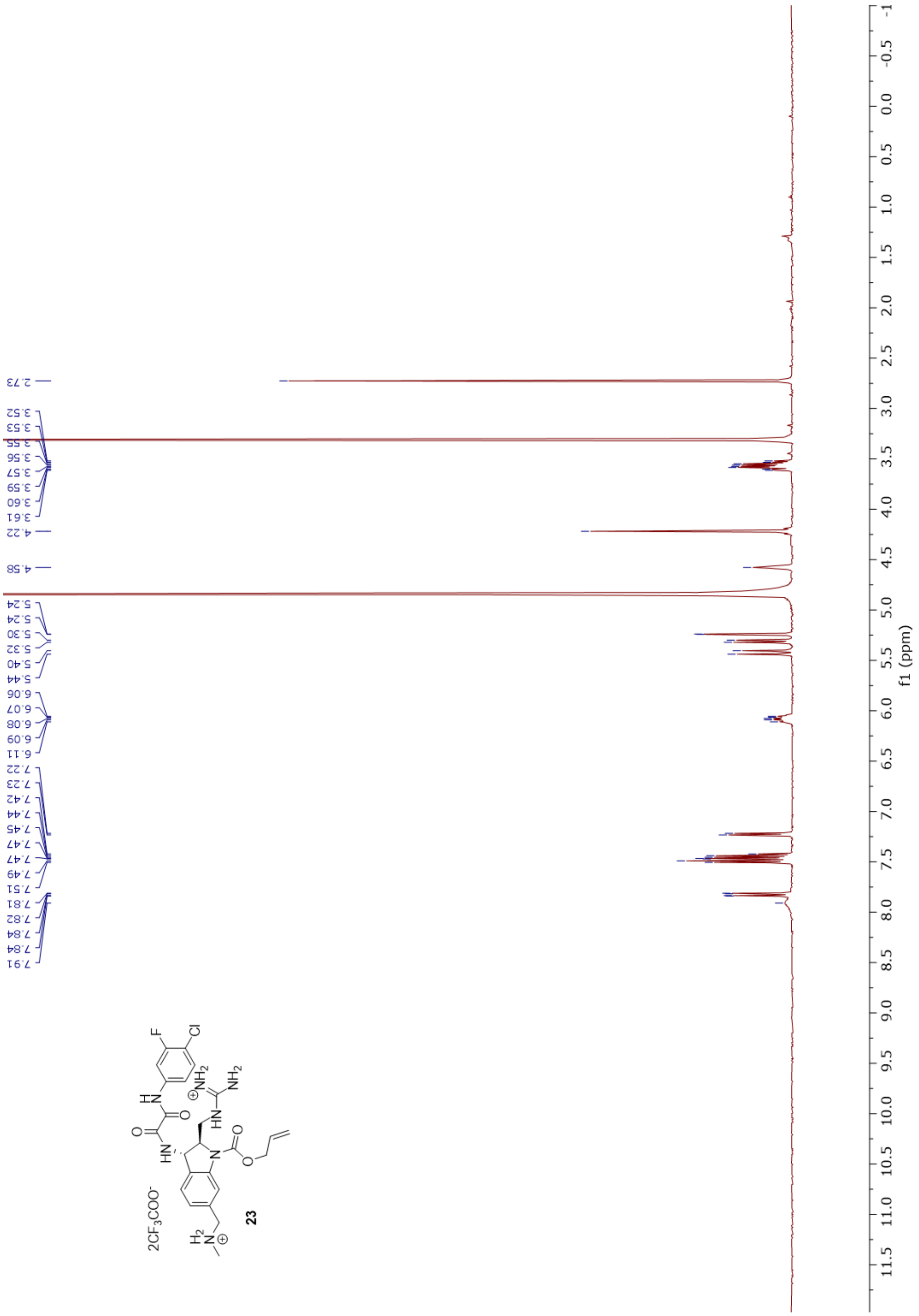
2.73
3.52
3.53
3.55
3.56
3.57
3.59
3.60
3.61
4.22
4.58
5.24
5.24
5.30
5.32
5.40
5.44
6.06
6.07
6.08
6.09
6.11
7.22
7.23
7.42
7.44
7.45
7.47
7.47
7.49
7.51
7.81
7.82
7.84
7.84
7.91
2CF3COO-
23
f1 (ppm)

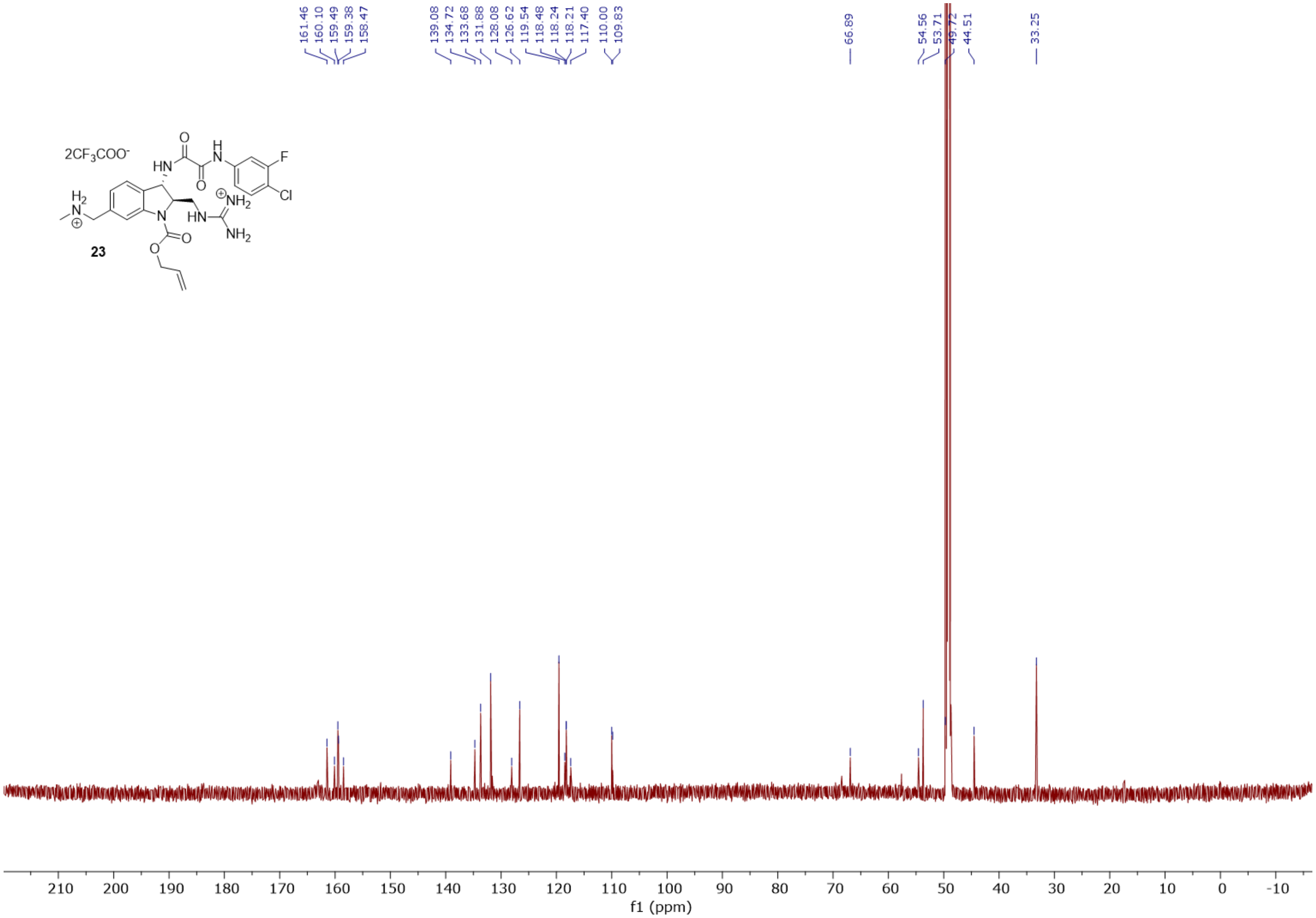
161.46
160.10
159.49
159.38
158.47
139.08
134.72
133.68
131.88
128.08
126.62
119.54
118.48
118.24
118.21
117.40
110.00
109.83
66.89
54.56
53.71
49.72
44.51
33.25
2CF3COO-
HN
O
H
N
F
Cl
O
H2
N
N
HN
NH2
NH2
O
O
23
210
200
190
180
170
160
150
140
130
120
110
100
90
80
70
60
50
40
30
20
10
0
-10
f1 (ppm)

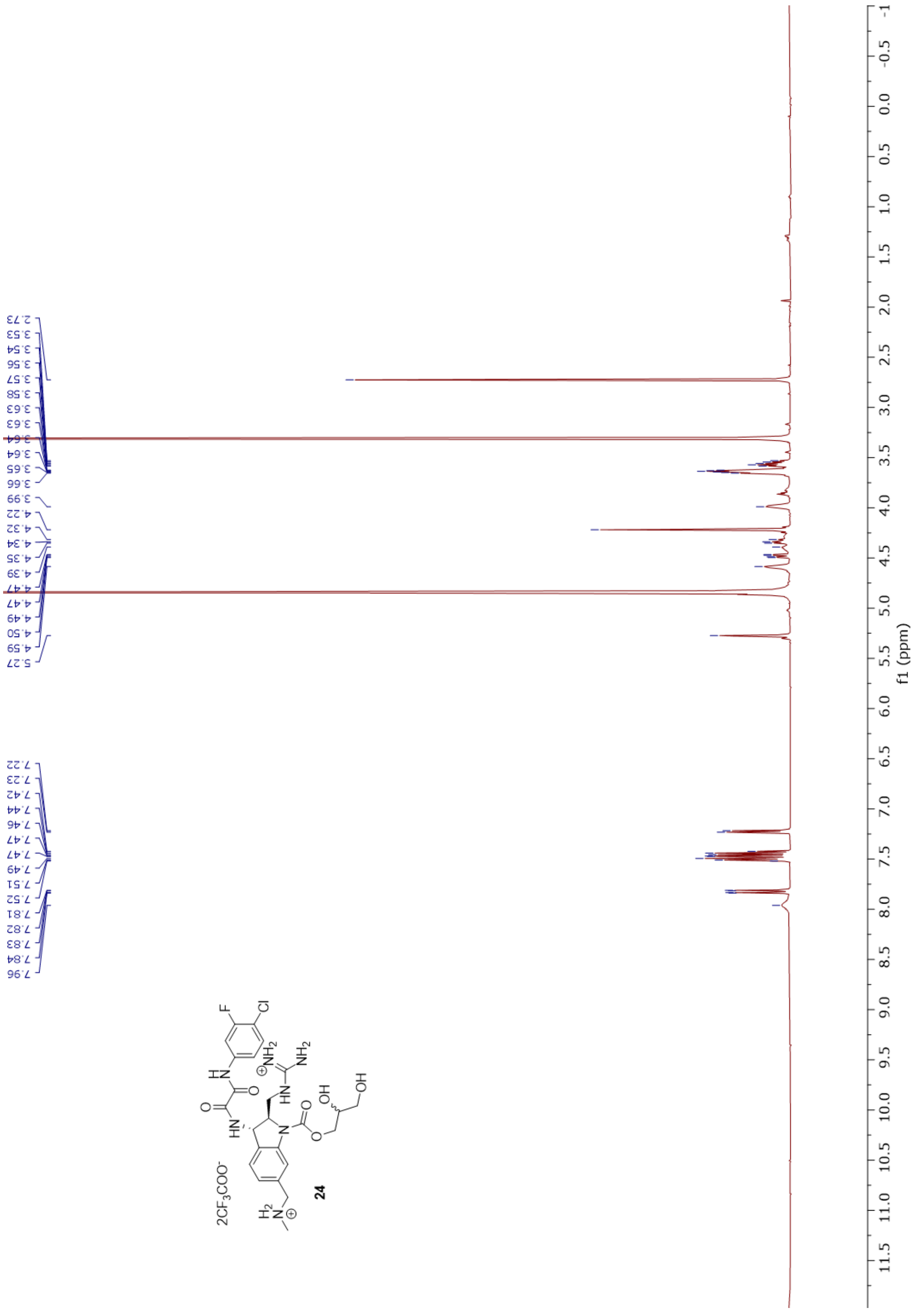
2.73
3.53
3.54
3.56
3.57
3.58
3.63
3.63
3.64
3.64
3.65
3.66
3.99
4.22
4.32
4.34
4.35
4.39
4.47
4.47
4.49
4.50
4.59
5.27
7.22
7.23
7.42
7.44
7.46
7.47
7.47
7.49
7.51
7.52
7.81
7.82
7.83
7.84
7.96
f1 (ppm)
2CF3COO-
24

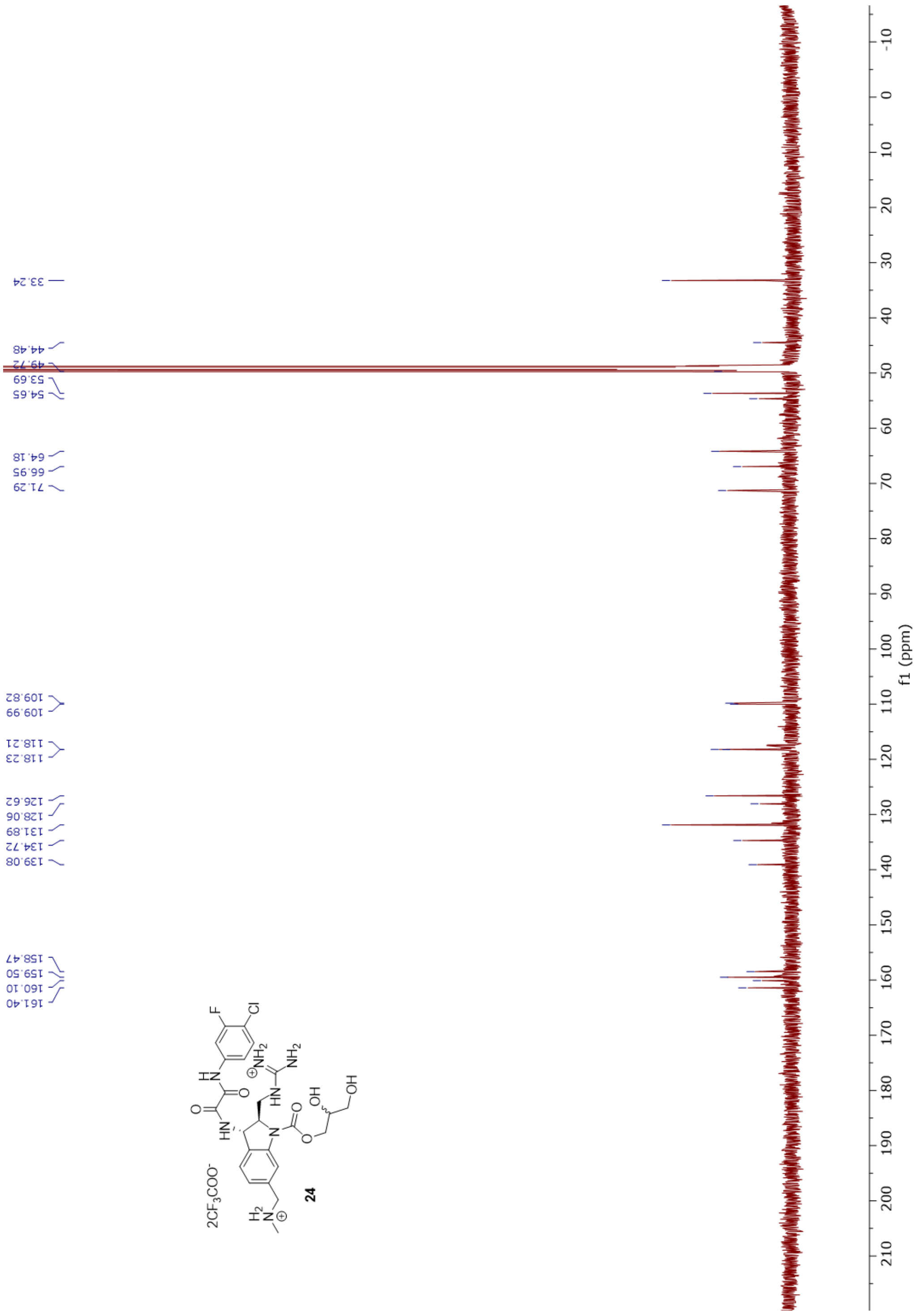
33.24
44.48
49.72
53.69
54.65
64.18
66.95
71.29
109.82
109.99
118.21
118.23
126.62
128.06
131.89
134.72
139.08
158.47
159.50
160.10
161.40
2CF3COO-
24
f1 (ppm)

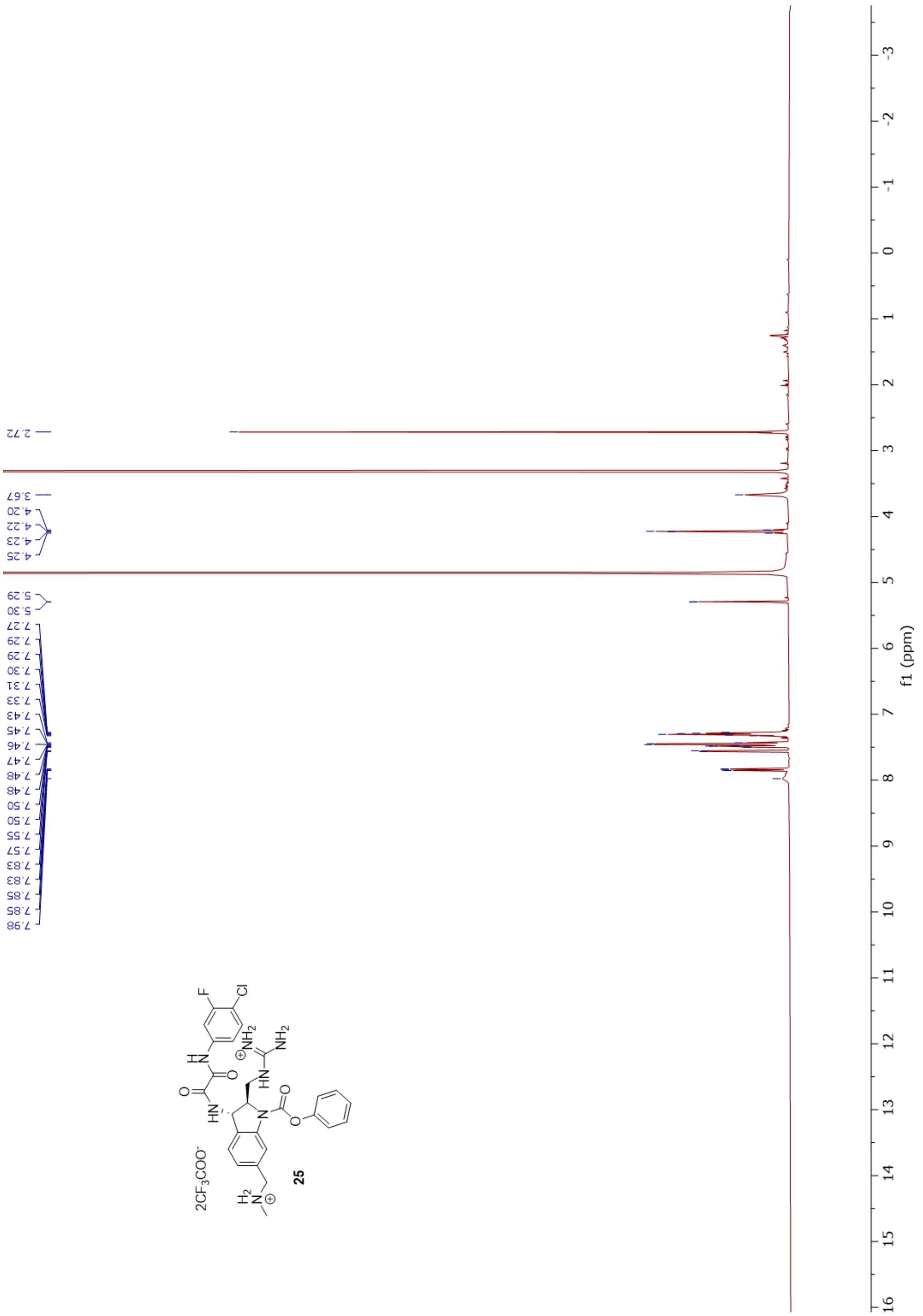
2.72
3.67
4.20
4.22
4.23
4.25
5.29
5.30
7.27
7.29
7.29
7.30
7.31
7.33
7.43
7.45
7.46
7.47
7.48
7.48
7.50
7.50
7.55
7.57
7.83
7.83
7.85
7.85
7.98
2CF3COO-
25
f1 (ppm)

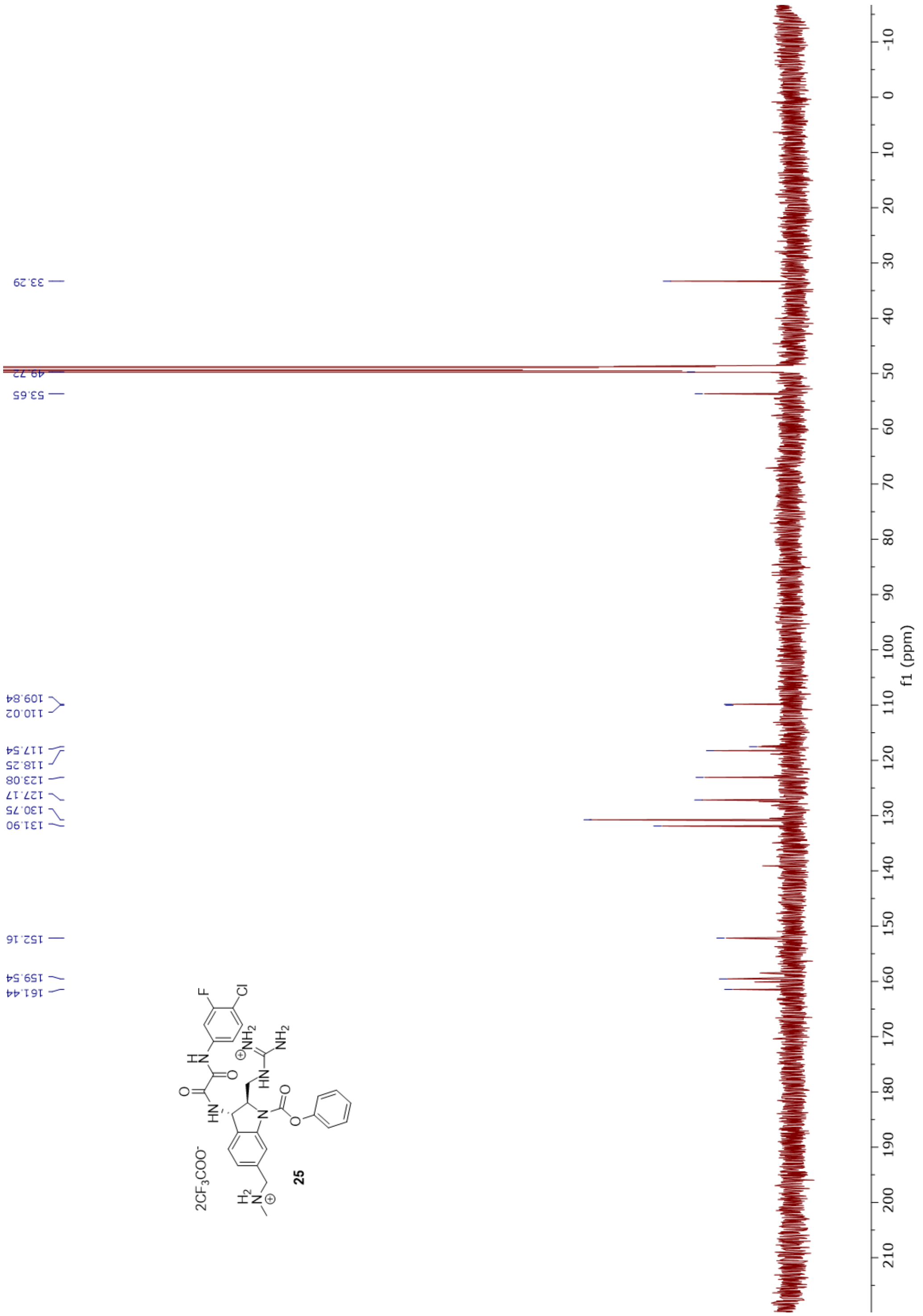

33.29
49.72
53.65
109.84
110.02
117.54
118.25
123.08
127.17
130.75
131.90
152.16
159.54
161.44
2CF3COO-
25
f1 (ppm)

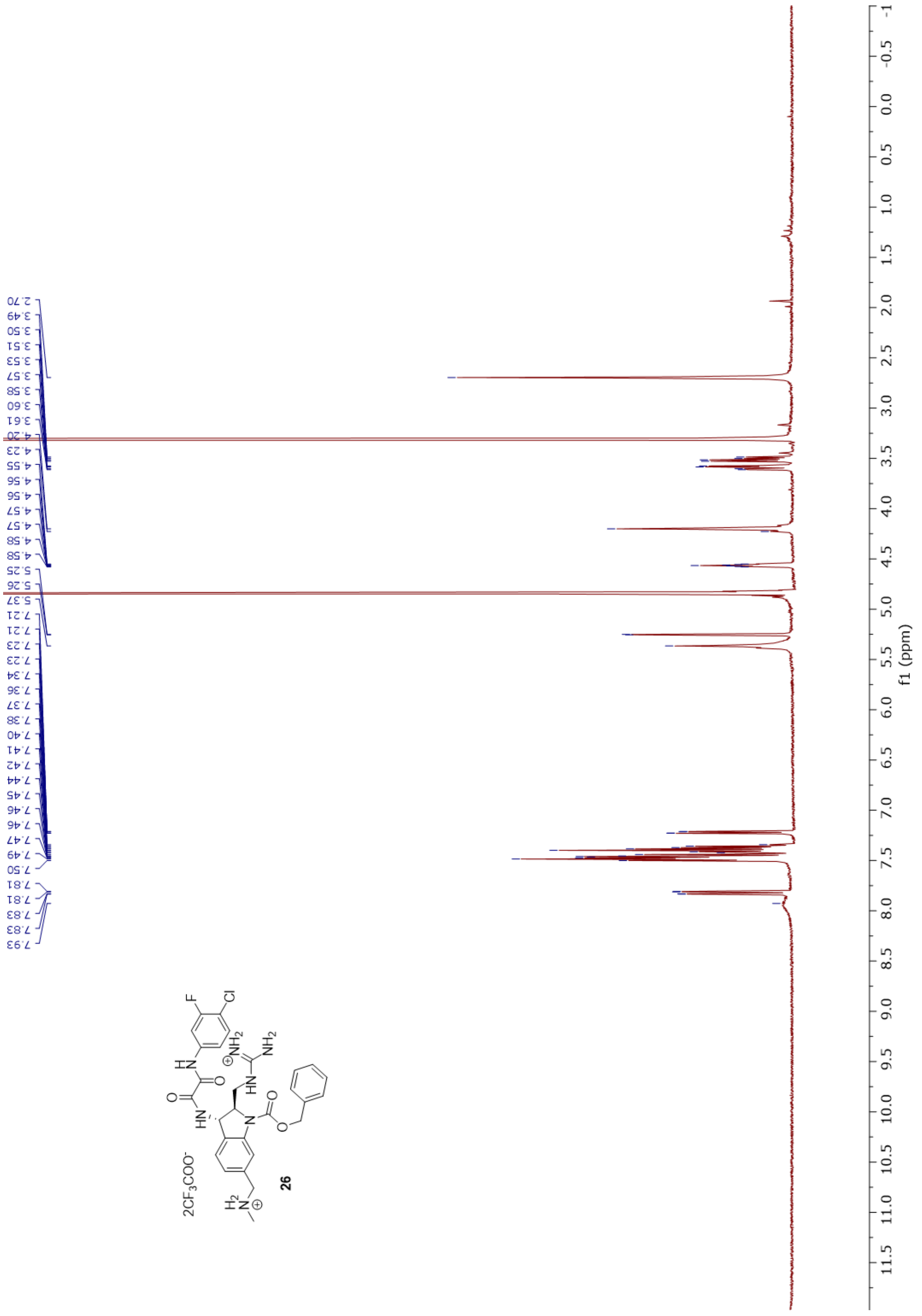
2.70
3.49
3.50
3.51
3.53
3.57
3.58
3.60
3.61
4.20
4.23
4.55
4.56
4.56
4.57
4.57
4.58
4.58
5.25
5.26
5.37
7.21
7.21
7.23
7.23
7.34
7.36
7.37
7.38
7.40
7.41
7.42
7.44
7.45
7.46
7.46
7.47
7.49
7.50
7.81
7.81
7.83
7.83
7.93
2CF3COO-
26
f1 (ppm)

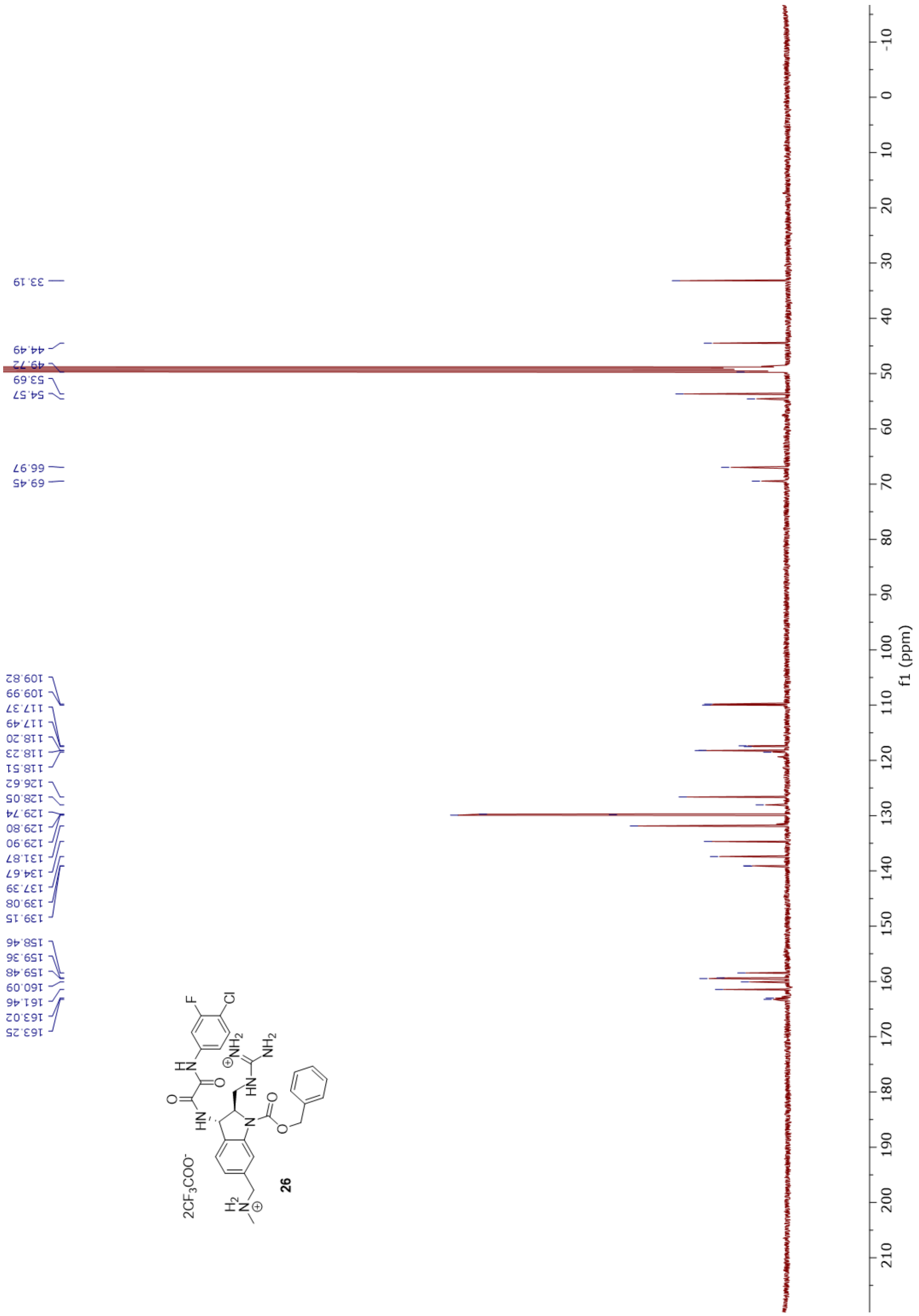
33.19
44.49
49.72
53.69
54.57
66.97
69.45
109.82
109.99
117.37
117.49
118.20
118.23
118.51
126.62
128.05
129.74
129.80
129.90
131.87
134.67
137.39
139.08
139.15
158.46
159.36
159.48
160.09
161.46
163.02
163.25
f1 (ppm)
2CF3COO-
26

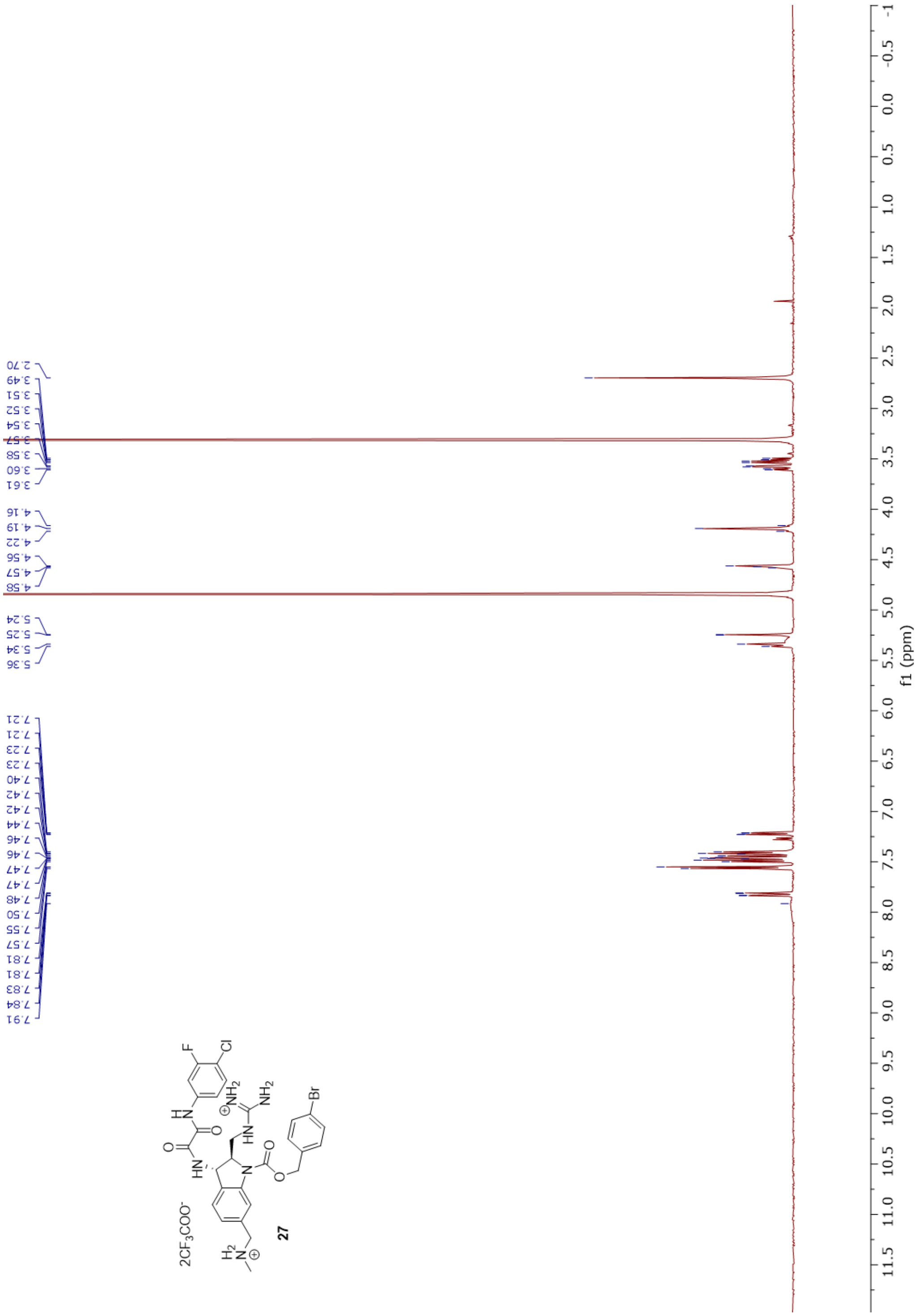

2.70
3.49
3.51
3.52
3.54
3.57
3.58
3.60
3.61
4.16
4.19
4.22
4.56
4.57
4.58
5.24
5.25
5.34
5.36
7.21
7.21
7.23
7.23
7.40
7.42
7.42
7.44
7.46
7.46
7.47
7.47
7.48
7.50
7.55
7.57
7.81
7.81
7.83
7.84
7.91
f1 (ppm)
27

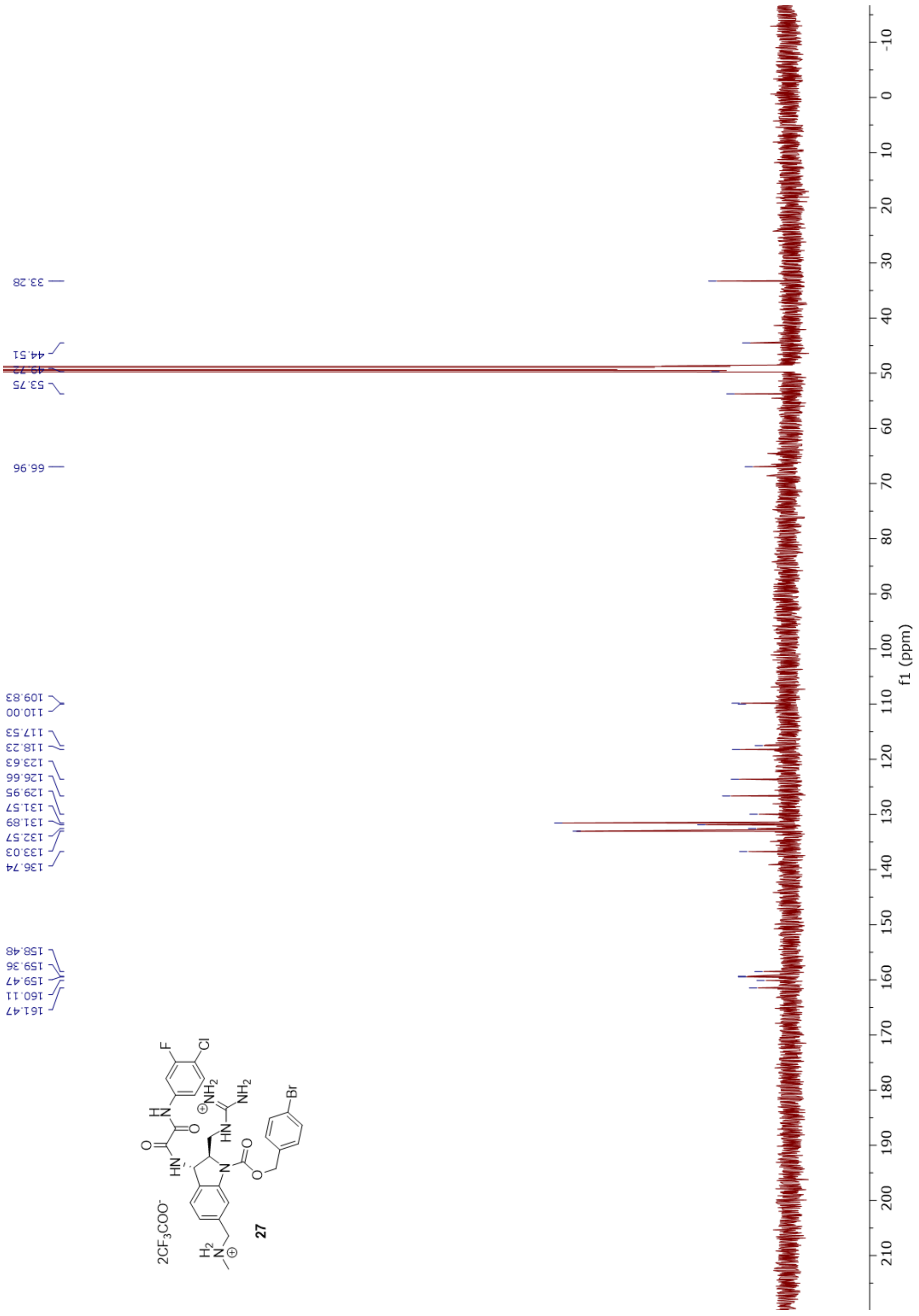
33.28
44.51
53.75
66.96
109.83
110.00
117.53
118.23
123.63
126.66
129.95
131.57
131.89
132.57
133.03
136.74
158.48
159.36
159.47
160.11
161.47
f1 (ppm)
2CF3COO-
27

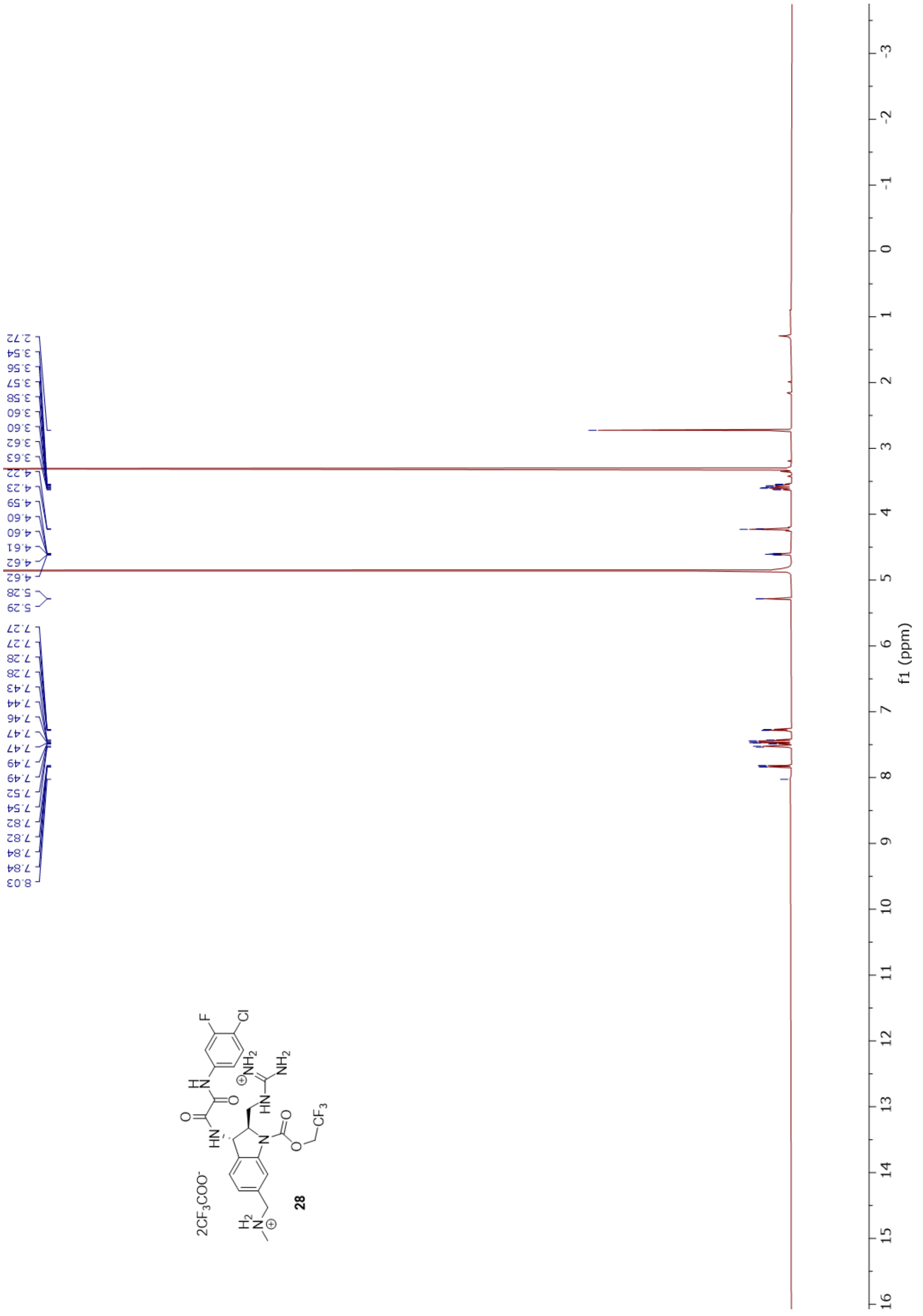
2.72
3.54
3.56
3.57
3.58
3.60
3.60
3.62
3.63
4.22
4.23
4.59
4.60
4.60
4.61
4.62
4.62
5.28
5.29
7.27
7.27
7.28
7.28
7.43
7.44
7.46
7.47
7.47
7.49
7.49
7.52
7.54
7.82
7.82
7.84
7.84
8.03
2CF3COO-
28
f1 (ppm)

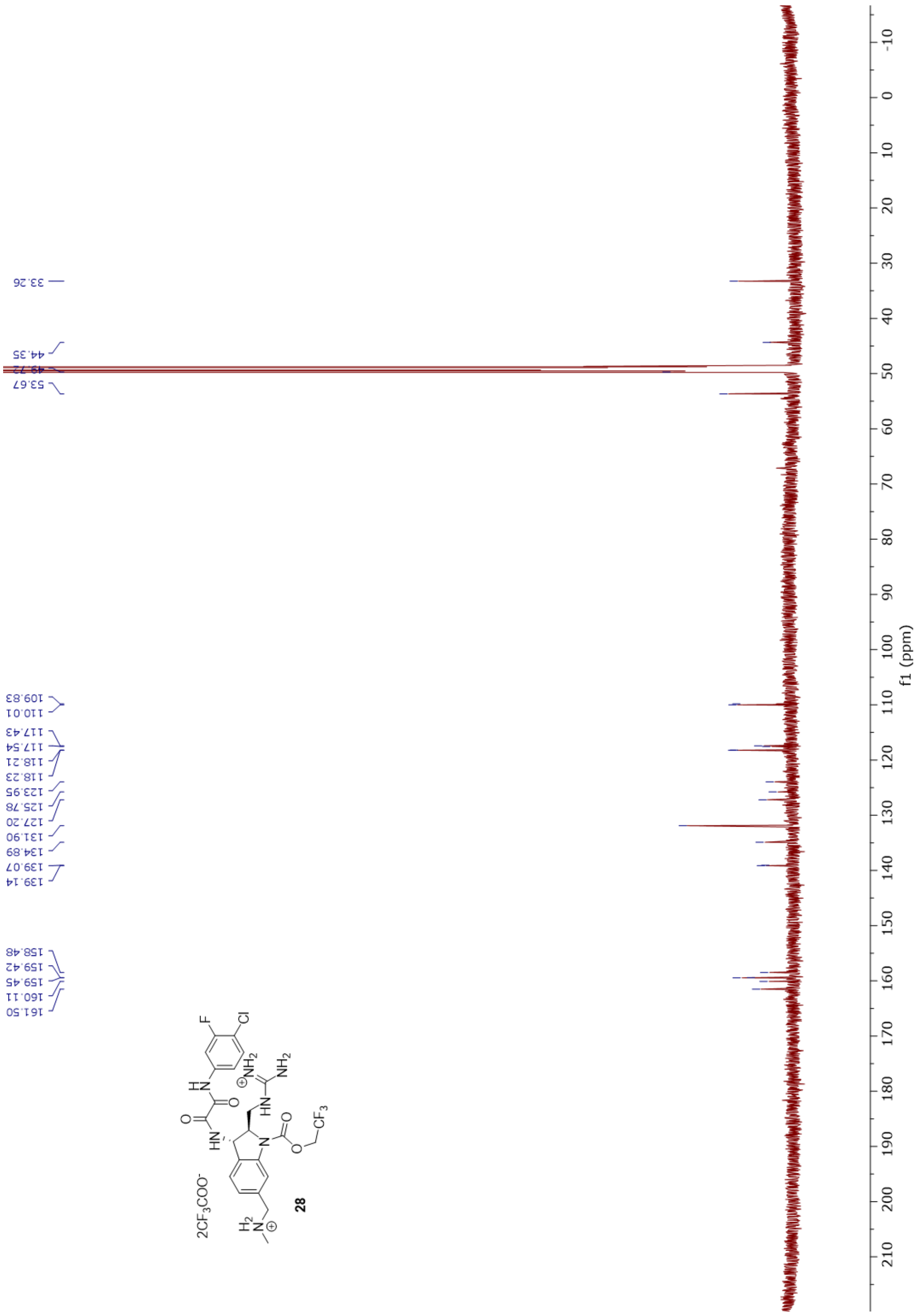
33.26
44.35
49.72
53.67
109.83
110.01
117.43
117.54
118.21
118.23
123.95
125.78
127.20
131.90
134.89
139.07
139.14
158.48
159.42
159.45
160.11
161.50
f1 (ppm)
2CF3COO-

28

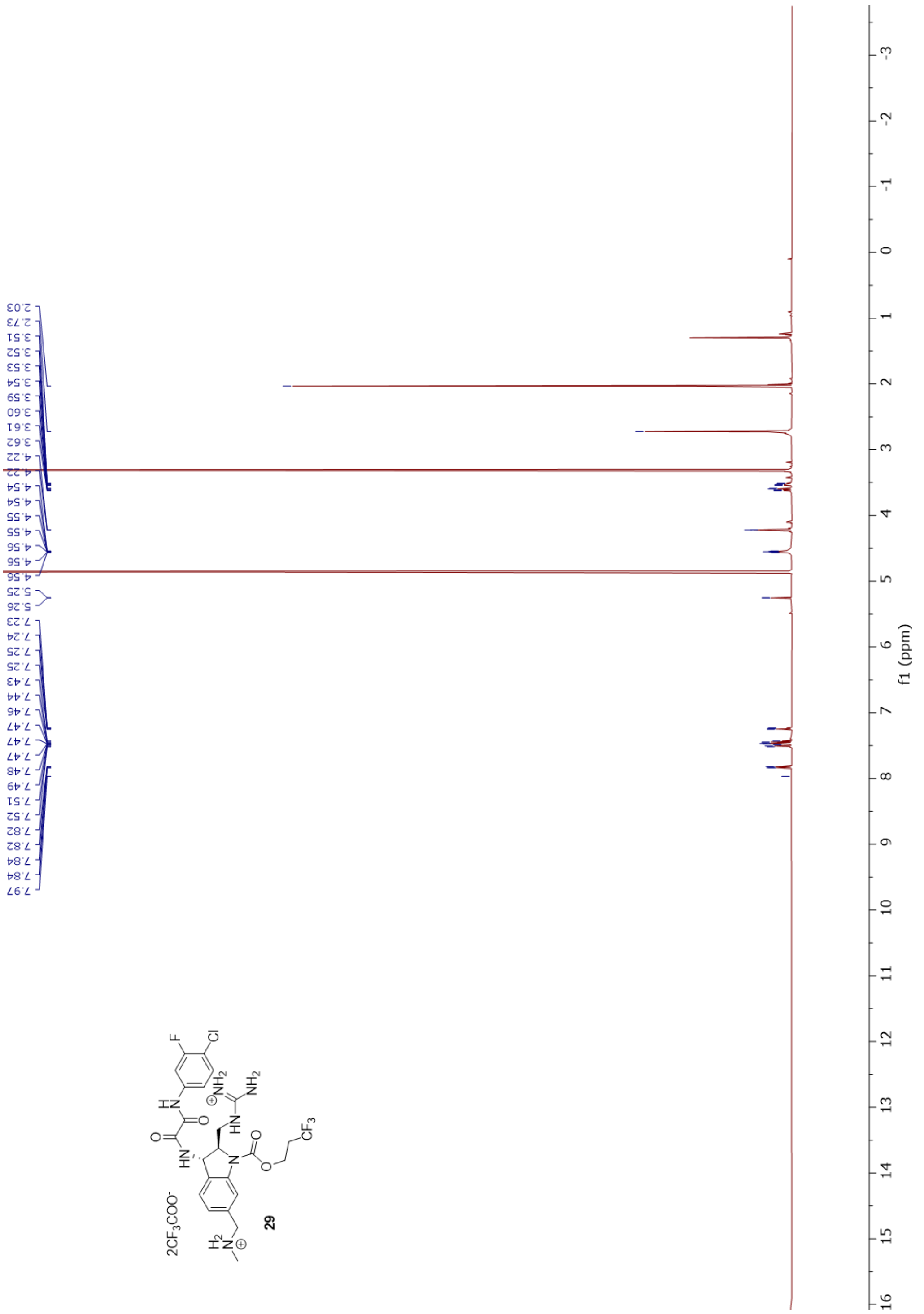
2.03
2.73
3.51
3.52
3.53
3.54
3.59
3.60
3.61
3.62
4.22
4.22
4.54
4.54
4.55
4.55
4.56
4.56
4.56
5.25
5.26
7.23
7.24
7.25
7.25
7.43
7.44
7.46
7.47
7.47
7.47
7.48
7.49
7.51
7.52
7.82
7.82
7.84
7.84
7.97
f1 (ppm)
29

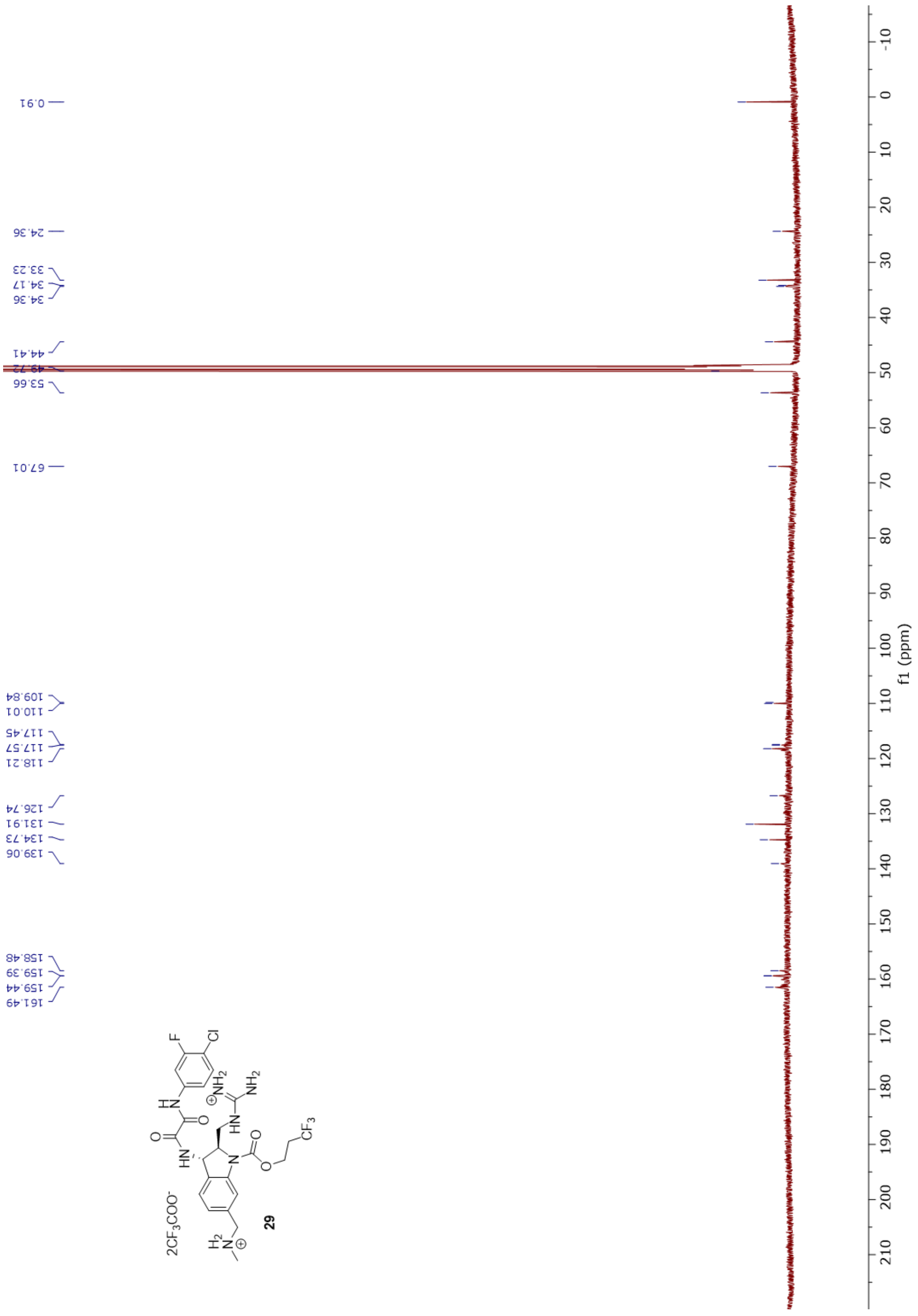
0.91
24.36
33.23
34.17
34.36
44.41
49.72
53.66
67.01
109.84
110.01
117.45
117.57
118.21
126.74
131.91
134.73
139.06
158.48
159.39
159.44
161.49
f1 (ppm)
2CF3COO-
29

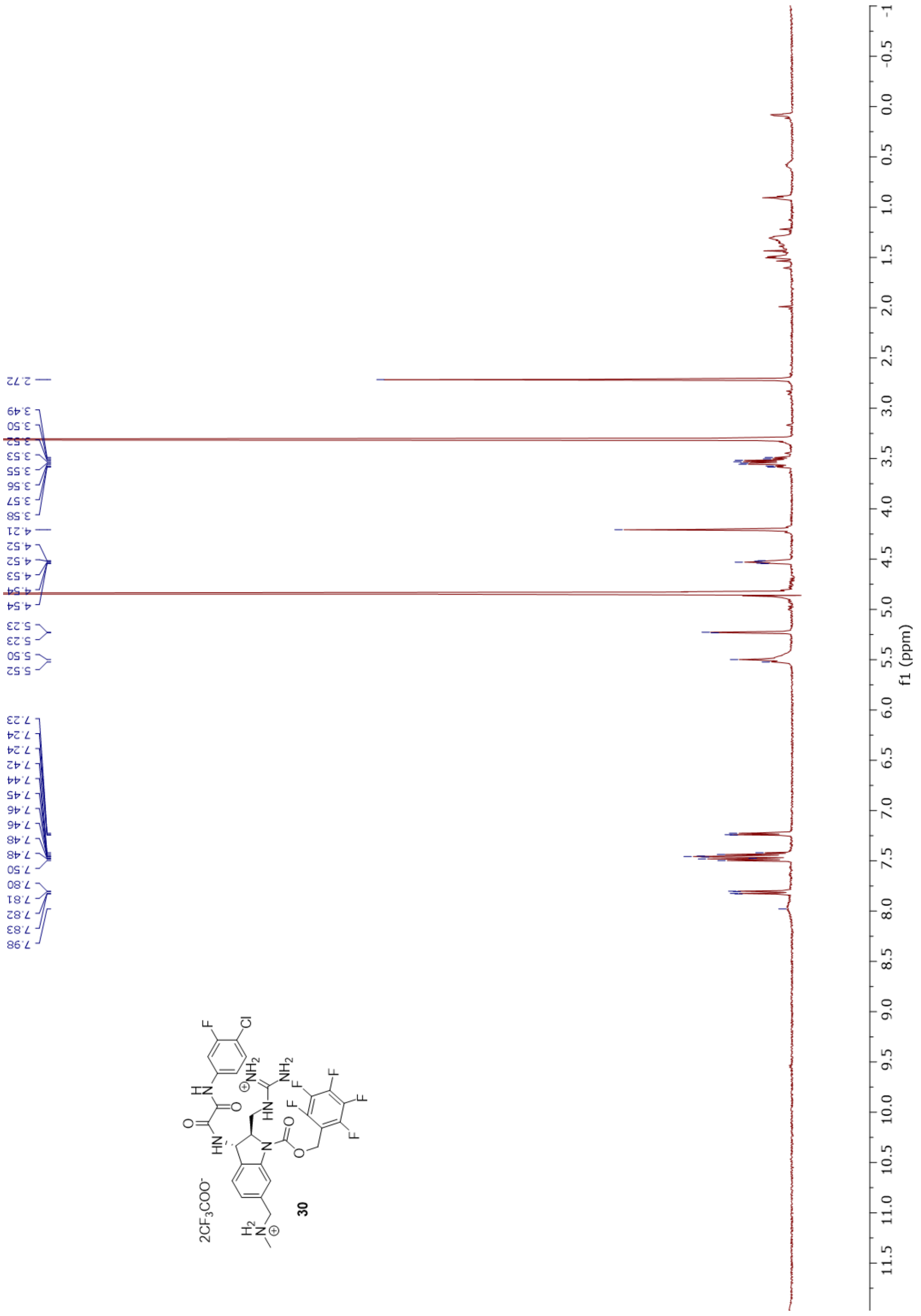

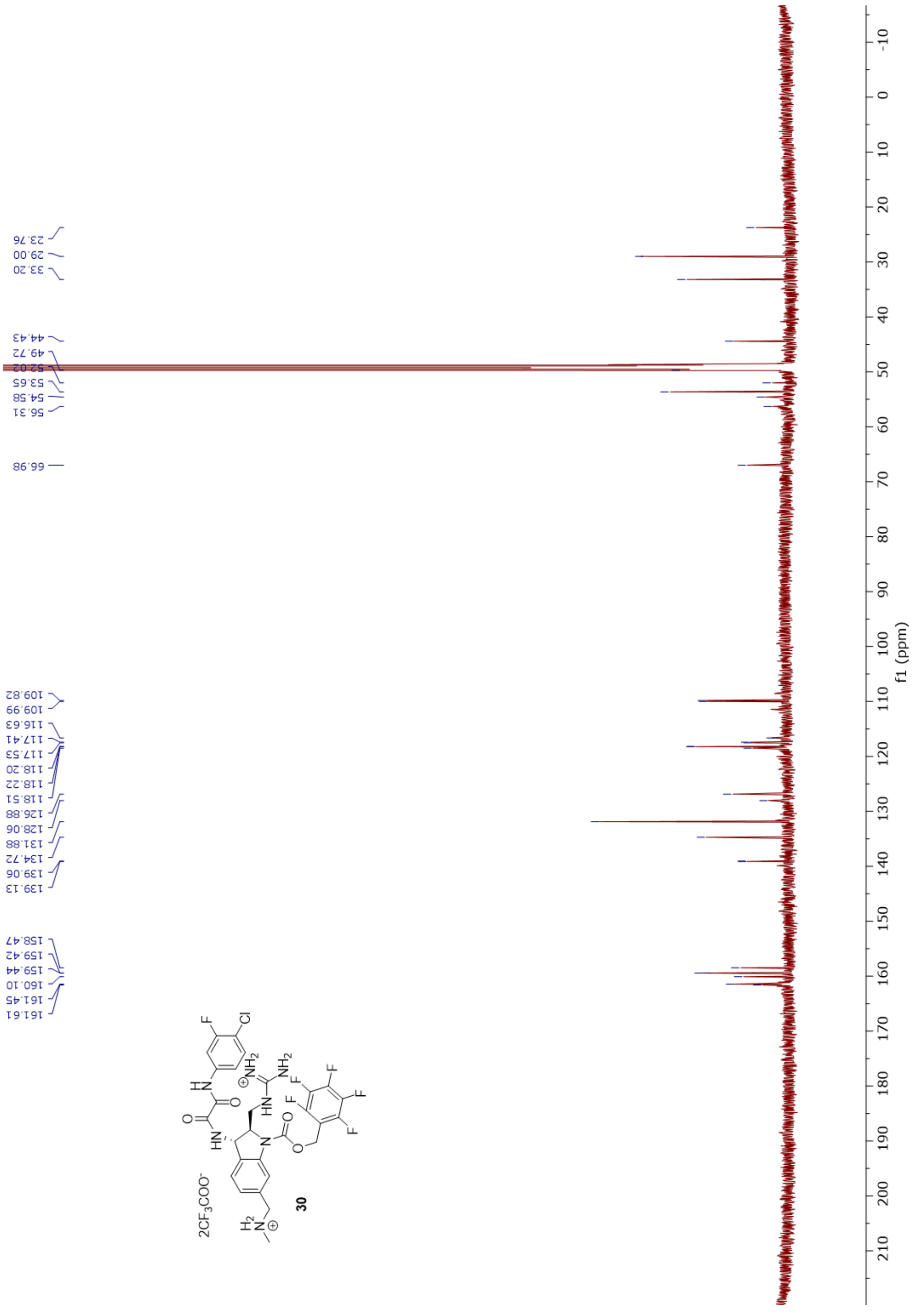

23.76
29.00
33.20
44.43
49.72
52.02
53.65
54.58
56.31
66.98
109.82
109.99
116.63
117.41
117.53
118.20
118.22
118.51
126.88
128.06
131.88
134.72
139.06
139.13
158.47
159.42
159.44
160.10
161.45
161.61
f1 (ppm)
30
